# Expansion and revision of the genus Xanthobacter and proposal of Roseixanthobacter gen. nov.

**DOI:** 10.1101/2024.07.11.603134

**Authors:** Maximillian P. M. Soltysiak, Amogh P. Jalihal, Caroline E. Christophersen, Audrey L. H. Ory, Andrew D. Lee, Jessica Boulton, Michael Springer

**Author notes:** **REPOSITORIES:**. The GenBank accession numbers for the 16S rRNA gene sequences and genome sequences are: *X. agilis* MA37 (PP328848; JBAFWL000000000), *X. agilis* MA40 (PP328849; JBAFWK000000000), *X. agilis* SA35^T^ (PP328869; JBAFWJ000000000), *X. albus* V0B-10 (PP328879; JBAFWC000000000), *X. albus* V0C-6^T^ (PP328881; JBAFWE000000000), *X. albus* V13C-5 (PP328883; JBAFWD000000000), *X. albus* V2C-4 (PP328888; JBAFWG000000000), *X. albus* V2C-8 (PP328889; JBAFWF000000000), *X. aminoxidans* 14a^T^ (PP328862; JBAFUV000000000), *X. aminoxidans* CB3 (PP328858; JBAFUS000000000), *X. aminoxidans* CB5 (PP328859; JBAFUR000000000), *X. aminoxidans* V7C-9 (PP328902; JBAFUT000000000), *X. aminoxidans* V8C-8 (PP328904; JBAFUU000000000), *X. autotrophicus* 124X (PP328873; JBAFVO000000000), *X. autotrophicus* 19/-/x (PP328864; JBAFVR000000000), *X. autotrophicus* 7C^T^ (PP328871; JBAFVK000000000), *X. autotrophicus* 7C SF (PP328866; JBAFVJ000000000), *X. autotrophicus* 7d (PP328865; JBAFVI000000000), *X. autotrophicus* CCUG 44692 (PP328853; JBAFVL000000000), *X. autotrophicus* GZ29 (PP328856; JBAFVQ000000000), *X. autotrophicus* JW33 (PP328863; JBAFVN000000000), *X. autotrophicus* NCIMB 11171 (PP328877; JBAFVM000000000), *X. autotrophicus* V0C-4 (PP328880; JBAFVP000000000), *X. cornucopiae* V4C-4^T^ (PP328896; JBAFWM000000000), *X. flavus* 301^T^ (PP328868; JBAFUM000000000), *X. flavus* R-10 (PP328861; JBAFUK000000000), *X. flavus* SS-20 (PP328875; JBAFUQ000000000), *X. flavus* SS-22 (PP328876; JBAFUP000000000), *X. flavus* V2C-3 (PP328887; JBAFUN000000000), *X. flavus* V4C-10 (PP328895; JBAFUJ000000000), *X. flavus* V4C-7 (PP328897; JBAFUL000000000), *X. flavus* V4C-9 (PP328899; JBAFUO000000000), *X. lutulentifluminis* V3C-3^T^ (PP328893; JBAFVU000000000), *X. nonsaccharivorans* 14g^T^ (PP328870; JBAFVV000000000), *X. oligotrophicus* 23A (PP328874; JBAFVH000000000), *X. pseudotagetidis* KA^T^ (PP328855; JBAFVY000000000), *X. sediminis* V3B-7B (PP328891; JBAFWI000000000), *X. sediminis* V8C-5^T^ (PP328903; JBAFWH000000000), *X. tagetidis* A2 (PP328850; JBAFVX000000000), *X. tagetidis* TagT2C^T^ (PP328854; JBAFVW000000000), *X. toluenivorans* T101^T^ (PP328851; JBAFVT000000000), *X. toluenivorans* T102 (PP328852; JBAFVS000000000), *X. variabilis* V13B-9B (PP328882; JBAFVZ000000000), *X. variabilis* V13C-7B (PP328884; JBAFWA000000000), *X. variabilis* V4C-8^T^ (PP328898; JBAFWB000000000), *X. versatilis* CB6 (PP328860; JBAFVA000000000), X*. versatilis* V1C-2 (PP328885; JBAFVD000000000), *X. versatilis* V1C-6B (PP328886; JBAFVC000000000), *X. versatilis* V2C-9 (PP328890; JBAFVB000000000), *X. versatilis* V3C-2 (PP328892; JBAFVG000000000), *X. versatilis* V3C-4 (PP328894; JBAFVF000000000), *X. versatilis* V7C-1B (PP328900; JBAFVE000000000), *X. versatilis* V7C-4 (PP328901; JBAFUZ000000000), *X. wiegelii* CB2 (PP328857; JBAFUW000000000), *X. wiegelii* NCIMB 11399 (PP328878; JBAFUX000000000), *X. wiegelii* RH 10^T^ (PP328872; JBAFUY000000000), *R. finlandensis* VTT E-85241^T^ (PP328909; JBAFWO000000000), *R. glucoisosaccharinivorans* VTT E-85242^T^ (PP328910; JBAFWN000000000), *R. liquoris* VTT E-85238^T^ (PP328906; JBAFWP000000000), *R. pseudopolyaromaticivorans* VTT E-85237 (PP328905; JBAFWS000000000), *R. pseudopolyaromaticivorans* VTT E-85239 (PP328907; JBAFWR000000000), *R. pseudopolyaromaticivorans* VTT E-85240^T^ (PP328908; JBAFWQ000000000), and *R. psychrophilus* W30^T^ (PP328867; JBAFWT000000000).

## Abstract

The nitrogen-fixing, chemolithoautotrophic genus Xanthobacter is found worldwide across numerous diverse environments and is an important member of many ecosystems. These species serve as model systems for their metabolic properties in academic settings and have industrial applications in bioremediation and sustainable protein, food, and fertilizer production. Despite their abundance, interest, and importance, the majority of Xanthobacter strains are without a genome sequence, and only 8 validly published species are known to date. To expand our understanding of the diversity and evolutionary history of the genus, we sequenced the genomes of 37 repository strains and 26 novel environmental strains we isolated. After performing comparative phylogenomic analyses, we expand and revise the genus Xanthobacter and propose the novel genus Roseixanthobacter gen. nov. For the Xanthobacter, we describe 10 novel species, bringing the total to 18: *Xanthobacter agilis*, *Xanthobacter albus* sp. nov., *Xanthobacter aminoxidans*, *Xanthobacter autotrophicus*, *Xanthobacter cornucopiae* sp. nov., *Xanthobacter dioxanivorans*, *Xanthobacter flavus*, *Xanthobacter lutulentifluminis* sp. nov., *Xanthobacter nonsaccharivorans* sp. nov., *Xanthobacter oligotrophicus*, *Xanthobacter pseudotagetidis* sp. nov., *Xanthobacter sediminis* sp. nov., *Xanthobacter tagetidis*, *Xanthobacter toluenivorans* sp. nov., *Xanthobacter variabilis* sp. nov., *Xanthobacter versatilis* sp. nov., *Xanthobacter viscosus*, and *Xanthobacter wiegelii* sp. nov. For the Roseixanthobacter gen. nov., we describe 5 novel species formerly classified as Xanthobacter: *Roseixanthobacter finlandensis* sp. nov., *Roseixanthobacter glucoisosaccharinivorans* sp. nov., *Roseixanthobacter liquoris* sp. nov., *Roseixanthobacter pseudopolyaromaticivorans* sp. nov., and *Roseixanthobacter psychrophilus* sp. nov. We characterized the phenotypic properties of these type strains, including temperature, salinity and pH ranges, carbon substrate utilization, motility, antibiotic susceptibility, slime production, autotrophic growth, and enzymatic activities. We discovered a more diverse range of phenotypes across the genus Xanthobacter than previously known and elucidated the evolutionary history within the genus. These findings and genome sequences will help further the application of Xanthobacter biology in academic, industrial, and environmental settings and provide additional insight into the unique biological properties that make these species attractive for such applications.

## INTRODUCTION

The genus Xanthobacter is known for its hydrogen-oxidizing, nitrogen-fixing, chemolithoautotrophic growth capabilities and yellow pigmentation from zeaxanthin dirhamnoside production (1–3). These strains are found worldwide across numerous diverse environments and are important members of ecosystems, including soil (2,4–6), wastewater sludge (7–10), sediment (3), water (11), root systems (12,13), and more. In ecosystems, Xanthobacter are fixers of carbon and nitrogen (3) and solubilize phosphate (14). Xanthobacter can detoxify environments and free up growth substrates by removing difficult moieties such as halogens from organic substrates (15–18). Furthermore, some species promote plant growth and form key plant-microbe and microbe-microbe interactions in ecosystems and microbiomes (12,13,19).

In academic and laboratory settings, Xanthobacter are model organisms for studying nitrogen fixation (20,21) and chemolithoautotrophy (22–24). They contain several novel and unique enzymes, including the first known alkane dehalogenases (15,16), and possess alkene monooxygenases with diverse substrate ranges (25–28). With these unique activities, Xanthobacter strains are being explored for bioremediation given that some strains can degrade difficult substrates of interest, such as haloalkanes (15,16), haloalkanoic acids (15,29), alkenes and chlorinated alkenes (18,25–28,30), toluene (31), cyclohexane (32), styrene (17), and acetone (33,34). Industrially, Xanthobacter strains have been recently used for the sustainable production of products ranging from protein (35–39) and foodstuffs (40) to fertilizer (19). Uniquely, these bacteria can grow in chambers directly in contact with water-splitting machinery needed to generate hydrogen and the chemolithoautotrophic conditions they need for sustainable growth (19,40).

Despite the abundance, importance, and academic and industrial interest in Xanthobacter, the vast majority of repository strains lack a genome sequence and taxonomic validation. To date, only eight species have been validly published (1,5,7,9–12), although more have been described (39,41–43), meaning we likely lack a great deal of knowledge on the diversity and evolutionary history of the genus. Previous taxonomic classification methods, such as 16S rRNA analyses, are notoriously inadequate for resolving genus and species boundaries between and within the genus Xanthobacter and the highly related genera Azorhizobium and Aquabacter due to high sequence similarity (44,45). Thus, there is a clear and immediate need for genome-level analyses that are not plagued by these issues.

In this study, we sought to expand and validate our understanding of the genus Xanthobacter and its diversity. We sequenced nearly every Xanthobacter repository strain we could obtain and isolated novel Xanthobacter from hydrogen-oxidizing, nitrogen-fixing, chemolithoautotrophic communities from environmental samples. After phylogenomic comparisons and phenotypic characterization, we have significantly expanded and revised the genus Xanthobacter, bringing the total known species from 8 to 18. Additionally, we propose a novel genus, Roseixanthobacter gen. nov., comprising of 5 total species.

## MATERIALS AND METHODS

### Bacterial strains

Strains previously identified and listed as Xanthobacter were acquired from culture collections worldwide (Table 1). We also included additional strains identified as Xanthobacter that we isolated from nitrogen-fixing, hydrogen-oxidizing, chemolithoautotrophic communities grown from environmental samples (Table 1). These environmental samples were taken from various sources in Boston, Massachusetts, USA from soil, water, mud, and sediment. Samples were topped up to 50 mL volumes with inorganic minimal media (IMM), and a 1/10 dilution was used to start the community enrichment cultures. The diluted environmental samples were grown at 30°C and max spinning in a 30 mL volume of IMM using the eVOLVER system (46). A mixed gas supply (∼3% H_2_, ∼2% O_2_, ∼10% CO_2_, ∼85% N_2_) was bubbled through the cultures using a sparger to sustain nitrogen-fixing hydrogen-oxidizing chemolithoautotrophic growth.

**Table 1.** List of bacterial strains sequenced and used in biological work for the study. Strain identifiers in bold are the accessions for the source repository used in this study. Abbreviations are: ATCC – American Type Culture Collection; CCUG – Culture Collection University of Gothenburg; DSMZ – Deutsche Sammlung von Mikroorganismen und Zellkulturen GmbH/German Collection of Microorganisms and Cell Cultures GmbH; NBRC – National Institute of Technology and Evaluation (NITE) Biological Resource Center; NCIMB – National Collection of Industrial, Food and Marine Bacteria; VTT – Valtion Teknillinen Tutkimuskeskus Culture Collection. Strains indicated with an asterisk were not newly sequenced in this study but were used in biological and bioinformatic work. A full list of all strains used for bioinformatic work is provided in Supplementary Table 2.

| Species designation | Prior species designation | Other strain identifiers | Source |
| --- | --- | --- | --- |
| *Xanthobacter agilis* MA37 | *Xanthobacter agilis* MA37 | **ATCC 43848**; NEU 2067 | ATCC |
| *Xanthobacter agilis* MA40 | *Xanthobacter agilis* MA40 | **ATCC 43849**; NEU 2069 | ATCC |
| *Xanthobacter agilis* SA35<sup>T</sup> | *Xanthobacter agilis* SA35<sup>T</sup> | ATCC 43847; **DSM 3770**;<br>LMG 16336; VKM B-2105;<br>LMG 7994; NCIMB 12683;<br>NEU 2015; NCAIM<br>B.01949 | DSMZ |
| *Xanthobacter albus* V0B-10 |  | DSM | This study;<br>Surface soil<br>sample, Boston,<br>MA, USA |
| *Xanthobacter albus* V0C-6<sup>T</sup> |  | ATCC; DSM | This study;<br>Surface soil<br>sample, Boston,<br>MA, USA |
| *Xanthobacter albus* V13C-5 |  | DSM | This study; Silt<br>and sediment<br>sample, Muddy<br>River, Boston, MA,<br>USA |
| *Xanthobacter albus* V2C-4 |  | DSM | This study; Water<br>sample, Muddy<br>River, Boston, MA |
| *Xanthobacter albus* V2C-8 |  | DSM | This study; Water<br>sample, Muddy<br>River, Boston, MA |
| *Xanthobacter aminoxidans* 14a<sup>T</sup> | *Xanthobacter aminoxidans* 14a<sup>T</sup> | ATCC BAA-299; CIP<br>108461; **DSM 15009**;<br>KCTC 12307; VKM B-2254 | DSMZ |
| *Xanthobacter aminoxidans* CB3 | *Xanthobacter* sp. CB3 | **DSM 14518** | DSMZ |
| *Xanthobacter aminoxidans* CB5 | *Xanthobacter* sp. CB5 | **DSM 14519** | DSMZ |
| *Xanthobacter aminoxidans* V7C-9 |  | DSM | This study;<br>Sediment sample,<br>Muddy River,<br>Boston, MA, USA |
| *Xanthobacter aminoxidans* V8C-8 |  | DSM | This study;<br>Sediment sample,<br>Muddy River,<br>Boston, MA, USA |
| *Xanthobacter autotrophicus* 124X | *Xanthobacter* sp. 124X | ATCC 49450; **DSM 6696** | DSMZ |
| *Xanthobacter autotrophicus* 19/-/x | *Xanthobacter autotrophicus* 19/-/x | **DSM 2009** | DSMZ |
| *Xanthobacter autotrophicus* 7C<sup>T</sup> | *Xanthobacter autotrophicus* 7C<sup>T</sup> | ATCC 35674; BCRC 12235; CCRC 12235; CIP 105431; **DSM 432**; IAM 12579; IAM 12636; JCM 1202; LMG 7043; NBRC 102463; NCAIM B0.1945; NCIB 10809; NCIMB 10809; NRRL B-14836 | DSMZ |
| *Xanthobacter autotrophicus* 7C SF | *Xanthobacter autotrophicus* 7C SF | **DSM 2267** | DSMZ |
| "*Xanthobacter viscosus*" 7d<sup>T</sup> | *Xanthobacter viscosus* 7d<sup>T</sup> | ATCC BAA-298; CIP 108462; **DSM 21355**; KCTC 12306; VKM B-2253 | DSMZ |
| *Xanthobacter autotrophicus* CCUG 44692 | *Xanthobacter autotrophicus* CCUG 44692 | **CCUG 44692** | CCUG |
| *Xanthobacter autotrophicus* GZ29 | *Xanthobacter autotrophicus* GZ29 | **DSM 1393**; LMG 7044 | DSMZ |
| *Xanthobacter autotrophicus* JW33 | *Xanthobacter autotrophicus* JW33 | **DSM 1618**; IFO 14758; NBRC 14758; NCIMB 12468 | DSMZ |
| *Xanthobacter autotrophicus* NCIMB 11171 | *Xanthobacter autotrophicus* NCIMB 11171 | LMD 90.18; NCCB 90018; **NCIMB 11171** | NCIMB |
| *Xanthobacter autotrophicus* V0C-4 |  | DSM | This study; Surface soil sample, Boston, MA, USA |
| *Xanthobacter cornucopiae* V4C-4<sup>T</sup> |  | ATCC; DSM | This study; root and sediment sample, Muddy River, Boston, MA, USA |
| *Xanthobacter flavus* 301<sup>T</sup> | *Xanthobacter flavus* 301<sup>T</sup> | ATCC 35867; BCRC 12271; CCM 4469; CCRC 12271; CIP 105434; **DSM 338**; IFO 14759; JCM 1204; LMG 7045; NBRC 14759; NCAIM B.01946; NCIB 10071; NCIMB 10071; NRRL B-14838; VKM B-2106 | DSMZ |
| *Xanthobacter flavus* GJ10* | *Xanthobacter autotrophicus* GJ10 | ATCC 43050; CCM 4398; **DSM 3874**; NCAIM B.01957; NRRL B-14837 | DSMZ |
| *Xanthobacter flavus* R-10 | *Xanthobacter* sp. R-10 | **DSM 14521** | DSMZ |
| *Xanthobacter flavus* SS-20 | *Xanthobacter* sp. SS-20 | IFO 15493; **NBRC 15493** | NBRC |
| *Xanthobacter flavus* SS-22 | *Xanthobacter* sp. SS-22 | IFO 15494; **NBRC 15494** | NBRC |
| *Xanthobacter flavus* V2C-3 |  | DSM | This study; Water sample, Muddy River, Boston, MA |
| *Xanthobacter flavus* V4C-10 |  | DSM | This study; root and sediment sample, Muddy River, Boston, MA, USA |
| *Xanthobacter flavus* V4C-7 |  | DSM | This study; root and sediment sample, Muddy River, Boston, MA, USA |
| *Xanthobacter flavus* V4C-9 |  | DSM | This study; root and sediment sample, Muddy River, Boston, MA, USA |
| *Xanthobacter lutulentifluminis* V3C-3<sup>T</sup> |  | ATCC; DSM | This study; Sediment sample, Muddy River, Boston, MA, USA |
| *Xanthobacter nonsaccharivorans* 14g<sup>T</sup> | *Xanthobacter autotrophicus* 14g | CIP 105432; **DSM 431**; IAM 12635; JCM 1201; NCIB 10811; NCIMB 10811 | DSMZ |
| *Xanthobacter oligotrophicus* 23A | *Xanthobacter autotrophicus* 23A | CIP 105433; **DSM 685**; JCM 1203 | DSMZ |
| *Xanthobacter pseudotagetidis* KA<sup>T</sup> | *Xanthobacter tagetidis* KA | **DSM 11602; KCTC** | DSMZ |
| *Xanthobacter sediminis* V3B-7B |  | DSM | This study; Sediment sample, Muddy River, Boston, MA, USA |
| *Xanthobacter sediminis* V8C-5<sup>T</sup> |  | ATCC; DSM | This study; Sediment sample, Muddy River, Boston, MA, USA |
| *Xanthobacter tagetidis* A2 | *Xanthobacter tagetidis* A2 | **ATCC 700315** | ATCC |
| *Xanthobacter tagetidis* TagT2C<sup>T</sup> | *Xanthobacter tagetidis* TagT2C<sup>T</sup> | ATCC 700314; **DSM 11105**; NCIMB 13547 | DSMZ |
| *Xanthobacter toluenivorans* T101<sup>T</sup> | *Xanthobacter autotrophicus* T101 | **ATCC 700551; DSM** | ATCC |
| *Xanthobacter toluenivorans* T102 | *Xanthobacter autotrophicus* T102 | **ATCC 700552** | ATCC |
| *Xanthobacter variabilis* V4C-8<sup>T</sup> |  | ATCC; DSM | This study; root and sediment sample, Muddy River, Boston, MA, USA |
| *Xanthobacter variabilis* V13B-9B |  | DSM | This study; Silt and sediment sample, Muddy River, Boston, MA, USA |
| *Xanthobacter variabilis* V13C-7B |  | DSM | This study; Silt and sediment sample, Muddy River, Boston, MA, USA |
| *Xanthobacter versatilis* CB6 | *Xanthobacter* sp. CB6 | **DSM 14520** | DSMZ |
| *Xanthobacter versatilis* Py2<sup>T*</sup> | *Xanthobacter autotrophicus* Py2 | **ATCC BAA-1158; DSM** | ATCC |
| *Xanthobacter versatilis* V1C-2 |  | DSM | This study; Flower bed soil and mulch sample, Boston, MA, USA |
| *Xanthobacter versatilis* V1C-6B |  | DSM | This study; Flower bed soil and mulch sample, Boston, MA, USA |
| *Xanthobacter versatilis* V2C-9 |  | DSM | This study; Water sample, Muddy River, Boston, MA |
| *Xanthobacter versatilis* V3C-2 |  | DSM | This study; Sediment sample, Muddy River, Boston, MA, USA |
| *Xanthobacter versatilis* V3C-4 |  | DSM | This study; Sediment sample, Muddy River, Boston, MA, USA |
| *Xanthobacter versatilis* V7C-1B |  | DSM | This study; Sediment sample, Muddy River, Boston, MA, USA |
| *Xanthobacter versatilis* V7C-4 |  | DSM | This study; Sediment sample, Muddy River, Boston, MA, USA |
| *Xanthobacter wiegelii* CB2 | *Xanthobacter* sp. CB2 | **DSM 14517** | DSMZ |
| *Xanthobacter wiegelii* NCIMB 11399 | *Xanthobacter autotrophicus* NCIMB 11399 | **NCIMB 11399** | NCIMB |
| *Xanthobacter wiegelii* RH 10<sup>T</sup> | *Xanthobacter autotrophicus* RH 10 | CIP 105437; **DSM 597**; JCM 7864; TK 0519 | DSMZ |
| *Roseixanthobacter finlandensis* VTT E-85241<sup>T</sup> | "*Xanthobacter polyaromaticivorans*" VTT E-85241 | **VTT E-85241**; BIO GISA 20; DSM | VTT |
| *Roseixanthobacter glucoisosaccharinivorans* VTT E-85242<sup>T</sup> | "*Xanthobacter polyaromaticivorans*" VTT E-85242 | **VTT E-85242**; BIO GISA 21; DSM | VTT |
| *Roseixanthobacter liquoris* VTT E-85238<sup>T</sup> | "*Xanthobacter polyaromaticivorans*" VTT E-85238 | **VTT E-85238**; BIO GISA 16; DSM | VTT |
| *Roseixanthobacter pseudopolyaromaticivorans* VTT E-85237 | "*Xanthobacter polyaromaticivorans*" VTT E-85237 | VTT E-85237; BIO GISA 14 | VTT |
| *Roseixanthobacter pseudopolyaromaticivorans* VTT E-85239 | "*Xanthobacter polyaromaticivorans*" VTT E-85239 | VTT E-85239; BIO GISA 18A | VTT |
| *Roseixanthobacter pseudopolyaromaticivorans* VTT E-85240<sup>T</sup> | "*Xanthobacter polyaromaticivorans*" VTT E-85240 | VTT E-85240; BIO GISA 19; DSM | VTT |
| *Roseixanthobacter psychrophilus* W30<sup>T</sup> | *Xanthobacter* sp. W30 | DSM 24535; KCTC | DSMZ |

After diluting the cultures 1/100 in fresh IMM every 2 weeks for approximately 10 weeks, cultures were diluted to 10^-2^, 10^-4^, and 10^-6^ with IMM, and 100 uL of each was plated onto rich media 1.5% agar plates of 2xM1. Following 7 days of growth at 30°C on the plates, single colonies were restruck and isolated. After examination using phase contrast microscopy, some cultures were restruck again on 2xM1 1.5% agar plates down to single colonies to obtain pure isolated cultures. Cultures were then grown in 5 mL liquid 2xM1 at 30°C before sequencing and long-term storage at -80°C in 40% v/v glycerol.

We deposited all Xanthobacter environmental isolates from this study in the Deutsche Sammlung von Mikroorganismen und Zellkulturen GmbH/German Collection of Microorganisms and Cell Cultures GmbH (DSMZ). Type strains were also deposited in the American Type Culture Collection (ATCC).

### Growth media and physiological condition testing

A standard rich growth medium (2xM1) was used to grow the strains heterotrophically in the laboratory. The 2xM1 medium is a double-concentrated version of the M1 (nutrient broth) recipe from the DSMZ (47), which we found led to faster and denser growth than the unmodified version. The 2xM1 medium contains 10 g peptone (Gibco Bacto Peptone, 211677) and 6 g meat extract powder (Gibco Difco Beef Extract Powder, 212303), and for plates, 15 g agar (Gibco Difco Agar, 214010) per litre of MilliQ water is added. The 2xM1 medium was autoclaved for sterilization. For salinity physiological condition testing, the 2xM1 medium was modified to include either 0.5%, 1%, 1.5%, 2%, 3%, 4%, 5%, or 6% NaCl w/v. For pH physiological testing, the 2xM1 medium was modified to include 50 mM buffering with citric acid buffer for pH 3, pH 4, and pH 5 (Na_2_HPO_4_, citric acid), potassium phosphate buffer for pH 6, 7, and 8 (K_2_HPO_4_, KH_2_PO_4_), and carbonate-bicarbonate buffer for pH 9 and pH 10 (NaHCO_3_, Na_2_CO_3_).

For additional heterotrophic media testing, R2A was also included. The R2A medium recipe was taken from the DSMZ (48) and includes 0.5 g yeast extract (Gibco Bacto Yeast Extract, 212750), 0.5 g proteose peptone (Gibco Bacto Proteose Peptone No. 3, 211693), 0.5 g casamino acids (Gibco Bacto Casamino Acids, 223050), 0.5 g glucose (Sigma Aldrich, G7021-100G), 0.5 g soluble starch (Sigma Aldrich S9765-100G), 0.3 g sodium pyruvate (Alfa Aesar, A11148), 0.3 g K_2_HPO_4_, 0.05 g MgSO_4_ • 7 H_2_O per litre of MilliQ water. The R2A medium was sterilized by autoclaving.

During nitrogen-fixing, hydrogen-oxidizing chemolithoautotrophic growth, a standard inorganic minimal medium (IMM) was used (19). The IMM contains (added in order): 1 g K_2_HPO4, 0.5 g KH_2_PO_4_, 2 g NaHCO_3_, 0.1 g MgSO_4_ • 7 H_2_O, 20 mL of a pre-dissolved 2 g/L water CaSO_4_ • 2 H_2_O solution, 0.01 g FeSO_4_ • 7 H_2_O, and 1 mL of a filter sterilized trace mineral medium solution (TMM) per litre of MilliQ water. The TMM contains: 2.8 g H_3_BO_3_, 2.1 g MnSO_4_ • 4 H_2_O, 0.75 g Na_2_MoO_4_ • 4 H_2_O, 0.24 g ZnSO_4_ • 7 H_2_O, 0.04 g Cu(NO_3_)_2_ • 3 H_2_O, and 0.13 g NiSO_4_ • 6 H_2_O. The IMM medium was either filter-sterilized or autoclaved for sterilization. After sterilization, 10 mL of a vitamin solution (ATCC Cat No. MD-VS) was added.

For carbon substrate utilization testing, a modified minimal mineral medium (4M) from Wiegel *et al.,* 1978 (1) was used and contained: 5.7 g K_2_HPO_4_, 1.95 g KH_2_PO_4_, 3 g (NH_4_)SO_4_, 0.1 g MgSO_4_ • 7 H_2_O, 20 mL of a pre-dissolved 2 g/L water CaSO_4_ • 2 H_2_O solution, 0.01 g FeSO_4_ • 7 H_2_O, 1 mL of a filter sterilized trace mineral medium solution (TMM above), and 10 mL of a vitamin solution (ATCC Cat No. MD-VS) per litre of MilliQ water. The 4M medium was filter sterilized. Individual carbon substrate sources (Supplementary Table 1) were added at 0.5% w/v to the minimal medium. For mixed carbon testing on the substrates, the base minimal medium also contained 0.01% succinate (or 0.01% pyruvate for *X. toluenivorans* T101^T^) in addition to the 0.5% carbon substrate being tested. For some select substrates, a dilution series from 1%, 0.5%, 0.1%, 0.05%, and 0.01% w/v substrate in base minimal medium was also tested. For the physiological testing in high-throughput liquid culture, 500 uL of each condition was inoculated with 0.01 OD_600_ of culture in 96-well plates, and plates were shaken at 999 RPM with an orbital shaker. A positive result for physiological testing was determined by an increase of at least one doubling in OD_600_ of the culture measured on a plate reader.

Xanthobacter and Roseixanthobacter gen. nov. strains were grown under standard heterotrophic or nitrogen-fixing chemolithoautotrophic conditions at 30°C. For temperature physiological testing, spot plates of serially diluted culture on 2xM1 were grown in incubators at 4°C, 10°C, 15°C, 20°C, 25°C, 30°C, 37°C, and 42°C.

Larger cultures were shaken at 200-250 RPM in typically 5 mL tube volumes or 50 mL flask volumes in 2xM1 at 30°C. Optical density measurements of larger cultures were taken on a photospectrometer at OD_600_.

### Genome sequencing and assembly

Strains were grown in 5 mL 2xM1 shaking at 30°C until saturation of the slowest cultures (7 days). For genomic DNA isolation, 3 mL of saturated culture was pelleted and used according to manufacturer instructions following the MasterPure Complete DNA and RNA Purification Kit (Lucigen MC85200). Isolated genomic DNA was quantified using a NanoDrop and used as input for the Illumina DNA Prep Tagmentation Kit (formerly Nextera DNA Flex) using the Illumina DNA/RNA UD Indexes (sets A and B). Libraries were pooled, and the Agilent 2100 BioAnalyzer system for the MiSeq pool or the Agilent TapeStation 4200 (with Agilent TapeStation High Sensitivity D1000 tape) for the NovaSeq pool were used for quality control. Pools were loaded and sequenced either with the MiSeq (MiSeq Reagent Kit v3, 150-cycle) system in-house (*X. nonsaccharivorans* 14g^T^, *X. oligotrophicus* 23A, *X. autotrophicus* 7C^T^, *X. autotrophicus* 7C SF, *X. autotrophicus* GZ29, *X. autotrophicus* JW33) or with the NovaSeq 6000 system (NovaSeq Reagent Kit SP v1.5, 150-cycle) at the Harvard Medical School Biopolymer Facility (all other strains in Table 1 except for *X. flavus* GJ10 and *X. versatilis* Py2^T^).

Reads were then analyzed using FastQC v0.12.1 (49) and processed using Trimmomatic v0.39 (50) under default settings. Paired, trimmed reads were used as input into the Unicycler assembler v0.5.0 (51) (a SPAdes (52) optimizer, using SPAdes v3.15.4) under default settings. The assembled genome for *Xanthobacter variabilis* V4C-8^T^ was found to be missing a large chunk of the genome following Unicycler assembly and was instead assembled using SPAdes alone. All contigs smaller than 200 bp were removed prior to NCBI submission. Genome assemblies were then uploaded to NCBI GenBank and annotated using NCBI’s automatic prokaryotic genome annotation pipeline (PGAP) tool (53).

The 16S rRNA genes were extracted from all newly sequenced genomes using barrnap v0.9 (54) (with Bedtools 2.27.1 (55), Hmmer 3.3.2 (56), Perl 5.32.1) and uploaded to NCBI. A complete 16S rRNA gene could be extracted from the SPAdes assembled *X. variabilis* V4C-8^T^ genome but not the Unicycler assembled version. Only partial 16S rRNA genes could be isolated from *X. autotrophicus* V0C-4 and *X. versatilis* V2C-9 regardless of assembly method, so Unicycler assembled versions were used.

Sequencing statistics and accession numbers for the newly sequenced genomes were obtained suing BBMap v39.06 (57) (for N50), NCBI’s PGAP (for gene counts), and BWA-MEM v0.7.17 (58) / Samtools v1.16.1(59) (for coverage, mapping paired-end reads back to assembled genomes). Accession numbers and summary statistics for new and all other prior sequenced genomes used for bioinformatic analyses are listed in Supplementary Table 2.

### Genome Taxonomy Database Toolkit (GTDB-Tk) classification

For initially confirming acquired Xanthobacter repository strains were Xanthobacter species and to identify the Xanthobacter strains from our enriched community isolates, genomes were run through the GTDB-Tk v2.3.2 software (60) using the release 214 dataset. All genomes that were identified as belonging to the genus Xanthobacter were used in downstream applications. Notably, strains that were identified as “*Xanthobacter polyaromaticivorans*” and one “*Xanthobacter sp.*” (W30) were assigned to the unnamed genus “NCHE01” (which we propose as Roseixanthobacter gen. nov.) instead of Xanthobacter with GTDB and were still used for downstream analysis based on prior taxonomic assignment.

### Whole genome phylogenomic analysis and species-level assignment

Newly and prior sequenced genomes (Supplementary Table 2) belonging to Xanthobacter and Roseixanthobacter were used for comparative phylogenomic analyses. Additionally, type strains from the highly related genera Aquabacter and Azorhizobium, and the less related Ancylobacter were included (2,3,44).

*Pseudoxanthobacter soli* CC4^T^ was included as an outgroup. In total, 40 species/genomes were used for the type strain phylogenomic analyses and 104 genomes were used for the phylogenomic analyses with all strains.

The 120 core, conserved, single-copy bacterial proteins (bac120) from the GTDB taxonomic analysis pipeline were used for protein-based phylogenomic analysis. These proteins were extracted using the GTDB-Tk identify tool (including -- write_single_copy_markers) and default settings. Proteins were aligned separately for the 108 strains using MUSCLE (61) in MEGA-X v10.2 (62) (default settings) and concatenated using catfasta2phyml v1.2.0 (63) (FASTA format, concatenation). The concatenated alignment of the bac120 protein set was then used in the MEGA-X software for phylogenomic analysis. Neighbor-joining (NJ) and maximum parsimony (MP) trees were generated under default settings (NJ (64) – amino acid, Poisson correction method (65), uniform rates, partial deletion, 95% cutoff; MP (66) – amino acid, partial deletion, 95% cutoff, search – Subtree-Pruning-Regrafting/SPR, 10 initial trees, 1 search 1 search level, 100 trees to retain) and bootstrapped 1000 times (67). The model finder function on MEGA-X was used to identify the best model for the bac120 alignment for maximum likelihood (ML) analysis, and the top model (FG + G(5) + I) (68) was chosen to create a maximum likelihood tree (partial deletion, 95% cutoff) and bootstrapped 500 times (and not 1000 due to computational constraints). All trees were rooted using *Pseudoxanthobacter soli* CC4^T^. All bac120 trees with all strains used a total of 44518 amino acid positions in the final dataset.

For further species-level assignment confirmation, average nucleotide identity/ANI (FastANI v0.1.3 (69)) and digital DNA-DNA hybridization/dDDH (Genome to Genome Distance Calculator v3.0 (70,71)) values were calculated using the 87 Xanthobacter and Roseixanthobacter gen. nov. genomes. Species-level boundaries were identified using a 95% cutoff for ANI (72–74). Additional validation came from dDDH values calculated using the 70% species-level threshold (71,75–77); however, the 95% ANI threshold was used for delineating species boundaries in cases where the dDDH value was close to but below 70%, yet the ANI value was above 95%. Unnamed species clusters were assigned a type strain, which was used for phenotypic testing and characterization, as well as for novel species descriptions.

A 16S rRNA phylogenetic tree was generated using 42 type strains in total for comparison with effectively but invalidly published species of Xanthobacter. The 16S rRNA sequences were aligned using MUSCLE (78) (default settings) and used in the MEGA-X v10.2 software (62) for phylogenetic analysis. Neighbor-joining and maximum parsimony trees were generated under default settings (NJ (64) – nucleotide, maximum composite likelihood, uniform rates, partial deletion, 95% cutoff; MP (66) – nucleotide, partial deletion, 95% cutoff, search – Subtree-Pruning-Regrafting/SPR, 10 initial trees, 1 search 1 search level, 100 trees to retain) and bootstrapped 1000 times. The model finder function on MEGA-X was used to identify the best maximum likelihood model (K2 + G(5) + I) (79) to create a maximum likelihood tree (partial deletion, 95% cutoff), and the resulting tree was bootstrapped 1000 times (67). All trees were rooted using *Pseudoxanthobacter soli* CC4^T^. All 16S rRNA trees used a total of 1354 nucleotide positions in the final dataset. MAFFT v7.511 (80) was used for the 16S rRNA gene alignment visualizations in Supplementary Figures 7 and 8.

### Phenotypic testing and characterization

To test for motility, type and representative strains were grown within a semisolid motility agar assay (81,82). The medium consisted of 0.3% agar, 150 uM 2,3,5-triphenyltetrazolium chloride/TTC (enhances visibility of cells as they uptake colorless TTC and convert it into an insoluble pigmented compound retained within the cell (83)), and 0.3% w/v or v/v of succinate, fumarate, gluconate, methanol, ethanol, propanol, or isopropanol in the 4M minimal medium described above. After growing strains on 2xM1 plates for 3 days at 30°C, a needle was used to pick up a few colonies and was inserted into the semisolid agar. Tubes were left upright for 2 weeks, with observations after 5 days and the 14 days. A positive result was indicated when a pink/red hue was observed across the tube beyond the inoculation line. We also searched through PGAP genome annotations of the type strains for flagellar motility pathway genes (*flg, fli, flh, mot* genes (84)).

To determine enzymatic profiles, the API ZYM (85) and API 20NE testing strips were used. Type and chosen representative strains were grown on 2xM1 plates (except for *X. toluenivorans* T101^T^*, X. agilis* SA35^T^*, X. sediminis* V8C-5^T^*, and X. albus* V0C-6^T^, which were grown in 2xM1 liquid and spun down) for 3 days prior to API strip testing and resuspended using the API Suspension medium. API ZYM activities were measured after 4.5 hours according to manufacturer instructions. API 20NE activities were measured after 24 hours, 48 hours, and 72 hours according to manufacturer instructions. A final timepoint after 7 days had passed was also taken, given the slow growth of many Xanthobacter, especially under suboptimal media conditions. Positive activities are only reported within the 72-hour window; however, all results can be found in Supplementary Tables 13 and 14.

Xanthobacter are known to produce an exopolysaccharide slime under some conditions (1,3,86), including when grown on media containing tricarboxylic acid cycle intermediates like succinate. To determine slime production, we spot-plated a serial dilution series of type and representative strains grown from rich media (2xM1 for Xanthobacter, R2A for Roseixanthobacter gen. nov.) onto 4M minimal media plates containing 0.5% w/v succinate and rich 2xM1 plates. After 7 days of growth at 30°C, spots and colonies were observed for the constitutive or inducible production of slime in comparison to growth on rich 2xM1 plates.

For determining cell size and pleomorphic branching characteristics, cells from each type and representative strain were grown in either rich 2xM1 media or 4M minimal media with 0.5% w/v succinate, a known inducer of branching morphology in Xanthobacter (1,3,21). Cell sizes were only calculated for rod-shaped cells in the rich 2xM1 media condition. After being grown to saturation, for the 2xM1 condition, cells were passaged 1/10 overnight in fresh 2xM1 prior to observation. For observing induced branching on succinate, cultures were grown in the succinate condition for 7 days to acquire adequate growth for imaging. Branching was observed after 24 hours in succinate, but cells were too dilute to acquire adequate images, and thus images were acquired after 7 days of growth instead. Cell size was estimated using Quantitative Phase Imaging (QPI). Imaging was performed on a Nikon Eclipse TI microscope modified for QPI (with a Phasics SID4BIO) as described in Liu *et al.,* 2020 .(87) Images were acquired using the Nikon Plan Apo 100X Phase objective with the 1.5X magnifier with a halogen lamp as the light source. A 10 µL volume of culture was loaded onto a 1% agarose pad assembled on a glass coverslip, and 64 fields of view were acquired automatically for each sample. Images were processed using the ceQPM MATLAB toolbox (v1.0.2) (87). Image segmentation was carried out using cellpose-3.0.8 (88,89). The ’deepbacs_cp3’ pre-trained model was used as a starting point. The cellpose GUI was then used to fine-tune the model using the output from the ceQPM pipeline, and this trained model was used for image segmentation with no additional parameters. The masks obtained from this pipeline were then filtered to remove clusters containing multiple cells or debris. Finally, the scikit-image-0.22.0 library was used to quantify the cell size metrics. In total, 150 cells were sampled from each species to compute the final cell length and width statistics (n=145 for *X. oligotrophicus*).

## RESULTS AND DISCUSSION

### Genome sequencing and initial taxonomic classification of Xanthobacter strains

Despite the ubiquitous presence of Xanthobacter in environments and ecosystems (3) and an immense interest in industrial, academic, and environmental applications, we our understanding of the diversity and evolutionary history of the genus is limited. The majority of Xanthobacter strains currently available from cell culture repositories lack genome sequences and taxonomic validation. Additionally, despite their abundance worldwide and in the literature, there were only eight validly published species of Xanthobacter (1,5,7,9–12).

Thus, we aimed to generate a larger collection of Xanthobacter genomes and strains to better capture and understand the natural diversity and evolutionary history of the genus. We first sought to obtain every Xanthobacter strain we could from repositories around the world, totaling 37 strains from the ATCC, CCUG, DSMZ, NBRC, NCIMB, and VTT. Additionally, we carried out an environmental isolation campaign from soil, water, and sediment samples to enrich for novel hydrogen-oxidizing, nitrogen-fixing, chemolithoautotrophic communities and isolates, with the aim of bolstering known Xanthobacter strains (90). In total, we isolated over 120 novel environmental isolates, of which 26 were ultimately identified as Xanthobacter strains as described below.

With the strains in hand, we sequenced these repository strains and isolates, obtaining draft genomes with 55x to 274x coverage depending on the strain, all with G+C% content in the mid to high 60s and genome sizes between 4.5 Mb to 5.7 Mb. We then used the Genome Taxonomy Database Toolkit (GTDB-Tk) to find an initial taxonomic classification for these genomes (60). The GTDB-Tk employs an alignment-based method using 120 core, conserved, single-copy bacterial genes (bac120) to obtain down to a genus-level classification. Afterward, GTDB-Tk uses the alignment-free average nucleotide identity method (ANI) to quickly classify genomes to the species-level. The GTDB-Tk species cluster assignments we obtained (Table 2) contained several interesting observations indicating that Xanthobacter taxonomy needed updating.

**Table 2.**
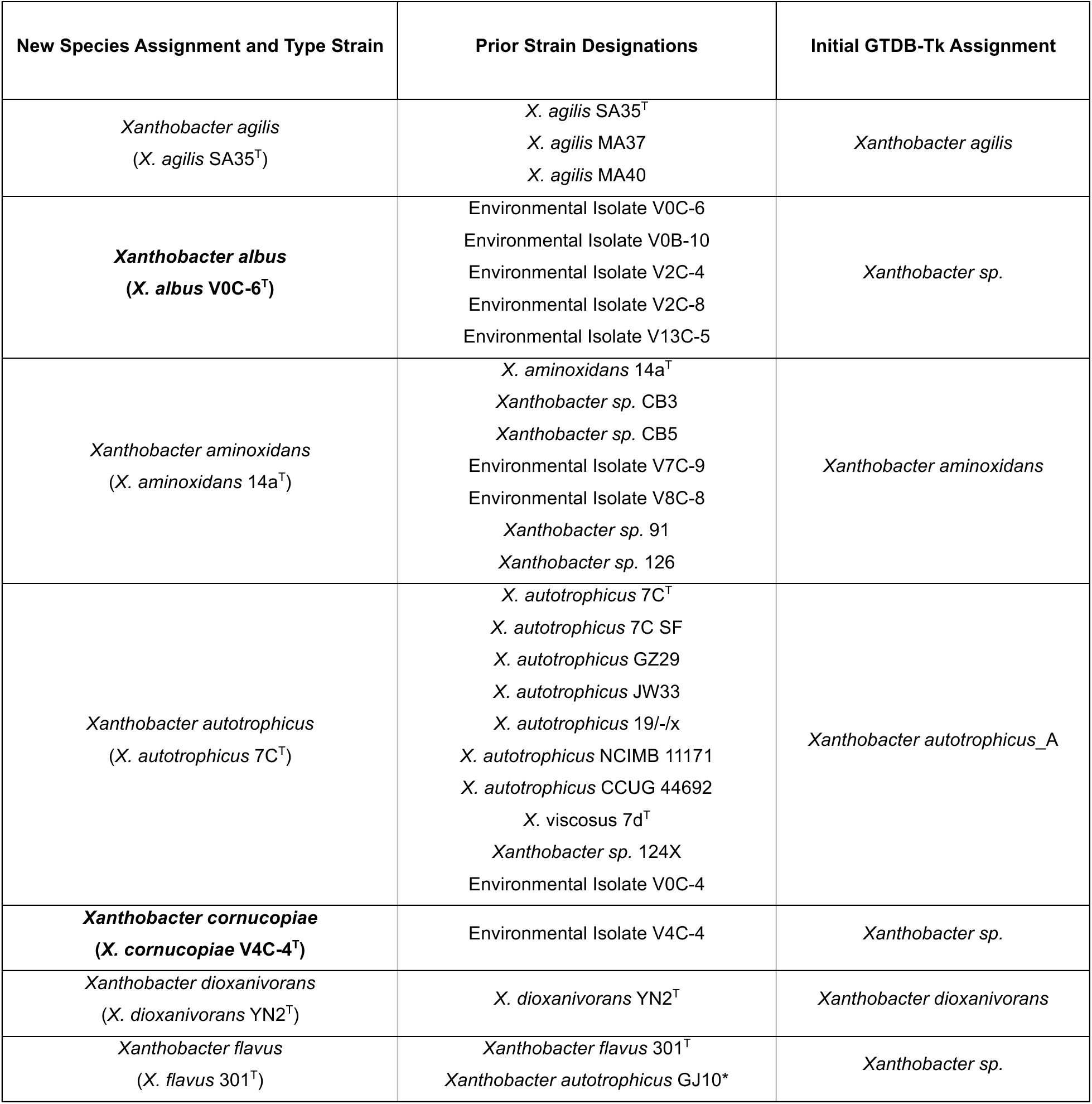
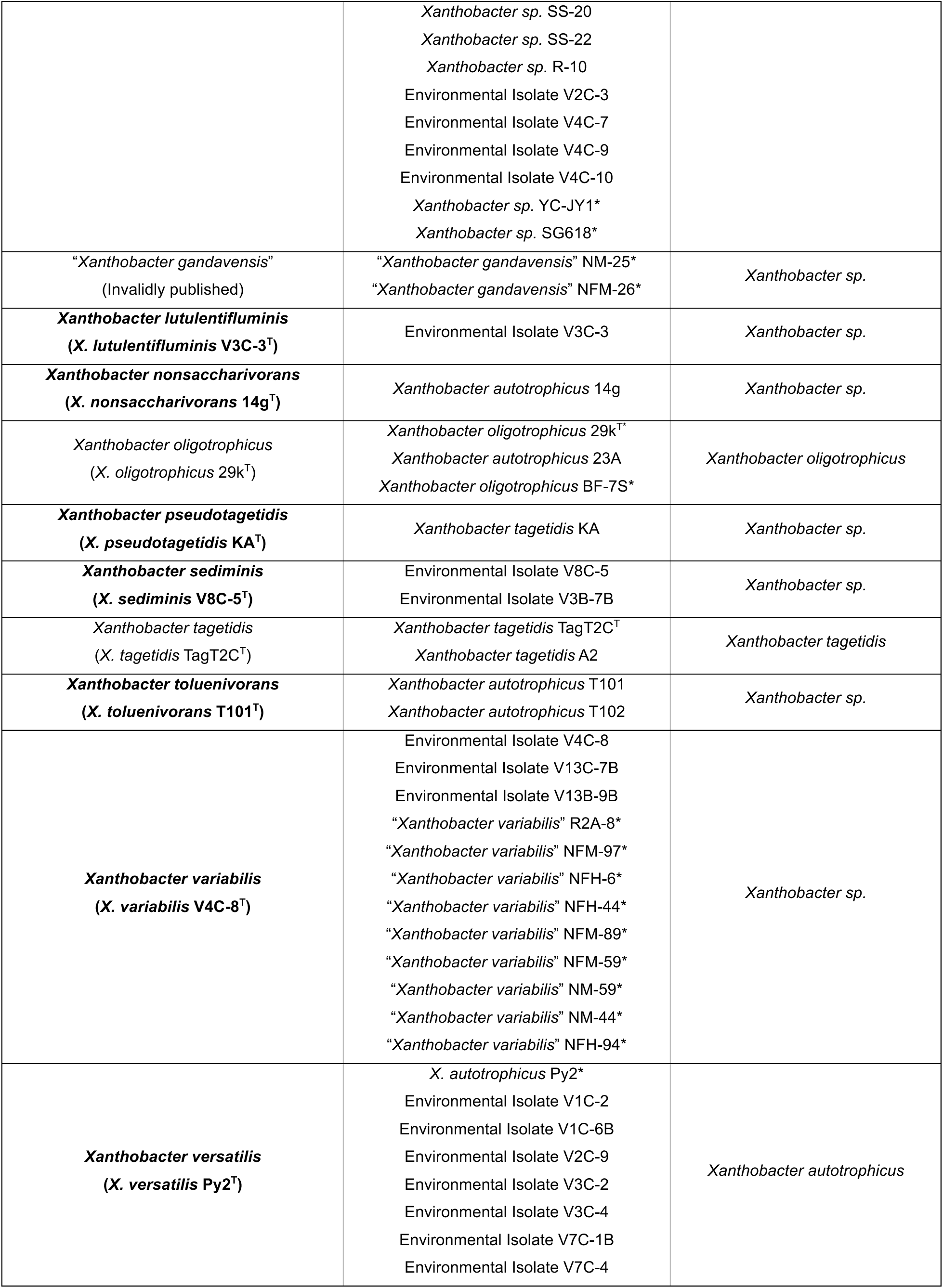
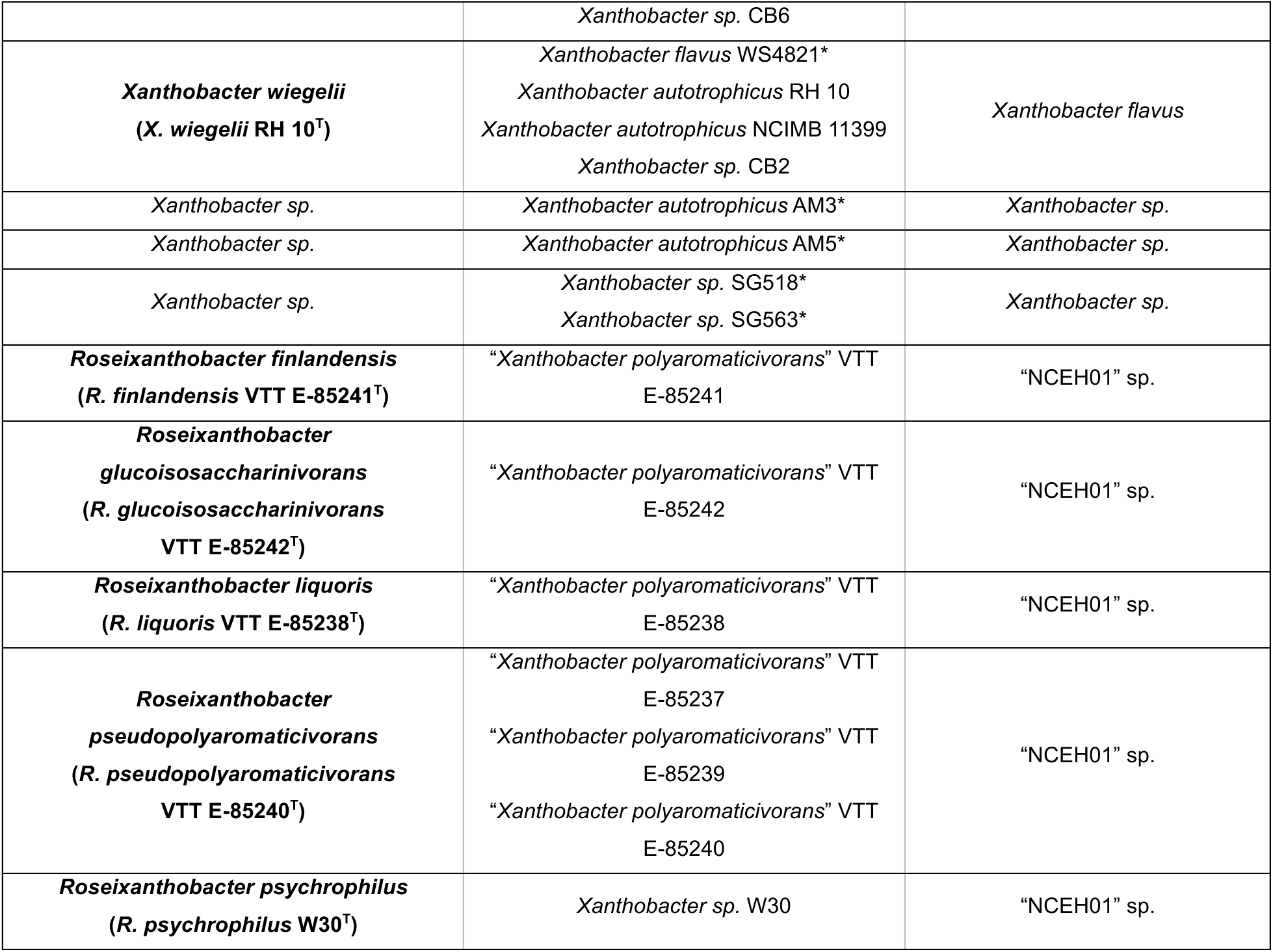
Xanthobacter strains can be initially classified into species clusters using the GTDB-Tk. The table includes the initial GTDB-Tk taxonomic classification of our sequenced repository Xanthobacter strains, other prior sequenced Xanthobacter genomes, and our novel environmental isolates. Species in bold are the novel species described in this study. Strains indicated with an asterisk are genomes that were not sequenced in this study but were available for comparison.

First, two distinct “*X. autotrophicus*” clusters are formed. One cluster, “*X. autotrophicus_A,”* contains the type strain *X. autotrophicus* 7C^T^ and many strains in the literature considered as canonical *X. autotrophicus* (such as GZ29 and JW33). Since this cluster contains the *X. autotrophicus* 7C^T^ type strain, it should be considered the true *X. autotrophicus* cluster. Surprisingly, *X. viscosus* 7d^T^ (which we obtained from DSMZ as DSM 21355^T^) was classified as belonging to this true *X. autotrophicus* cluster and not its separate species. The second cluster was dubbed “*X. autotrophicus*” in the GTDB and contained the well-studied strain Py2—previously considered an *X. autotrophicus*—and many novel environmental isolates. Given this cluster is separate from the *X. autotrophicus* 7C^T^ type strain species cluster, we considered it a separate species, and renamed it *X. versatilis* sp. nov. Along similar lines, another exclusion was that the type strain *X. flavus* 301^T^ was missing from the GTDB release 214, thus the GTDB “*X. flavus”* does not contain the type strain. Instead, a separate *“Xanthobacter sp.”* cluster that contained the *X. flavus* 301^T^ type strain is the true *X. flavus* cluster. The prior incorrectly labelled GTDB “*X. flavus*” cluster we treated as a separate species and renamed it to *X. wiegelii* sp. nov. Notably, the well-studied strain GJ10 was classified as *X. flavus* in our analysis and not its prior assignment in the literature—*X. autotrophicus*—and was renamed accordingly.

Many other repository strains formed unnamed species clusters (“*Xanthobacter sp.*”) separate from their prior pre-genomic taxonomic assignment and represent novel Xanthobacter species, including KA^T^ (previously *X. tagetidis,* now *X. pseudotagetidis* sp. nov), 14g^T^ (previously *X. autotrophicus*, now *X. nonsaccharivorans* sp. nov.), and T101^T^ (previously *X. autotrophicus*, now *X. toluenivorans* sp. nov.). Several of our novel environmental isolates also formed unnamed species clusters and represent novel Xanthobacter species: *X. lutulentifluminis* sp. nov (V3C-3^T^), *X. cornucopiae* sp. nov. (V4C-4^T^), *X sediminis* sp. nov. (V8C-5^T^), and *X. albus* sp. nov. (V0C-6^T^). Other known Xanthobacter species were assigned to their own species clusters as expected, albeit some minor changes: *X. agilis* (no changes), *X. oligotrophicus* (gained new strains), *X. dioxanivorans* (no changes), *X. aminoxidans* (gained new strains), and *X. tagetidis* (lost a member). Furthermore, two previously invalidly published Xanthobacter species remained as separate species clusters: *“Xanthobacter gandavensis”* remains without a repository strain but for *X. variabilis* sp. nov. we found novel environmental isolates and can now validly publish the species (with the type strain V4C-8^T^).

Finally, there were strains previously considered Xanthobacter that were not classified as such and were instead classified as an unnamed genus “NCEH01” (a GTDB placeholder name based on metagenomic identifiers). All strains from the VTT repository previously assigned as *“Xanthobacter polyaromaticivorans*” and strain W30 from the DSMZ fell into several species clusters assigned to this novel genus. Strikingly, these strains all are pink-orangish in color (Supplementary Figure 9), instead of the typical yellow color of Xanthobacter and do not fix CO_2_ under hydrogen-oxidizing, nitrogen-fixing conditions like Xanthobacter do in our hands (Table 3). Thus, with this initial GTDB-Tk taxonomic assignment and these stark phenotypic differences, we assigned these strains to a novel genus, which we proposed as Roseixanthobacter gen. nov.

**Table 3.** A summary of phenotypic properties of Xanthobacter and Roseixanthobacter gen. nov. species. All results taken from this study unless indicated otherwise. † Data obtained from Tikhonova et al., 2021; ‡ Data obtained from Wang et al., 2021. Abbreviations for species are: XVE – *Xanthobacter versatilis* sp. nov. Py2^T^; XAU – *Xanthobacter autotrophicus* 7C^T^; XOG – *Xanthobacter oligotrophicus* 29k^T^; XTO – *Xanthobacter toluenivorans* sp. nov. T101^T^; XDI – *Xanthobacter dioxanivorans* YN2^T^; XLU – *Xanthobacter lutulentifluminis* sp. nov. V3C-3^T^; XNS – *Xanthobacter nonsaccharivorans* sp. nov. 14g^T^; XAM – Xanthobacter aminoxidans 14a^T^; XFL – Xanthobacter flavus 301^T^; XWE – Xanthobacter wiegelii sp. nov. RH 10^T^; XPS – *Xanthobacter pseudotagetidis* sp. nov. KA^T^; XTA – *Xanthobacter tagetidis* TagT2C^T^; XCO – *Xanthobacter cornucopiae* sp. nov. V4C-4^T^; XAG – *Xanthobacter agilis* SA35^T^; XSE – *Xanthobacter sediminis* sp. nov. V8C-5^T^; XAL – *Xanthobacter albus* sp. nov. V0C-6^T^; XVA – *Xanthobacter variabilis* sp. nov. V4C-8^T^; RFI – *Roseixanthobacter finlandensis* sp. nov. VTT E-85241^T^; RLI – *Roseixanthobacter liquoris* sp. nov. VTT E-85238^T^; RGL – *Roseixanthobacter glucoisosaccharinivorans* sp. nov. VTT E-85242^T^; RPP – *Roseixanthobacter pseudopolyaromaticivorans* sp. nov. VTT E-85240^T^; RSY – *Roseixanthobacter psychrophilus* sp. nov. W30^T^; CO^2^ Fix – Carbon Dioxide Fixation; N^2^ Fix – Dinitrogen Fixation; GC Content – GC content of the genome sequence; Temp – Temperature (Optimal Temperature); pH – pH (Optimal pH); Salinity % w/v – Salinity % w/v (Optimal Salinity). + indicates a positive result, -indicates a negative result.

| Species | Clade | Color | Slime on Succinate | Branched Morph. on Succinate | Semisolid Agar Motility | CO<sub>2</sub> Fix. | N<sub>2</sub> Fix. | G+C Content | Temp. °C (Opt) | pH (Opt) | Salinity % w/v (Opt) |
| --- | --- | --- | --- | --- | --- | --- | --- | --- | --- | --- | --- |
| XVE Py2<sup>T</sup> | I | Yellow | + | + | - | + | + | 67.3% | 10-37 (30-37) | 5-8 (6) | 0-3 (0) |
| XAU 7C<sup>T</sup> | I | Yellow | + | + | - | + | + | 67.6% | 10-37 (30-37) | 5-8 (6-8) | 0-5 (0.5) |
| <sup>†</sup>XOG 29k<sup>T</sup> | I | Yellow | N/A | + | N/A | + | + | 67.9% | 4-37 (25-30) | 6-8.5 (6.5-7) | 0-0.5 (0) |
| XTO T101<sup>T</sup> | I | Yellow | + | + | - | + | + | 67.6% | 10-30 (30) | 5-6 (6) | 0-1.5 (0) |
| <sup>‡</sup>XDI YN2<sup>T</sup> | I | Yellow | + | N/A | N/A | + | + | 68.0% | 10-40 (30) | 5-8 (7) | 0-1 (0.1) |
| XLU V3C-3<sup>T</sup> | IIA | Yellow | + | + | - | + | + | 69.8% | 15-37 (30-37) | 5-8 (6) | 0-5 (0) |
| XNS 14g<sup>T</sup> | IIA | Yellow | - | + | - | + | + | 69.6% | 15-37 (30-37) | 5-8 (6-7) | 0-5 (0) |
| XAM 14a<sup>T</sup> | IIB | Yellow | + | + | + | + | + | 67.9% | 10-42 (30-37) | 5-8 (5) | 0-3 (0) |
| XFL 301<sup>T</sup> | IIB | Yellow | + | + | + | + | + | 67.9% | 10-42 (30-37) | 5-8 (6-7) | 0-4 (0) |
| XWE RH 10<sup>T</sup> | IIB | Yellow | + | + | + | + | + | 67.7% | 10-37 (30-37) | 5-8 (6) | 0-2 (0) |
| XCO V4C-4<sup>T</sup> | III | Pale Yellow | - | - | + | + | + | 69.2% | 10-37 (30-37) | 5-8 (8) | 0-2 (0.5) |
| XAG SA35<sup>T</sup> | III | Pale Yellow | - | - | + | + | + | 67.4% | 10-37 (30-37) | 5-8 (7) | 0-1.5 (0.5) |
| XSE V8C-5<sup>T</sup> | III | Pale Yellow | - | - | + | + | + | 68.5% | 15-37 (30-37) | 5-8 (8) | 0-6 (0.5) |
| XAL V0C-6<sup>T</sup> | III | Pale Yellow | - | - | + | + | + | 69.3% | 15-37 (30-37) | 5-8 (6-7) | 0-4 (0.5) |
| XVA V4C-8<sup>T</sup> | III | White | - | - | + | + | + | 67.2% | 15-42 (30-37) | 5-8 (8) | 0-6 (1) |
| XPS KA<sup>T</sup> | IV | Yellow | - | - | - | + | + | 69.7% | 15-42 (30-37) | 5-8 (8) | 0-3 (0) |
| XTA TagT2C<sup>T</sup> | IV | Yellow | + | - | - | + | + | 69.6% | 10-37 (30-37) | 6-8 (8) | 0-3 (0.5) |
| RFI E-85238<sup>T</sup> | Ros. | Pink-Orange | - | - | + | - | + | 67.2% | 10-37 (25-30) | 5-8 (5) | 0-2 (0-1) |
| RLI E-85241<sup>T</sup> | Ros. | Pink-Orange | - | - | + | - | + | 65.9% | 10-37 (25-30) | 5-7 (6) | 0-1.5 (0.5) |
| RGL E-85242<sup>T</sup> | Ros. | Pink-Orange | - | - | + | - | + | 66.0% | 10-37 (25-30) | 5-8 (5) | 0-3 (1) |
| RPP E-85240<sup>T</sup> | Ros. | Pink-Orange | - | - | + | - | + | 65.7% | 10-37 (25-30) | 5-7 (6) | 0-1.5 (0-0.5) |
| RSY W30<sup>T</sup> | Ros. | Pink-Orange | - | - | - | - | + | 65.3% | 10-30 (25-30) | 5-8 (5) | 0-2 (0.5) |

### Genus- and species-level delineation of Xanthobacter and Roseixanthobacter

Following these initial taxonomic classifications by the GTDB, we sought to verify species-level delineations and their evolutionary history using a polyphasic phylogenomic approach. First, we compared all Xanthobacter and Roseixanthobacter gen. nov. genomes available using a concatenated alignment of the bac120 core, conserved, single-copy bacterial genes and evaluated their evolutionary history using phylogenomic trees (Figure 1 for type strain tree with maximum likelihood, Figure 2 for all strains with maximum likelihood, Supplementary Figures 1-2 for all strains with neighbor-joining, and maximum parsimony trees). We included closely related type and representative strains from the Azorhizobium, Aquabacter, and Ancylobacter genera in the analysis (3,44,45), and used *Pseudoxanthobacter soli* CC4^T^ as an outgroup to root the trees.

**Figure 1.**
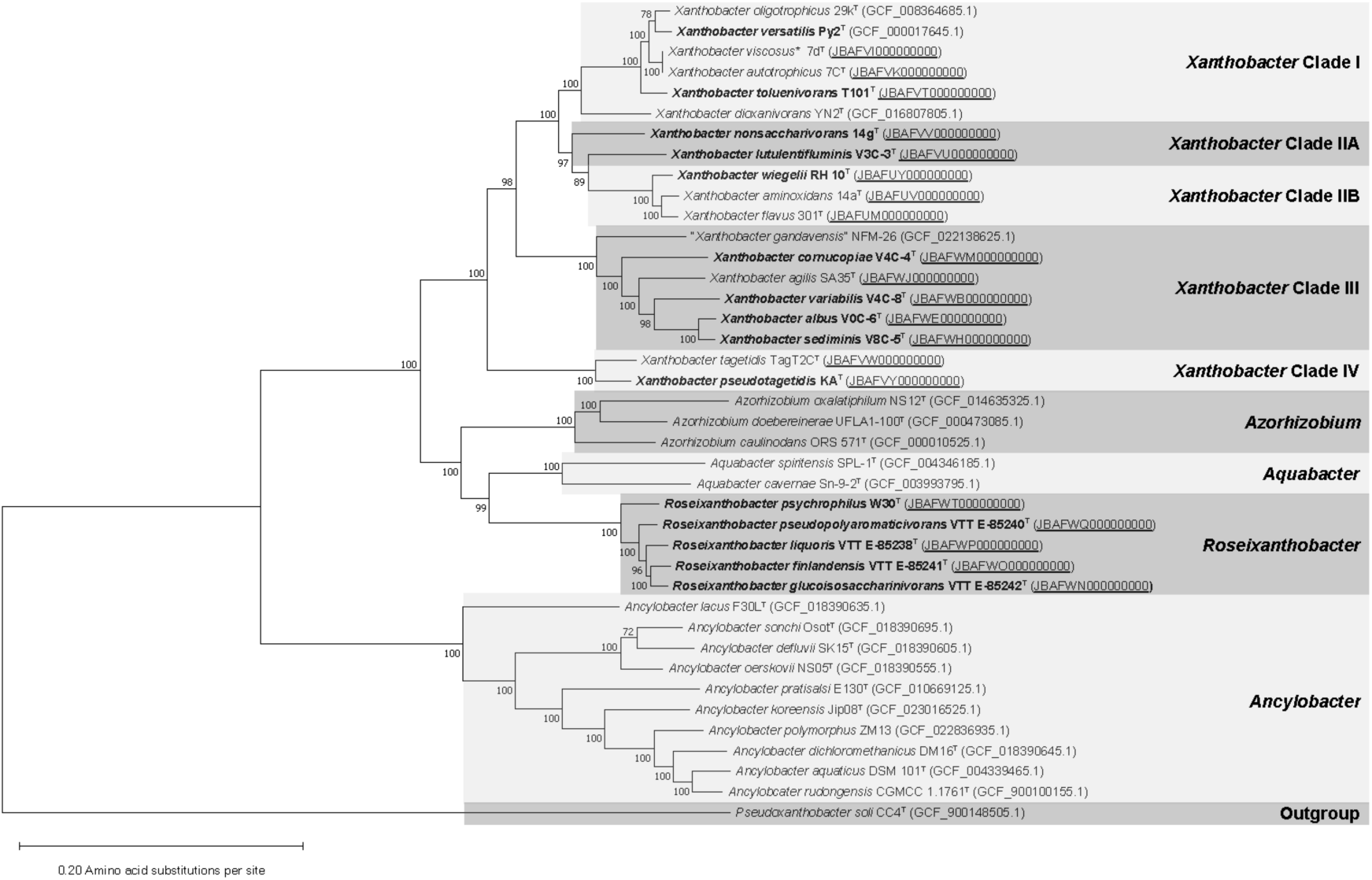
The evolutionary history of the Xanthobacter and Roseixanthobacter gen. nov. species type strains inferred using the maximum likelihood method with 120 core, conserved, single-copy bacterial genes indicates robust evolutionary clades. The phylogenomic tree was generated using MEGA-X with the FG+G(5)+I model with the partial deletion (95% cutoff) option and was bootstrapped 500 times (with bootstrap percentages shown next to the branch). Novel species described in this paper are bolded while genome sequence accessions that we sequenced in this study are underlined. Related genera Azorhizobium, Aquabacter, and Ancylobacter are included. *Pseudoxanthobacter soli* CC4^T^ is used as an outgroup. The scale bar represents 0.2 amino acid substitutions per site.

**Figure 2.**
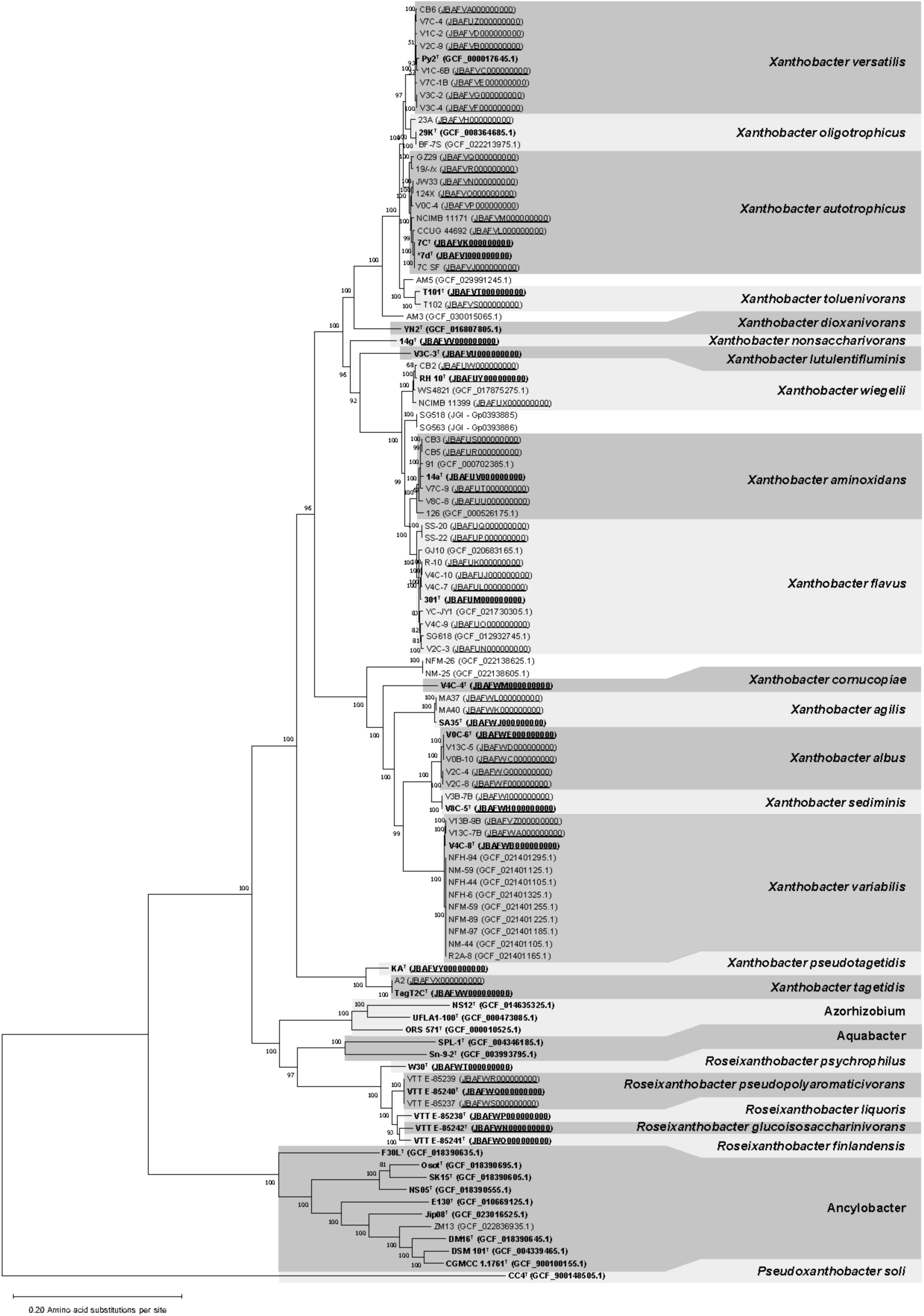
The evolutionary history of all Xanthobacter and Roseixanthobacter gen. nov. strains inferred using the maximum likelihood method with 120 core, conserved, single-copy bacterial genes demonstrates robust species-level boundaries. The phylogenomic tree was generated using MEGA-X with the FG+G(5)+I maximum likelihood model with the partial deletion (95% cutoff) option and was bootstrapped 500 times (with bootstrap percentages shown next to the branch). Type strains are bolded and genomes that we sequenced in this study are underlined. Related genera Azorhizobium, Aquabacter, and Ancylobacter are included. *Pseudoxanthobacter soli* CC4^T^ is used as an outgroup. The scale bar represents 0.2 amino acid substitutions per site.

We found that placements of species and strains within the trees were robust and reproducible, as determined through bootstrapping. Xanthobacter formed a monophyletic group that could be delineated into clades with similar phenotypic properties, as described later in this study. The Roseixanthobacter gen. nov. species formed their own branch alongside the Azorhizobium and Aquabacter genera, further reiterating that these strains belong to a novel genus within the Xanthobacteraceae and are not members of the genus Xanthobacter. The phylogenomic trees show marked improvement over prior 16S rRNA phylogenetic trees from the literature; in the 16S rRNA phylogenetic trees Xanthobacter does not form a monophyletic group (44,45).

Instead, when using 16S rRNA gene data alone, there is a polyphyletic group of Xanthobacter-Azorhizobium-Aquabacter, and species- and genus-level boundaries cannot be robustly demarcated. We also generated 16S rRNA phylogenetic trees with our novel species to draw comparisons to past literature and invalidly published Xanthobacter species that only have 16S rRNA gene data available (Supplementary Figures 3-5). Just as in previous literature attempts, we were unable to produce robust Xanthobacter monophyletic groups with 16S rRNA gene data. Instead, Xanthobacter, Azorhizobium, Aquabacter, and Roseixanthobacter form a shuffled group, and reliable genus boundaries could not be drawn. This again highlights that genome-level comparisons are crucial for genus- and species-level resolution for this group of bacteria.

In the bac120 phylogenomic tree, all strains clustered within species boundaries defined by the GTDB-Tk (Figure 2, Supplementary Figures 1-2). We ran a separate average nucleotide identity analysis using FastANI (69) with a 95% species boundary threshold (72–74) and found that these species boundaries were consistent across the GTDB-Tk, ANI, and the phylogenomic trees (Supplementary Table 3). Additionally, a separate genome similarity metric used for informing species boundaries known as digital DNA-DNA hybridization (dDDH) via the Genome-to-Genome Distance Calculator (70,71) with a 70% threshold also aligned well with species-level boundaries as defined by ANI, with only five strains (23A, V8C-8, 126, SS-20, SS-22) falling slightly outside the dDDH species threshold (Supplementary Table 4). Given the consistency across this polyphasic approach, we used the species boundaries defined by the 95% ANI species-level threshold for our final classification.

We defined 10 novel Xanthobacter species (bringing the genus up to 18 species total) and 5 novel Roseixanthobacter gen. nov. species:

***X. agilis* Jenni and Aragno 1988** – Remains as previously described (11), a motile white/pale-yellow species with SA35^T^ as the type strain. No changes or updates in taxonomy.

***X. albus* sp. nov.** – A novel species containing some of our novel environmental isolates with V0C-6^T^ assigned as the type strain. The species is named for its white/pale-yellow appearance.

***X. aminoxidans* Doronina and Trotsenko 2003** – As previously described (10), a yellow species with 14a^T^ as the type strain. Additional repository strains and environmental isolates were assigned to the species.

***X. autotrophicus* (Baumgarten *et al.* 1974) Wiegel *et al.* 1978** – Remains as previously described (1), a yellow species with 7C^T^ as the type strain. Many canonical *X. autotrophicus* strains such as GZ29 and JW33 are verified as being true members, as well as the styrene-degrading strain 124X. Notably, the type strain *X. viscosus* 7d^T^ we obtained from DSMZ (DSM 21355^T^) was assigned as *X. autotrophicus* across all our analyses. See under *X. viscosus* for more details.

***X. cornucopiae* sp. nov**. – A novel species containing a single pale-yellow environmental isolate, V4C-4^T^. The species is named after the project from which it was discovered.

***X. dioxanivorans* Wang *et al.* 2021** – Remains as previously described (9), a yellow species with YN2^T^ as the type strain. No changes or updates in taxonomy.

***X. flavus* Malik and Claus 1979** – Remains as previously described (5), a yellow inducibly-motile species with 301^T^ as the type strain. Several additional repository strains and novel environmental isolates are assigned to this species. Notably, the very well-studied strain GJ10 is reclassified as *X. flavus*.

***X. lutulentifluminis* sp. nov.** – A novel species containing a single yellow novel environmental isolate, V3C-3^T^. The species is named after the Muddy River from which it was discovered.

***X. nonsaccharivorans* sp. nov.** – A novel species containing the singleton strain 14g^T^ (91,92) as the type strain. This species is named after the type strain’s notorious inability to grow on sugars in the past literature (92).

***X. oligotrophicus* Tikhonova *et al.* 2021** – Remains as previously described (7), a yellow species with 29k^T^ as the type strain. The species gains an additional repository strain.

***X. pseudotagetidis* sp. nov.** – A novel species containing the lone yellow type strain KA^T^, formerly known as and highly related to *X. tagetidis*.

***X. sediminis* sp. nov.** – A novel species consisting of pale-yellow environmental isolates, with V8C-5^T^ as the type strain. The species is named after sediment from which the type strain was isolated.

***X. tagetidis* Padden *et al.* 1997** – Remains as previously described (12), a yellow species with TagT2C^T^ as the type strain.

***X. toluenivorans* sp. nov.** – A novel species consisting of yellow toluene-degrading strains, with T101^T^ (31) assigned as the type strain.

***X. variabilis* sp. nov.** – A novel species consisting of pure-white environmental isolates, with V4C-8^T^ assigned as the type strain. The species lacks the carotenoid biosynthetic pathway. It is named for the strain-dependent growth properties of the species from its initial description in a dissertation (39).

***X. versatilis* sp. nov.** – A novel species containing the well-studied yellow strain Py2^T^ (formerly *X. autotrophicus*) and several novel environmental isolates. Due to its importance in the Xanthobacter literature (18,25–28,30,33,34), Py2^T^ was assigned as the type strain, and the name reflects the remarkable and varied substrate range that it can degrade and or grow on.

***X. viscosus* Doronina and Trotsenko 2003** – A non-motile yellow species of Xanthobacter (10). After sequencing the genome of the type strain *X. viscosus* 7d^T^ (which we obtained from the DSMZ as DSM 21355^T^), we found it was consistently assigned as *X. autotrophicus* across all our analyses. Its genome has a ∼99.99% ANI score to the *X. autotrophicus* 7C^T^ genome we sequenced in our study and to a previously sequenced 7C^T^ genome on NBCI (GCF_005871085.1), highly indicating that this strain belongs to *X. autotrophicus* (Supplementary Figure 6). A separate research group also previously published the *X. viscosus* 7d^T^ genome (JGI Gp0538788), which when analyzed with the same tests above yielded the exact same result. The previously published *X. viscosus* 7d^T^ 16S rRNA sequence (NR_025173.1, which was used in part to assign the strain as a new Xanthobacter species) does not match the 16S rRNA sequences extracted from either our 7d^T^ genome or the other released JGI 7d^T^ genome sequence (Supplementary Figure 7). There are 10 locations across the 7d^T^ 16S rRNA gene sequence from its original publication (NR_025173.1) that do not match any other highly related Xanthobacter (*X. autotrophicus*, *X. oligotrophicus*, *X. versatilis*, *X. toluenivorans*) nor any Xanthobacter type strain 16S rRNA gene at all (Supplementary Figure 7 and 8). Due to this high number of sites from the originally published 16S rRNA gene sequence that are not found anywhere across the genus Xanthobacter (and unlike any other Xanthobacter species), the originally published 7d^T^ 16S rRNA sequence raises concerns and should be revisited. While additional *X. viscosus* 7d^T^ repository sources (e.g., from VKM or KCTC) need to be sequenced and verified, it is likely that with the new genome sequences and 16S rRNA gene sequences available, this species should be reassigned to *X. autotrophicus*.

***X. wiegelii* sp. nov.** – A novel species containing the yellow strain RH 10^T^ (91) as the type strain. The species is named for Dr. Juergen Wiegel for his important and numerous contributions to the study of Xanthobacter.

***R. finlandensis* sp. nov.** – A novel species containing the pink-orangish strain VTT E-85241^T^ (93) as the type strain. The species is named for Finland from which it was isolated.

***R. glucoisosaccharinivorans* sp. nov.** – A novel species containing the pink-orangish strain VTT E-85242^T^ (93) as the type strain. The species is named for glucoisosaccharinic acid on which the VTT Roseixanthobacter strains were initially isolated.

***R. liquoris* sp. nov.** – A novel species containing the pink-orangish strain VTT E-85238^T^ (93) as the type strain. The species is named after the liquor contaminated pond from which it was isolated.

***R. pseudopolyaromaticivorans* sp. nov.** – A novel species consisting of several pink-orangish strains, with VTT E-85240^T^ (93) as the type strain. The species is named after its initial classification as the invalidly published “Xanthobacter polyaromaticivorans”.

***R. psychrophilus* sp. nov.** – A novel species containing the lone pink-orangish strain W30^T^ as the type strain. The species is named after its initial assignment of the type strain as a psychrophilic strain in its BacDive resource page (94).

Several other Xanthobacter strains consistently formed novel unnamed species clusters and branches in our analyses. However, these strains are not available from repositories, and hence they cannot be properly analyzed to obtain the necessary data to be classified as a novel species. Thus, there is likely known diversity yet to be characterized in the genus Xanthobacter.

### Physiological and phenotypic characterization of Xanthobacter and Roseixanthobacter species

With the newly defined species boundaries and assigned type strains, we then characterized the vast majority of the species based on a number of physiological parameters. This included color (Supplementary Figure 9), slime production (Table 3 and Supplementary Figure 9), cell size and morphology (Supplementary Figures 10 and 11 and Supplementary Table 5), motility (Supplementary Table 6 and 7), temperature ranges (Supplementary Table 8), antibiotic susceptibility (Supplementary Table 9), pH ranges (Supplementary Table 10), salinity ranges (Supplementary Table 11), autotrophic growth (Table 3), heterotrophic growth (Supplementary Table 12), carbon substrate usage (Supplementary Tables 13 and 14), and enzymatic properties (Supplementary Tables 15 and 16). A summary of phenotypic properties in Xanthobacter and Roseixanthobacter gen. nov. can be found in Table 3.

Overall, we found that phenotypic and physiological characteristics largely followed evolutionarily related branches as determined in the phylogenomic trees (Figure 1). Based on the evolutionary relatedness and shared characteristics, we found it helpful to split Xanthobacter species into clades that assist in comparing broader trends across the genus. All known Xanthobacter species are aerobic/microaerobic and capable of hydrogen-oxidizing, nitrogen-fixing, chemolithoautotrophic growth.

**Clade I** – Includes *X. versatilis* sp. nov., *X. autotrophicus*, *X. oligotrophicus*, and *X. toluenivorans* sp. nov.

**Clade IIA** – Includes the singletons *X. lutulentifluminis* sp. nov. and *X. nonsaccharivorans* sp. nov. These species are more closely related evolutionarily to Clade IIB Xanthobacter but have phenotypic differences, such as a lack of motility and different carbon substrate usage.

**Clade IIB** – Includes *X. aminoxidans*, *X. flavus*, and *X. wiegelii* sp. nov.

**Clade III** – Includes *X. cornucopiae* sp. nov., *X. agilis*, *X. sediminis* sp. nov., *X. albus* sp. nov., and *X. variabilis* sp. nov. Pigmentation is greatly diminished or downregulated in this clade leading to whiteish/pale-yellow color or is completely absent (as is the case for *X. variabilis* which is missing the carotenoid biosynthetic pathway).

**Clade IV** – Includes *X. tagetidis* and *X. pseudotagetidis*.

#### Color and Pigmentation

As for physical characteristics, the Xanthobacter are known for their yellow color (1,3), originating from the presence of zeaxanthin dirhamnoside in their membrane (95,96). Not all Xanthobacter are yellow, however (11) (Supplementary Figure 9). Clade III Xanthobacter (*X. agilis*, *X. albus* sp. nov., *X. sediminis* sp. nov., and *X. cornucopiae* sp. nov.) appear as whiteish/pale yellow and contain the same carotenoid operon for zeaxanthin production as the rest of the Xanthobacter. This lack of bright yellow color may stem from differences in regulation of the pathway, given zeaxanthin dirhamnoside was still identified as the responsible pigment (3,11), albeit at low concentrations in *X. agilis* compared to *X. autotrophicus* (3). However, *X. variabilis* is pure white in color and completely lacks the presence of carotenoid pathway genes in its genome that other Xanthobacter species have (e.g., *crtE, crtB, crtI, crtY,* and *crtZ*). This finding further raises questions on the utilization, reliance, and nature of the carotenoid pathway for Xanthobacter metabolism. The Roseixanthobacter gen. nov. species appear pink-orangish in color. In “*Xanthobacter polyaromaticivorans*”, a Roseixanthobacter-adjacent but invalidly published species, this pigment was identified as a zeaxanthin (41) but likely has different terminal substitutions (2). Images of dense growth and colonies from each of the type strains can be found in Supplementary Figure 9.

#### Slime Production

Xanthobacter species are also known for their ability to produce an exopolysaccharide slime, giving colonies the appearance of a “fried egg” on a plate (3). Slime production is apparent under almost all conditions in some species, (such as *X. versatilis* sp. nov. Py2^T^ as seen in this study), but is greatly increased during growth on media containing tricarboxylic acid cycle intermediates, such as succinate for others (1,3,86), like *X. autotrophicus* 7C^T^. Thus, we grew each of the type strains on rich media and minimal media containing succinate to observe slime production (Supplementary Figure 9). We found that Clade I, *X. lutulentifluminis* sp. nov. V3C-3^T^, Clade IIB, and *X. tagetidis* TagT2C^T^ produce visible amounts of slime when grown on succinate, however, Clade III Xanthobacter and Roseixanthobacter do not appear to produce any slime under these conditions tested. Visible amounts of slime are produced in *X. versatilis* sp. nov. Py2^T^ when grown on rich media (most noticeable when looking at single colonies) but significantly more slime is made when the strain is grown on succinate.

#### Cell Size and Morphology

Cell size and shape vary within Xanthobacter, as all species previously described are pleomorphic except *X. agilis* (1,5,7,9–12), although occasional branching has been noted for it, too (3). Typically rod-shaped, Xanthobacter species can also bend, twist, and may form coccoid structures (on alcohols), elongated structures, or branched structures (on tricarboxylic acid cycle intermediates). We imaged Xanthobacter and Roseixanthobacter gen. nov. grown on rich media and on minimal media containing succinate to observe morphological differences (Supplementary Figure 10). We found that Clade I, Clade IIA, and Clade IIB Xanthobacter shift from primarily rod-shaped morphologies to a branched cell morphology when grown on succinate alone. Clade III and Clade IV Xanthobacter do not undergo this striking shift, although branched cells can be occasionally seen under both growth conditions in these clades. Rather, Clade III Xanthobacter appear to become elongated when grown on succinate alone (and bent for *X. cornucopiae* sp. nov. V4C-4^T^). Roseixanthobacter gen. nov. species were largely rod-shaped or bent regardless of the media condition, however, similarly to Clade III and IV Xanthobacter, branched cells can be occasionally observed. We calculated cell sizes for the rod-shaped cells of Xanthobacter and Roseixanthobacter gen. nov. under the rich media growth condition and found that across both genera, cells were typically 1-2 µm long and 0.35-0.55 µm wide (Supplementary Table 5). Cell size dimensions across the evolutionary clades were similar among species within those clades (Supplementary Figure 11).

#### Motility

Xanthobacter species are motile through the expression of long peritrichous flagella, as described in studies on *X. agilis* and *X. flavus* (11,97). *X. flavus* 301^T^ was previously described as inducibly motile on some substrates (like alcohols, and inducibly produces long peritrichous flagella) compared to the constitutive motility observed in *X. agilis* SA35^T^ (97). Other Xanthobacter (*X. oligotrophicus*, *X. aminoxidans*, *X. autotrophicus*) were described as non-motile, although occasionally slime-free mutants of the *X. autotrophicus* type strain 7C^T^ (known as 7C SF) were inconsistently observed as motile (3,97). In their initial publications, *X. tagetidis* and *X. dioxanivorans* were also described to be motile (9,12); however, they are pictured with short projections unlike the long peritrichous flagella found in *X. agilis* and inducibly in *X. flavus*.

Given the range of results in the literature, first we checked for the presence of flagella proteins (84) within the genome annotations of type strains (*flg, fli, flh,* and *mot* genes). We found that Clade IIB (with the inducibly motile *X. flavus*), Clade III (with the constitutively motile *X. agilis*), and all Roseixanthobacter (except *R. psychrophilus* sp. nov. W30^T^) possess a large number of flagella genes in operonic units contained within an approximately 40 kb region of the genome (Supplementary Table 6). Notably, *X. tagetidis* and *X. dioxanivorans* lack this large operonic flagella region. However, all species (including those described as non-motile) contained scattered, non-operon copies of some flagella genes annotated as *fliO, fliJ, motA* (adjacent to a peptidoglycan-binding protein likely similar to *motB*) and in some species, *flgA*. These genes were isolated in different genome regions, often standalone, and other flagellar proteins were not found. It may be that the short protrusions observed in *X. tagetidis* and *X. dioxanivorans* are from these scattered genes or another unidentified pathway.

However, those Xanthobacter with observed long peritrichous flagella contain the large operonic flagella region in addition to those scattered non-operonic flagella proteins. Based on the previous literature on the inducible nature of motility in *X. flavus* (97), we tested for motility using a semisolid agar motility assay (81–83) under motility-inducing substrates (methanol, ethanol, propanol, isopropanol, gluconate) and non-motility-inducing growth substrates (succinate, fumarate). Our observations (Supplementary Table 7) indicate motility in semisolid agar appears to occur inducibly within Clade IIB (including *X. flavus*) and constitutively in Clade III (including *X. agilis*) of Xanthobacter. As for the Roseixanthobacter gen. nov. species, *R. finlandensis* sp. nov. VTT E-85241^T^ was constitutively motile and motility was inducible in all other Roseixanthobacter gen. nov. except *R. psychrophilus* sp. nov. W30^T^. Constitutively motile strains produced a strongly positive assay readout (deep red throughout the tube) after 5 days of incubation, while the inducibly motile strains needed 10–14 days of incubation for a weaker positive readout (light red throughout the tube).

Our bioinformatic confirmation of the large operonic flagella region aligns with our observations of motility in the semisolid agar assay (Supplementary Table 6). Notably, *X. tagetidis* TagT2C^T^ is missing this large operonic flagella region and was non-motile in the semisolid agar assay. It may be that the short protrusions described as flagella in Padden *et al.,* 1997 might be able to cause motility observed under light microscopy as previously described (12). However, these protrusions are not produced by operonic flagella genes. They are distinctly unlike the large peritrichous flagella observed in the inducibly motile *X. flavus* and constitutively motile *X. agilis* that did produce a positive result on the semisolid agar assay and that do contain large operonic flagella regions. Also of note, *X. aminoxidans* 14a^T^ and *X. wiegelii* RH 10^T^ were previously described as non-motile but were inducibly motile in semisolid agar under the conditions we tested and do contain large operonic flagella regions in their genome. We note that motility within Xanthobacter needs to be revisited and systematically tested across the genus. The expression of specific flagellar structures under different growth substrates and their potential impact on motility will likely need to be addressed to emend previous Xanthobacter species descriptions properly.

#### Temperature Range and Antibiotic Susceptibility

We tested for the temperature range over which the strains can grow by spot-plating cultures onto rich media plates incubated between 4°C to 42°C (Supplementary Table 8). All growth ranges and maxima were within expectations (3), with the Xanthobacter species growing well at 30°C and 37°C on agar plates (except for *X. toluenivorans* sp. nov. T101^T^), and Roseixanthobacter gen. nov. species at 25°C and 30°C. Antibiotic susceptibility was tested in a similar manner by spot-plating cultures onto rich media plates containing increasing amounts of gentamicin, kanamycin, streptomycin, and tetracycline (Supplementary Table 9). Antibiotic susceptibility appears to be strain-specific in the genus Xanthobacter, but we found a concentration that each type strain was susceptible to for antibiotic tested.

#### pH and Salinity Ranges

For finding pH and salinity ranges, we assayed for growth in liquid rich media containing 50 mM buffering agents at different pH values (pH 3–10) or different sodium chloride concentrations (0–6% w/v), respectively (Supplementary Tables 10 and 11). We find that Xanthobacter species typically prefer low salinity (≤ 1%) and near neutral pH (6–8) and that Roseixanthobacter gen. nov. species also prefer low salinity but a slightly more acidic pH (5–6).

#### Autotrophic and Heterotrophic Growth

A hallmark of the genus Xanthobacter is their ability to grow under hydrogen-oxidizing, nitrogen-fixing, chemolithoautotrophic conditions on minimal media and heterotrophically on rich media (1–3). We checked for autotrophic growth using the eVOLVER system (46) with a microaerobic chemolithoautotrophic gas mixture (∼3% H_2_, ∼10% CO_2_, ∼2% O_2_, ∼85% N_2_) bubbled through a sparger into vials containing inorganic minimal media lacking a nitrogen and carbon source. We found that all Xanthobacter species described can grow under these hydrogen-oxidizing, nitrogen-fixing, chemolithoautotrophic conditions (Table 3). Meanwhile, none of the Roseixanthobacter gen. nov. species could grow under this condition and their genomes lack the genes necessary for fixing carbon dioxide through the Calvin-Benson-Bassham Cycle. After adding succinate (0.5% w/v) to the vials, all Roseixanthobacter gen. nov. species were able to fix nitrogen under microaerobic conditions. All species tested were able to grow heterotrophically on 2xM1, M1, and R2A media (Supplementary Table 12). Almost all type strains preferred 2xM1, however, *X. toluenivorans* sp. nov. T101^T^, *X. agilis* SA35^T^, *X. tagetidis* TagT2C^T^, *R. liquoris* sp. nov. VTT E-85238^T^, and *R. psychrophilus* sp. nov. W30^T^ grew to a higher density in R2A.

#### Carbon Substrate Utilization

We first sought to characterize carbon substrate utilization across the two genera using a high-throughput liquid growth assay on minimal media containing 0.5% w/v or v/v of the substrate to be tested (Supplementary Table 13). Additionally, we provided a mixed carbon substrate condition, where 0.5% w/v of the substrate was combined with 0.01% succinate, a broadly used carbon source in the two genera observed in the first assay. This condition tested whether the presence of small amounts of a preferred carbon substrate could permit growth on other substrates that cannot be used alone. For *X. toluenivorans* sp. nov. T101^T^, we used 0.01% pyruvate instead of succinate due to its inability to grow to detectable OD_600_ values on succinate in liquid media.

We found that all Xanthobacter species, except *X. lutulentifluminis* sp. nov. V3C-3^T^, *X. pseudotagetidis* sp. nov. KA^T^, and *X. tagetidis* TagT2C^T^ can grow on alcohols alone, while only *R. finlandensis* sp. nov. VTT E-85241^T^ could do so from the Roseixanthobacter gen. nov. When provided a small amount of succinate, *X. tagetidis* TagT2C^T^ and *R. psychrophilus* sp. nov. W30^T^ were able to metabolize alcohols as well. As for amino acids, L-glutamine could be widely used by all Xanthobacter except *X. toluenivorans* sp. nov. T101^T^ and *X. pseudotagetidis* sp. nov. KA^T^, while usage of L-alanine followed along evolutionary lines with Clade IIB and Clade III Xanthobacter using the amino acid. β-hydroxybutyrate, pyruvate, and succinate were widely used substrates across all species tested except *X. toluenivorans* T101^T^. Some substrate usage followed along evolutionary clades, such as gluconate being used by all Xanthobacter except *X. toluenivorans* T101^T^ and Clade III Xanthobacter. Species like *X. autotrophicus* 7C^T^ and Clade IIB Xanthobacter grew exceptionally well on substrates like glycerol. Saccharide usage was extremely sparse across these two genera. Clade I Xanthobacter could utilize D-fructose while Clade IIB Xanthobacter could utilize D-mannose. Other saccharide usage was largely species-specific.

We also tested a subset of carbon substrates at different concentrations (1% w/v, 0.5% w/v, 0.1% w/v, 0.05% w/v) to identify concentration dependent growth effects. We found that for some substrates, like glyoxylate, there is a concentration-dependent growth effect with growth observed only at low concentrations—0.1% and 0.05% (Supplementary Table 14). Thus, for some of the substrates we tested above in the high-throughput carbon substrate assay at 0.5% w/v and saw no growth, concentrations may exist that permit growth. In summary, we find that some substrates could be used immediately by Xanthobacter and Roseixanthobacter gen. nov. species and others when additionally provided with small amounts of a preferred substrate. Metabolic preferences mostly followed along evolutionary lines, but utilization was sometimes species-specific.

#### Enzymatic Reactions

Finally, we also performed enzymatic and metabolic testing of all type strains using the API ZYM and API 20NE strip tests (Supplementary Tables 15 and 16. These strip tests are semi-quantitative assays contained within plastic cupules, often depicting a color change in response to an enzymatic or metabolic activity (85). We found that enzymatic activities were often genus-wide (e.g., acid phosphatase, trypsin, and cytochrome oxidase activity), within evolutionary clades (e.g., nitrate reduction to nitrites, cysteine arylamidase activity), or occasionally species- specific.

Overall, through our genome sequencing efforts and our environmental isolation campaign, we expand and revise the genus Xanthobacter and propose the novel genus Roseixanthobacter gen. nov. We investigated the evolutionary history and diversity of the genera, finding evolutionarily related clades, and we discovered a diverse array of phenotypes across the genus Xanthobacter that are often shared along evolutionary lines. With these findings and new genome sequences, it will help further the application of Xanthobacter biology and their unique biological properties that make them attractive in academic, industrial, and environmental settings.

Formal descriptions of each of the 10 novel Xanthobacter species, emendation of *X. aminoxidans*, and 5 novel Roseixanthobacter species are as follows.

### Description of *Xanthobacter albus* sp. nov

*Xanthobacter albus* (al’bus. L. masc. adj. *albus*, white, referring to the whiteish color of the species)

Cells are Gram-negative, pleomorphic, mostly rod-shaped (1.30 ± 0.34 μm in length and 0.49 ± 0.09 μm in width), with occasional branching cells, and can enlarge when grown on substrates such as succinate. Motile. Colonies are pale-yellow/whiteish, smooth, round, convex, with no observable slime produced under rich media or succinate.

Can grow under aerobic and microaerobic conditions. Able to grow heterotrophically and under hydrogen-oxidizing, nitrogen-fixing, chemolithoautotrophic conditions when supplied H_2_, N_2_, CO_2_, and microaerobic levels of O_2_. Growth with alcohols including methanol, ethanol, isopropanol, and propanol. Can grow on amino acids such as L-alanine, L-arginine, L-glutamine, L-histidine, L-lysine, L-proline, L-serine, and L-valine. Growth on other substrates such as acetate, β-hydroxybutyrate, butyrate, formate, fumarate, L-malate, propionate, pyruvate, glyoxylate, and succinate. Cannot grow on saccharides other than inconsistent minimal growth on sucrose.

Mesophilic and can grow between 15°C and 37°C, optimally between 30°C and 37°C. Neutrophilic and can grow between pH 3 to 8, optimally at pH 7. Growth observed between 0% to 4% w/v NaCl, optimally at 0.5% w/v NaCl.

The G+C content is 69.3% and its genome size is 4.5 Mb. The GenBank accession numbers for its genome sequence and its 16S rRNA gene sequence are JBAFWE000000000 and PP32888, respectively.

The type strain of *Xanthobacter albus* is represented by V0C-6^T^. The type strain was isolated in 2023 from an enriched autotrophic, nitrogen-fixing, hydrogen-oxidizing microbial community in minimal mineral media from a surface soil sample from Boston, Massachusetts, USA. It has been deposited in the Deutsche Sammlung von Mikroorganismen und Zellkulturen (DSMZ) under DSM (Under Review) and in the American Type Culture Collection under ATCC (Under Review).

### Description of Xanthobacter cornucopiae sp. nov

*Xanthobacter cornucopiae* (cor.nu.co.pi.ae. L. gen. n. *cornucopiae*, of the horn of plenty, referring to the Cornucopia project, during which the type strain was isolated)

Cells are Gram-negative, pleomorphic, mostly rod-shaped (1.64 ± 0.54 μm in length and 0.48 ± 0.18 μm in width), with occasional branching cells, and can enlarge and bend when grown on substrates such as succinate. Motile. Colonies are pale-yellow/whiteish, smooth, round, and convex on rich media and rough and uneven on substrates such as succinate. No observable slime produced on rich media or succinate.

Can grow under aerobic and microaerobic conditions. Able to grow heterotrophically and under hydrogen-oxidizing, nitrogen-fixing, chemolithoautotrophic conditions when supplied H_2_, N_2_, CO_2_, and microaerobic levels of O_2_. Growth with alcohols including methanol, ethanol, isopropanol, and propanol. Can grow on amino acids such as L-alanine, L-arginine, L-cysteine, L-glutamine, L-histidine, L-lysine, L-proline, and L-serine. Growth on other substrates such as acetate, β-hydroxybutyrate, butyrate, formate, fumarate, DL-lactate, D-malate, L-malate, propionate, pyruvate, glyoxylate, and succinate. Cannot grow on saccharides other than inconsistent minimal growth on ribose.

Mesophilic and can grow between 10°C and 37°C, optimally between 30°C and 37°C. Neutrophilic and can grow between pH 5 to 8, optimally at pH 8. Growth observed between 0% to 2% w/v NaCl, optimally at 0.5% w/v NaCl.

The G+C content is 69.2% and its genome size is 5.0 Mb. The GenBank accession numbers for its genome sequence and its 16S rRNA gene sequence are JBAFWM000000000 and PP328896, respectively.

The type strain of *Xanthobacter cornucopiae* is represented by V4C-4^T^. The type strain was isolated in 2023 from an enriched autotrophic, nitrogen-fixing, hydrogen-oxidizing microbial community in minimal mineral media from a root and sediment sample by the Muddy River in Boston, Massachusetts, USA. It has been deposited in the Deutsche Sammlung von Mikroorganismen und Zellkulturen (DSMZ) under DSM (Under Review) and in the American Type Culture Collection under ATCC (Under Review).

### Description of Xanthobacter lutulentifluminis sp. nov

*Xanthobacter lutulentifluminis* (lu.tu.len.ti.flu’mi.nis. L. masc. adj. *lutulentus*, muddy; L. neut. n. *flumen*, river; N.L. gen. n. *lutulentifluminis*, of the Muddy River, referring to where the type strain was isolated).

Cells are Gram-negative, pleomorphic, mostly rod-shaped (1.30 ± 0.36 μm in length and 0.41 ± 0.08 μm in width) but are heavily branched when grown on substrates such as succinate. Non-motile. Colonies are yellow, smooth, round, and convex on rich media with excessive slime produced when grown on succinate.

Can grow under aerobic and microaerobic conditions. Able to grow heterotrophically and under hydrogen-oxidizing, nitrogen-fixing, chemolithoautotrophic conditions when supplied H_2_, N_2_, CO_2_, and microaerobic levels of O_2_. Growth observed with isopropanol but not with other alcohols. Can consistently grow on amino acids such as L-glutamine and L-proline, inconsistently on L-isoleucine and L-threonine, and conditionally on L-cysteine. Growth on other substrates such as acetate, β- hydroxybutyrate, butyrate, formate, fumarate, gluconate, D-glucuronic acid, L-malate, propionate, pyruvate, glyoxylate, and succinate, and conditionally on glycerol and L-tartrate. Growth observed on L-sorbose and D-ribose, and conditionally on D-sorbitol.

Mesophilic and can grow between 15°C and 37°C, optimally between 30°C and 37°C. Neutrophilic and can grow between pH 5 to 8, optimally at pH 6. Growth observed between 0% to 5% w/v NaCl, optimally at 0% w/v NaCl.

The G+C content is 69.8% and its genome size is 4.8 Mb. The GenBank accession numbers for its genome sequence and its 16S rRNA gene sequence are JBAFVU000000000 and PP328893, respectively.

The type strain of *Xanthobacter lutulentifluminis* is represented by V3C-3^T^. The type strain was isolated in 2023 from an enriched autotrophic, nitrogen-fixing, hydrogen-oxidizing microbial community in minimal mineral media from a sediment sample by the Muddy River in Boston, Massachusetts, USA. It has been deposited in the Deutsche Sammlung von Mikroorganismen und Zellkulturen (DSMZ) under DSM (Under Review) and in the American Type Culture Collection under ATCC (Under Review).

### Description of Xanthobacter nonsaccharivorans sp. nov

*Xanthobacter nonsaccharivorans* (non.sac.cha.ri.vo’rans. L. prep. *non*, not; N.L. neut. n. *saccharum*, sugar; L. pres. part. *vorans*, devouring; N.L. part. adj. *nonsaccharivorans*, referring to the lack of ability of the type strain to grow on sugars)

Cells are Gram-negative, pleomorphic, mostly rod-shaped (1.42 ± 0.36 μm in length and 0.48 ± 0.09 μm in width) but are heavily branched when grown on substrates such as succinate. Non-motile. Colonies are yellow, smooth, round, and convex.

Can grow under aerobic and microaerobic conditions. Able to grow heterotrophically and under hydrogen-oxidizing, nitrogen-fixing, chemolithoautotrophic conditions when supplied H_2_, N_2_, CO_2_, and microaerobic levels of O_2_. Growth with alcohols including ethanol, isopropanol, and propanol. Can grow on amino acids such as L-alanine, L-cysteine, L-glutamine, L-histidine, and L-isoleucine, and conditionally on L-valine. Growth on other substrates such as acetate, β-hydroxybutyrate, butyrate, citrate, formate, fumarate, gluconate, glyoxylate, DL-lactate, D-malate, L-malate, pyruvate, and succinate. Cannot grow on saccharides other than inconsistent minimal growth seen on ribose.

Mesophilic and can grow between 15°C and 37°C, optimally between 30°C and 37°C. Neutrophilic and can grow between pH 5 to 8, optimally at pH 7. Growth observed between 0% to 5% w/v NaCl, optimally at 0% w/v NaCl.

The G+C content is 68.6% and its genome size is 5.3 Mb. The GenBank accession numbers for its genome sequence and its 16S rRNA gene sequence are JBAFVV000000000 and PP328870, respectively.

The type strain of *Xanthobacter nonsaccharivorans* is represented by 14g^T^ (=DSM 431^T^=JCM 1201^T^=CIP 105432^T^=IAM 12635^T^=NCIMB 10811^T^). The type strain was isolated from enrichment culture on hydrogen, carbon dioxide, and oxygen from a soil sample near Gottingen, Germany by Rudolph, 1968 initially as a coryneform bacterium, revised to *Corynebacterium autotrophicum*, and then to *Xanthobacter autotrophicus*.

### Description of Xanthobacter pseudotagetidis sp. nov

*Xanthobacter pseudotagetidis* (pseu.do.ta.ge’ti.dis. Gr. masc. adj. *pseudes*, false; N.L. gen. n. *tagetidis* (‘of Tagetes’), a specific epithet; N.L. gen. n. *pseudotagetidis*, a false (Xanthobacter) tagetidis)

Cells are Gram-negative, pleomorphic, mostly rod-shaped (1.22 ± 0.32 μm in length and 0.36 ± 0.09 μm in width) but are occasionally branched. Non-motile.Colonies are yellow, pinpoint, smooth, round, and convex. No observable slime is produced on rich media or on succinate.

Can grow under aerobic and microaerobic conditions. Able to grow heterotrophically and under hydrogen-oxidizing, nitrogen-fixing, chemolithoautotrophic conditions when supplied H_2_, N_2_, CO_2_, and microaerobic levels of O_2_. Growth not observed with alcohols at 0.5% v/v. Can conditionally grow on amino acids such as L-alanine, L-glutamine, L-proline, L-serine, L-threonine, and L-valine. Growth on other substrates such as acetate, β-hydroxybutyrate, butyrate, fumarate, gluconate, glyoxylate, L-malate, pyruvate, and succinate, and conditionally on D-glucuronic acid. Conditional growth observed on saccharides such as D-glucose, L-sorbose and D-ribose, and sucrose.

Mesophilic and can grow between 15°C and 42°C, optimally between 30°C and 37°C. Neutrophilic and can grow between pH 5 to 8, optimally at pH 8. Growth observed between 0% to 3% w/v NaCl, optimally at 0% w/v NaCl.

The G+C content is 69.7% and its genome size is 4.8 Mb. The GenBank accession numbers for its genome sequence and its 16S rRNA gene sequence are JBAFVY000000000 and PP328855, respectively.

The type strain of *Xanthobacter pseudotagetidis* is represented by KA^T^ (=DSM 11602^T^=KCTC 8467^T^). The type strain was isolated from sludge in Karlsruhe, Germany.

### Description of *Xanthobacter sediminis* sp. nov

*Xanthobacter sediminis* (se.di’mi.nis. L. gen. n. *sediminis*, of sediment, referring to the source from which the type strain was isolated)

Cells are Gram-negative, pleomorphic, mostly rod-shaped (1.54 ± 0.38 μm in length and 0.46 ± 0.08 μm in width) with occasional branching cells, and can enlarge and bend when grown on substrates such as succinate. Motile. Colonies are pale-yellow/whiteish, smooth, round, and convex on rich media and rough and uneven on substrates such as succinate. No observable slime produced on rich media or succinate.

Can grow under aerobic and microaerobic conditions. Able to grow heterotrophically and under hydrogen-oxidizing, nitrogen-fixing, chemolithoautotrophic conditions when supplied H_2_, N_2_, CO_2_, and microaerobic levels of O_2_. Growth with alcohols including methanol, ethanol, isopropanol, and propanol. Can grow on amino acids such as L-alanine, L-arginine, L-cysteine, L-glutamine, L-histidine, L-lysine, L-proline, L-serine, L-tryptophan, and L-valine. Growth on other substrates such as acetate, β-hydroxybutyrate, butyrate, formate, fumarate, glyoxylate, L-malate, propionate, pyruvate, and succinate. Some growth observed on saccharides such as D-cellobiose, L-rhamnose, and D-ribose.

Mesophilic and can grow between 15°C and 37°C, optimally between 30°C and 37°C. Neutrophilic and can grow between pH 3 to 8, optimally at pH 8. Growth observed between 0% to 6% w/v NaCl, optimally at 0% w/v NaCl.

The G+C content is 68.5% and its genome size is 4.7 Mb. The GenBank accession numbers for its genome sequence and its 16S rRNA gene sequence are JBAFWH000000000 and PP328903, respectively.

The type strain of *Xanthobacter sediminis* is represented by V8C-5^T^. The type strain was isolated in 2023 from an enriched autotrophic, nitrogen-fixing, hydrogen-oxidizing microbial community in minimal mineral media from a sediment sample taken by the Muddy River in Boston, Massachusetts, USA. It has been deposited in the Deutsche Sammlung von Mikroorganismen und Zellkulturen (DSMZ) under DSM (Under Review) and in the American Type Culture Collection under ATCC (Under Review).

### Description of Xanthobacter toluenivorans sp. nov

*Xanthobacter toluenivorans* (to.lu.e.ni.vo’rans. N.L. neut. n. *toluenum*, toluene; L. pres. part. *vorans*, devouring; N.L. part. adj. *toluenivorans*, toluene-devouring)

Cells are Gram-negative, pleomorphic, mostly rod-shaped (1.43 ± 0.50 μm in length and 0.51 ± 0.19 μm in width) but are largely branched when grown on substrates such as succinate. Non-motile. Colonies are yellow, pinpoint, smooth, round, translucent, and convex. No observable slime is produced on rich media.

Can grow under aerobic and microaerobic conditions. Able to grow heterotrophically and under hydrogen-oxidizing, nitrogen-fixing, chemolithoautotrophic conditions when supplied H_2_, N_2_, CO_2_, and microaerobic levels of O_2_. Growth observed with alcohols such as ethanol, isopropanol, and propanol. Can conditionally grow on amino acids such as L-alanine, L-cysteine, and L-serine. Growth on other substrates such as β-hydroxybutyrate, gluconate, glyoxylate, pyruvate, and inconsistently on succinate. No growth observed on saccharides. The type strain is known for its ability to degrade and grow on toluene.

Mesophilic and can grow between 10°C and 30°C, optimally at 30°C. Slightly acidophilic to neutrophilic, and can grow between pH 5 to 6, optimally at pH 6. Growth observed between 0% to 3% w/v NaCl, optimally at 0% w/v NaCl.

The G+C content is 67.6% and its genome size is 5.2 Mb. The GenBank accession numbers for its genome sequence and its 16S rRNA gene sequence are JBAFVT000000000 and PP328851, respectively.

The type strain of *Xanthobacter toluenivorans* is represented by T101^T^ (=ATCC 700551^T^). The type strain was isolated from a sample enriched on toluene from rock surface biomass in a toluene contaminated freshwater stream in a drainage ditch in Wilmington, Massachusetts, USA in 1992.

### Description of Xanthobacter variabilis sp. nov

*Xanthobacter variabilis* (va.ri.a’bi.lis. L. masc. adj. *variabilis*, variable, referring to the heterogenous, strain-dependent growth properties of the species)

Cells are Gram-negative, pleomorphic, mostly rod-shaped (1.35 ± 0.33 μm in length and 0.44 ± 0.09 μm in width) with occasional branching cells and can enlarge and bend when grown on substrates such as succinate. Motile. Colonies are white, smooth, round, and convex. No observable slime produced on rich media or succinate.

Can grow under aerobic and microaerobic conditions. Able to grow heterotrophically and under hydrogen-oxidizing, nitrogen-fixing, chemolithoautotrophic conditions when supplied H_2_, N_2_, CO_2_, and microaerobic levels of O_2_. Growth with alcohols including methanol, ethanol, isopropanol, and propanol. Can grow on amino acids such as L-alanine, L-arginine, L-cysteine, L-glutamine, L-histidine, L-lysine, L-proline, L-serine, L-threonine, L-tryptophan, and L-valine. Growth on other substrates such as acetate, β-hydroxybutyrate, butyrate, formate, fumarate, glycerol, L-malate, propionate, pyruvate, and succinate. The type strain is not observed to grow on saccharides except for D-melibiose.

Mesophilic and can grow between 15°C and 42°C, optimally between 30°C and 37°C. Neutrophilic and can grow between pH 4 to 8, optimally at pH 8. Growth observed between 0% to 6% w/v NaCl, optimally at 1% w/v NaCl.

The G+C content is 67.2% and its genome size is 4.6 Mb. The GenBank accession numbers for its genome sequence and its 16S rRNA gene sequence are JBAFWB000000000 and PP328898, respectively.

The type strain of *Xanthobacter variabilis* is represented by V4C-8^T^. The type strain was isolated in 2023 from an enriched autotrophic, nitrogen-fixing, hydrogen-oxidizing microbial community in minimal mineral media from a root and sediment sample taken by the Muddy River in Boston, Massachusetts, USA. It has been deposited in the Deutsche Sammlung von Mikroorganismen und Zellkulturen (DSMZ) under DSM (Under Review) and in the American Type Culture Collection under ATCC (Under Review).

### Description of Xanthobacter versatilis sp. nov

*Xanthobacter versatilis* (ver.sa’ti.lis. L. masc. adj. *versatilis*, referring to the versatile degradation properties of the species)

Cells are Gram-negative, pleomorphic, mostly rod-shaped (1.29 ± 0.38 μm in length and 0.47 ± 0.09 μm in width) but are heavily branched when grown on substrates such as succinate. Non-motile. Colonies are yellow, smooth, round, and convex on rich media. Slime is visible around colonies and growth on rich media however excessive amounts of slime are produced when grown on substrates such as succinate.

Can grow under aerobic and microaerobic conditions. Able to grow heterotrophically and under hydrogen-oxidizing, nitrogen-fixing, chemolithoautotrophic conditions when supplied H_2_, N_2_, CO_2_, and microaerobic levels of O_2_. Has a diverse growth substrate range. Growth observed with alcohols such as methanol, ethanol, isopropanol, and propanol. Can grow on amino acids such as L-cysteine, L-glutamine, L-histidine, L-isoleucine, L-phenylalanine, L-proline, and L-threonine, and conditionally on L-valine. Growth on other substrates such as acetate, β-hydroxybutyrate, butyrate, citrate, formate, fumarate, gluconate, D-glucuronic acid, glyoxylate, DL-lactate, D-malate, L-malate, pyruvate, and succinate, and conditionally on salicine. Growth observed on D-fructose, D-glucose, L-arabinose, D-sorbitol, conditionally on D-mannitol, and inconsistently on D-ribose. The type strain is known for its ability to grow on acetone and alkenes (such as ethene, propene, and butene).

Mesophilic and can grow between 10°C and 37°C, optimally between 30°C and 37°C. Neutrophilic and can grow between pH 5 to 8, optimally at pH 6. Growth observed between 0% to 3% w/v NaCl, optimally at 0% w/v NaCl.

The G+C content is 67.3% and its genome size is 5.6 Mb. The GenBank accession numbers for its genome sequence and its 16S rRNA gene sequence are GCF_000017645.1 and NR_074255.1, respectively.

The type strain of *Xanthobacter versatilis* is represented by Py2^T^ (=ATCC BAA- 1158^T^=DSM (Pending)). The type strain was isolated from enrichments with propene from freshwater in 1986.

### Description of *Xanthobacter wiegelii* sp. nov

*Xanthobacter wiegelii* (wie.gel’i.i. N.L. gen. n. *wiegelii*, of Juergen Wiegel, in recognition of his contributions to the study of Xanthobacter)

Cells are Gram-negative, pleomorphic, mostly rod-shaped (1.38 ± 0.41 μm in length and 0.43 ± 0.08 μm in width) but are heavily branched when grown on substrates such as succinate. Inducibly motile on alcohols such as methanol, ethanol, propanol, and isopropanol, and on substrates such as fumarate and gluconate. Colonies are yellow, waxy, smooth, round, and convex on rich media. Slime production is visible when grown on substrates such as succinate.

Can grow under aerobic and microaerobic conditions. Able to grow heterotrophically and under hydrogen-oxidizing, nitrogen-fixing, chemolithoautotrophic conditions when supplied H_2_, N_2_, CO_2_, and microaerobic levels of O_2_. Growth observed with alcohols such as methanol, ethanol, isopropanol, and propanol. Can grow on amino acids such as L-alanine, L-arginine, L-cysteine, L-glutamine, L-histidine, L-isoleucine, and L-proline. Growth on other substrates such as acetate, β-hydroxybutyrate, butyrate, citrate, fumarate, gluconate, D-glucuronic acid, glycerol, glyoxylate, DL-lactate, D-malate, L-malate, pyruvate, and succinate, and conditionally on propionate. Growth observed on saccharides such as D-mannose and inconsistently on D-glucose and D-ribose.

Mesophilic and can grow between 10°C and 37°C, optimally between 30°C and 37°C. Neutrophilic and can grow between pH 5 to 8, optimally at pH 6. Growth observed between 0% to 2% w/v NaCl, optimally at 0% w/v NaCl.

The G+C content is 67.7% and its genome size is 5.6 Mb. The GenBank accession numbers for its genome sequence and its 16S rRNA gene sequence are JBAFUY000000000 and PP328872, respectively.

The type strain of *Xanthobacter wiegelii* is represented by RH 10^T^ (=DSM 597^T^=JCM 7864^T^=CIP 105437^T^). The type strain was deposited in the Deutsche Sammlung von Mikroorganismen und Zellkulturen (DSMZ) originally in 1974.

### Emended Description of *Xanthobacter aminoxidans* Doronina and Trotsenko 2003

The description is as before in Doronina and Trotsenko 2003 (10) with the following change. Inducibly motile when grown on alcohols such as methanol, ethanol, and propanol and substrates such as gluconate.

### Description of Roseixanthobacter gen. nov

*Roseixanthobacter* (Ro.se.i.xan.tho.bac’ter. L. masc. adj. *roseus*, rose-colored; N.L. masc. n. *Xanthobacter*, a bacterial genus name; N.L. masc. n. *Roseixanthobacter*, rose-colored Xanthobacter)

Cells are Gram-negative, pleomorphic but mostly rod-shaped. Capable of aerobic and microaerobic growth. Mesophilic. Can fix dinitrogen gas under microaerobic conditions. Colonies are pink-orange in color. The type species is *Roseixanthobacter finlandensis*.

### Description of Roseixanthobacter finlandensis sp. nov

*Roseixanthobacter finlandensis* (fin.land.en’sis. N.L. masc. adj. *finlandensis*, of Finland, from which the type strain was isolated)

Cells are Gram-negative, pleomorphic, mostly rod-shaped (1.25 ± 0.48 μm in length and 0.39 ± 0.13 μm in width) but are occasionally branched. Motile. Colonies are pink/orange, smooth, round, and convex.

Can grow under aerobic and microaerobic conditions. Able to grow heterotrophically. Can fix dinitrogen gas as a nitrogen source under microaerobic levels of O_2_. Growth observed with alcohols such as methanol, ethanol, isopropanol, and propanol. Can grow on amino acids such as L-alanine, L-glutamine, L-proline, and L-serine and conditionally on L-cysteine and L-tryptophan. Growth on other substrates such as β-hydroxybutyrate, fumarate, gluconate, glyoxylate, DL-lactate, D-malate, L-malate, pyruvate, and succinate. Growth not observed on saccharides except for D-ribose. Can also degrade and grow on glucoisosaccharinic acid.

Mesophilic and can grow between 10°C and 37°C, optimally between 25°C and 30°C. Slightly acidophilic and neutrophilic and can grow between pH 5 to 8, optimally at pH 5. Growth observed between 0% to 2% w/v NaCl, optimally between 0% to 1% w/v NaCl.

The G+C content is 65.9% and its genome size is 5.6 Mb. The GenBank accession numbers for its genome sequence and its 16S rRNA gene sequence are JBAFWO000000000 and PP328909, respectively.

The type strain of *Roseixanthobacter finlandensis* is represented by VTT E- 85241^T^ (=DSM (Under Review)). The type strain was isolated and maintained by enrichment on glucoisosaccharinic acid in 1986 from a pond mildly contaminated with pulping wastes in Finland.

### Description of Roseixanthobacter glucoisosaccharinivorans sp. nov

*Roseixanthobacter glucoisosaccharinivorans* (glu.co.i.so.sac.cha.ri.ni.vo’rans. N.L. neut. n. *acidum glucoisosaccharinicum*, glucoisosaccharinic acid; L. pres. part. *vorans*, devouring; N.L. part. adj. *glucoisosaccharinivorans*, glucoisosaccharinic acid-devouring, eating)

Cells are Gram-negative, pleomorphic, mostly rod-shaped (1.71 ± 0.58 μm in length and 0.38 ± 0.10 μm in width) but are occasionally branched. Inducibly motile on substrates such as gluconate and fumarate. Colonies are pink/orange, smooth, round, and convex.

Can grow under aerobic and microaerobic conditions. Able to grow heterotrophically. Can fix dinitrogen gas as a nitrogen source under microaerobic levels of O_2_. Growth not observed with alcohols. Can grow on amino acids such as L-cysteine, L-glutamine, L-isoleucine, and L-phenylalanine, and conditionally on L-alanine, L-serine, and L-threonine. Growth on other substrates such as β-hydroxybutyrate, fumarate, gluconate, glyoxylate, L-malate, pyruvate, and succinate, conditionally on acetate, and inconsistently on citrate and D-tartrate. Growth not observed on saccharides. Can also degrade and grow on glucoisosaccharinic acid.

Mesophilic and can grow between 10°C and 37°C, optimally between 25°C and 30°C. Slightly acidophilic and neutrophilic and can grow between pH 3 to 8, optimally at pH 5. Growth observed between 0% to 3% w/v NaCl, optimally at 1% w/v NaCl.

The G+C content is 66.0% and its genome size is 5.4 Mb. The GenBank accession numbers for its genome sequence and its 16S rRNA gene sequence are JBAFWN000000000 and PP328910, respectively.

The type strain of *Roseixanthobacter glucoisosaccharinivorans* is represented by VTT E-85242^T^ (=DSM (Under Review)). The type strain was isolated and maintained by enrichment on glucoisosaccharinic acid in 1986 from the surface of waterlogged wood in a lake receiving wastes in Finland.

### Description of Roseixanthobacter liquoris sp. nov

*Roseixanthobacter liquoris* (li’quo.ris. L. gen. n. *liquoris*, of liquor, referring to the heavily liquor contaminated pond from which the type strain was isolated)

Cells are Gram-negative, pleomorphic, mostly rod-shaped (1.40 ± 0.60 μm in length and 0.41 ± 0.13 μm in width) but are occasionally branched. Inducibly motile on substrates such as gluconate. Colonies are pink/orange, smooth, round, and convex.

Can grow under aerobic and microaerobic conditions. Able to grow heterotrophically. Can fix dinitrogen gas as a nitrogen source under microaerobic levels of O_2_. Growth not observed with alcohols other than isopropanol. Can grow on amino acids such as L-cysteine, L-isoleucine, and L-phenylalanine, and conditionally on L-alanine, L-glutamine, L-proline, and L-serine. Growth on other substrates such as acetate, β-hydroxybutyrate, fumarate, gluconate, glyoxylate, L-malate, pyruvate, and succinate, and D-tartrate, and conditionally on butyrate. Growth not observed on saccharides except conditionally on L-xylose and D-ribose. Can also degrade and grow on glucoisosaccharinic acid.

Mesophilic and can grow between 10°C and 37°C, optimally between 25°C and 30°C. Slightly acidophilic and neutrophilic and can grow between pH 3 to 7, optimally at pH 5. Growth observed between 0% to 1.5% w/v NaCl, optimally at 0.5% w/v NaCl.

The G+C content is 67.2% and its genome size is 5.0 Mb. The GenBank accession numbers for its genome sequence and its 16S rRNA gene sequence are JBAFWP000000000 and PP328906, respectively.

The type strain of *Roseixanthobacter liquoris* is represented by VTT E-85238^T^ (=DSM (Under Review)). The type strain was isolated and maintained by enrichment on glucoisosaccharinic acid in 1986 from a pond heavily contaminated with black liquor on the grounds of a puling mill in Finland.

### Description of Roseixanthobacter pseudopolyaromaticivorans sp. nov

*Roseixanthobacter pseudopolyaromaticivorans* (pseu.do.po.ly.a.ro.ma.ti.ci.vo’rans. Gr. masc. adj. *pseudes*, false; Gr. masc. adj. *polys*, many; L. masc. adj. *aromaticus*, aromatic; L. pres. part. *vorans*, devouring; N.L. part. adj. *pseudopolyaromaticivorans*, a false polyaromaticivorans, referring to the species’ initial assessment as a *Xanthobacter polyaromaticivorans*)

Cells are Gram-negative, pleomorphic, mostly rod-shaped (1.27 ± 0.41 μm in length and 0.36 ± 0.07 μm in width) but are occasionally branched. Inducibly motile on substrates such as gluconate and fumarate. Colonies are pink/orange, smooth, round, and convex.

Can grow under aerobic and microaerobic conditions. Able to grow heterotrophically. Can fix dinitrogen gas as a nitrogen source under microaerobic levels of O_2_. Growth not observed with alcohols. Can grow on amino acids such as L-glutamine, L-isoleucine, and L-phenylalanine. Growth on other substrates such as acetate, β-hydroxybutyrate, fumarate, gluconate, glyoxylate, L-malate, pyruvate, and succinate. Growth not observed on saccharides except conditionally on L-sorbose and D-ribose. Can also degrade and grow on glucoisosaccharinic acid.

Mesophilic and can grow between 10°C and 37°C, optimally between 25°C and 30°C. Neutrophilic and can grow between pH 5 to 7, optimally at pH 6. Growth observed between 0% to 1.5% w/v NaCl, optimally at 0 to 0.5% w/v NaCl.

The G+C content is 65.7% and its genome size is 5.5 Mb. The GenBank accession numbers for its genome sequence and its 16S rRNA gene sequence are JBAFWQ000000000 and PP328908, respectively.

The type strain of *Roseixanthobacter pseudopolyaromaticivorans* is represented by VTT E-85240^T^ (=DSM (Under Review)). The type strain was isolated and maintained by enrichment on glucoisosaccharinic acid in 1986 from a stream contaminated with pulping wastes in Finland.

### Description of Roseixanthobacter psychrophilus sp. nov

*Roseixanthobacter psychrophilus* (psy.chro’phi.lus. Gr. masc. adj. *psychros*, cold; Gr. mac. adj. *philos*, loving; N.L. masc. adj. *psychrophilus*, cold-loving)

Cells are Gram-negative, pleomorphic, mostly rod-shaped (1.30 ± 0.38 μm in length and 0.42 ± 0.09 μm in width) but are occasionally branched. Non-motile. Colonies are pink/orange, rough, round, and convex.

Can grow under aerobic and microaerobic conditions. Able to grow heterotrophically. Can fix dinitrogen gas as a nitrogen source under microaerobic levels of O_2_. Growth not observed with alcohols unless grown with a small amount of a substrate such as succinate. Can grow on amino acids such as L-glutamine and L-isoleucine, and conditionally on L-alanine, L-arginine, L-cysteine, L-histidine, L-lysine, and L-ornithine. Growth on other substrates such as glyoxylate, L-malate, pyruvate, and succinate, and conditionally on acetate, β-hydroxybutyrate, butyrate, formate, fumarate, D-glucuronic acid, L-tartrate, and propionate. Conditional growth on saccharides such as D-mannose, L-sorbose, D-ribose, and sucrose.

Mesophilic/slightly psychrophilic and can grow between 10°C and 30°C, well at 20°C, optimally between 25°C and 30°C. Slightly acidophilic and neutrophilic and can grow between pH 3 to 8, optimally at pH 5. Growth observed between 0% to 2% w/v NaCl, optimally at 0.5% w/v NaCl.

The G+C content is 65.3% and its genome size is 6.0 Mb. The GenBank accession numbers for its genome sequence and its 16S rRNA gene sequence are JBAFWT000000000 and PP328867, respectively.

The type strain of *Roseixanthobacter psychrophilus* is represented by W30^T^ (=DSM 24535^T^=KCTC 8466^T^). The type strain was isolated from soil in Welsberg, South Tyrol, Italy in 2008.

## ACKNOWLEDGEMENTS

We would like to thank Dr. Aharon Oren for the assistance with the bacterial etymology. We would also like to thank Dr. Scott Ensign and Dr. Andrew Crombie for their help with the deposition of strain Py2. Finally, we would like to thank Dr. Vera Thiel and Dr. Irina Tsitko for their assistance with type strain deposition.

