## Supplemental for "Expansion and revision of the genus Xanthobacter and proposal of Roseixanthobacter gen. nov."

### Supplementary Table 1 – Carbon substrate sources for physiological condition testing in this study.

| Substrate type | Substrate | Product Number |
|---|---|---|
| Sugars | Cellobiose (D-(+)-Cellobiose) | Sigma Aldrich C7252-100G |
| | D-Fructose (D-(-)-Fructose) | Sigma Aldrich F0127-500G |
| | D-Glucose (D-(+)-Glucose) | Sigma Aldrich G7021-100G |
| | D-Mannose (D-(+)-Mannose) | Sigma Aldrich M6020-100G |
| | D-Raffinose (D-(+)-Raffinose Pentahydrate) | Sigma Aldrich R0250-1KG |
| | D-Xylose (D-(+)-Xylose) | Sigma Aldrich X1500-500G |
| | Galactose (D-(+)-Galactose) | Sigma Aldrich G0625-1KG |
| | Inositol (Myo-Inositol) | Sigma Aldrich I5125-50G |
| | Lactose (D-(+)-Lactose Monohydrate) | JT Baker 2248-01 |
| | L-Arabinose | GoldBio A-300-100 |
| | L-Sorbose | Thermo Scientific Chemicals AC225961000 |
| | L-Xylose | Thermo Scientific Chemicals B21622.14 |
| | Maltose (D-(+)-Maltose Monohydrate) | Sigma Aldrich M5895-500G |
| | Mannitol (D-Mannitol) | Sigma Aldrich M4125-500G |
| | Melibiose (D-Melibiose Monohydrate) | GoldBio M-150-50 |
| | Rhamnose (L-Rhamnose Monohydrate) | GoldBio R-105-50 |
| | Ribose (D-(-)-Ribose) | Sigma Aldrich R7500-100G |
| | Sorbitol (D-Sorbitol) | Sigma Aldrich S1876-1KG |
| | Sucrose (D-Sucrose) | Sigma Aldrich S0389-1KG |
| | Trehalose (D-Trehalose) | Thermo Scientific Chemicals 309871000 |
| Metabolites | 2-Oxogluturate (2-Oxoglutaric Acid) | TCI America K0005100G |
| | Acetate (Potassium Acetate) | Sigma Aldrich P1147-1KG |
| | Alpha-Aminobutyrate (DL-2-Aminobutyric Acid) | Thermo Scientific Chemicals L06035.14 |
| | Beta-Hydroxybutryate (DL-3-Hydroxybutyric Acid Sodium Salt) | Thermo Scientific Chemicals 215010250 |
| | Butyrate (Sodium Butyrate) | Thermo Scientific Chemicals A11079.22 |
| | Citrate (Sodium Citrate Tribasic Dihydrate) | Sigma Aldrich C3434-1KG |
| | DL-Lactate (Sodium DL-Lactate) | Thermo Scientific Chemicals 041529.AK |
| | D-Malate (D-(+)-Malic Acid) | Thermo Scientific Chemicals 153500250 |
| | D-Mandelate (D-(-)-Mandelic Acid) | TCI America M0662100G |
| | D-Tartrate (D-(-)-Tartaric Acid) | Sigma Aldrich T206-100G |
| | Formate (Sodium Formate) | Sigma Aldrich 71541-250G |
| | Fumarate (Sodium Fumarate Dibasic) | Sigma Aldrich F1506-100G |
| | Gluconate (Sodium Gluconate) | Sigma Aldrich S2054-100G |
| | Glucuronic acid (D-Glucuronic Acid Sodium Salt) | Sigma Aldrich G8645-25G |
| | Glutarate (Glutaric Acid) | Thermo Scientific Chemicals A14595.22 |
| | Glyoxylate (Glyoxylic Acid Monohydrate) | Thermo Scientific Chemicals A16058.22 |
| | L-Malate (Sodium L-Malate) | MP Biomedicals 0521807310 |
| | L-Mandelate (L-(+)-Mandelic Acid) | TCI America M066125G |
| | L-Tartrate (Sodium L-(+)-Tartrate Dihydrate) | Thermo Scientific Chemicals A16187.30 |
| | Malonate (Malonic Acid) | Thermo Scientific Chemicals A11526.22 |
| | N-Acetyl-Glucosamine | Sigma Aldrich A3286-25G |
| | Oxalate (Sodium Oxalate) | Sigma Aldrich O0136-100G |
| | Propionate (Sodium Proprionate) | Sigma Aldrich P1880-100G |
| | Pyruvate (Sodium Pyruvate) | Alfa Aesar A11148 |
| | Succinate (Di-sodium Succinate) ***Batch-to-batch variability | Sigma Aldrich 8.18601.0500 |
| Alcohols | Ethanol | Koptec 200 Proof Ethanol V1001 |
| | Isopropanol | VWR 0918-1L |
| | Methanol | Thermo Scientific Chemicals 268280010 |
| | Propanol (1-Propanol) | Thermo Scientific Chemicals 445775000 |
| Other | Benzoate (Sodium Benzoate) | Thermo Scientific Chemicals 447802500 |
| | Glycerol | Thermo Scientific Chemicals 17904 |
| | Salicine (D-(-)-Salicin) | Thermo Scientific Chemicals 132590250 |
| | Urea | Sigma Aldrich U1250-1KG |
| Amino acids | L-Alanine | Sigma Aldrich A7627-1G |
| | L-Arginine (Monohydrochloride) | Sigma Aldrich A5131-1G |
| | L-Aspartate | Sigma Aldrich A9256-1G |
| | L-Cysteine | Sigma Aldrich C-7352 |
| | L-Glutamine | Sigma Aldrich G3126-1G |
| | L-Histidine (Monohydrochloride) | Sigma Aldrich H8125-1G |
| | L-Isoleucine | Sigma Aldrich I2752-1G |
| | L-Leucine | Sigma Aldrich L8000-1G |
| | L-Lysine (Monohydrochloride) | Sigma Aldrich L5626-1G |
| | L-Methionine | Sigma Aldrich M9625-1G |
| | L-Ornithine (Monohydrochloride) | Sigma Aldrich O2375-25G |
| | L-Phenylalanine | Sigma Aldrich P2126-1G |
| | L-Proline | Sigma Aldrich P0380-1G |
| | L-Serine | Sigma Aldrich S4500-1G |
| | L-Threonine | Sigma Aldrich T8625-1G |
| | L-Tryptophan | Sigma Aldrich T0254-1G |
| | L-Valine | Sigma Aldrich V0500-1G |

**Supplementary Table 2 – Genome accession numbers and sequencing statistics for all newly sequenced genomes and prior sequenced genomes used for bioinformatic work in this study.** Accessions in bold under “Other strain identifiers” are the repository we obtained the strain from. Strains without a value under “Prior designation” are strains we isolated ourselves from the environment.

| Strain designation | Prior designation | Sequenced in this study | Other strain identifiers | Genome accession | 16S rRNA accession | Genome GC content | N50 | Contigs | CDS | tRNAs | rRNAs | Genome size | Median contig coverage |
|---|---|---|---|---|---|---|---|---|---|---|---|---|---|
| Xanthobacter agilis MA37 | Xanthobacter agilis MA37 | X | **ATCC 43848**; NEU 2067 | JBAFWL000000000 | PP328848 | 67.3% | 214.9 Kb | 77 | 4124 | 49 | 9 | 4.7 Mb | 185x |
| Xanthobacter agilis MA40 | Xanthobacter agilis MA40 | X | **ATCC 43849**; NEU 2069 | JBAFWK000000000 | PP328849 | 67.3% | 214.9 Kb | 74 | 4125 | 49 | 9 | 4.7 Mb | 203x |
| Xanthobacter agilis SA35 | Xanthobacter agilis SA35 | X | ATCC 43847; **DSM 3770**; LMG 16336; VKM B-2105; LMG 7994; NCIMB 12683; NEU 2015; NCAIM B.01949 | JBAFWJ000000000 | PP328869 | 67.4% | 240.7 Kb | 63 | 4290 | 48 | 9 | 4.8 Mb | 242x |
| Xanthobacter albus V0B-10 | - | X | DSM | JBAFWC000000000 | PP328879 | 69.3% | 209.5 Kb | 69 | 3953 | 51 | 12 | 4.5 Mb | 235x |
| Xanthobacter albus V0C-6 | - | X | ATCC; DSM | JBAFWE000000000 | PP328881 | 69.3% | 275.8 Kb | 61 | 3955 | 52 | 12 | 4.5 Mb | 159x |
| Xanthobacter albus V13C-5 | - | X | DSM | JBAFWD000000000 | PP328883 | 69.3% | 368 Kb | 65 | 3955 | 45 | 3 | 4.5 Mb | 205x |
| Xanthobacter albus V2C-4 | - | X | DSM | JBAFWG000000000 | PP328888 | 69.1% | 204 Kb | 63 | 4073 | 49 | 9 | 4.6 Mb | 187x |
| Xanthobacter albus V2C-8 | - | X | DSM | JBAFWF000000000 | PP328889 | 69.0% | 247.5 Kb | 82 | 4224 | 49 | 9 | 4.7 Mb | 180x |
| Xanthobacter aminoxidans 14a | Xanthobacter aminoxidans 14a | X | ATCC BAA-299; CIP 108461; **DSM 15009**; KCTC 12307; VKM B-2254 | JBAFUV000000000 | PP328862 | 67.9% | 241.9 Kb | 95 | 5468 | 44 | 3 | 5.8 Mb | 199x |
| Xanthobacter aminoxidans CB3 | Xanthobacter sp. CB3 | X | **DSM 14518** | JBAFUS000000000 | PP328858 | 68.0% | 1.3 Mb | 16 | 5138 | 47 | 3 | 5.5 Mb | 177x |
| Xanthobacter aminoxidans CB5 | Xanthobacter sp. CB5 | X | **DSM 14519** | JBAFUR000000000 | PP328859 | 68.0% | 758.4 Kb | 18 | 5137 | 47 | 3 | 5.5 Mb | 146x |
| Xanthobacter aminoxidans V7C-9 | - | X | DSM | JBAFUT000000000 | PP328902 | 67.9% | 352.3 Kb | 52 | 5291 | 47 | 3 | 5.7 Mb | 162x |
| Xanthobacter aminoxidans V8C-8 | - | X | DSM | JBAFUU000000000 | PP328904 | 67.9% | 649.6 Kb | 22 | 4675 | 46 | 3 | 5.0 Mb | 175x |
| Xanthobacter autotrophicus 124X | Xanthobacter sp. 124X | X | ATCC 49450; **DSM 6696** | JBAFVO000000000 | PP328873 | 67.1% | 260.5 Kb | 122 | 5043 | 48 | 3 | 5.5 Mb | 204x |
| Xanthobacter autotrophicus 19/-/x | Xanthobacter autotrophicus 19/-/x | X | **DSM 2009** | JBAFVR000000000 | PP328864 | 67.7% | 338.3 Kb | 49 | 4347 | 47 | 3 | 4.8 Mb | 220x |
| Xanthobacter autotrophicus 7C | Xanthobacter autotrophicus 7C | X | ATCC 35674; BCRC 12235; CCRC 12235; CIP 105431; **DSM 432**; IAM 12579; IAM 12636; JCM 1202; LMG 7043; NBRC 102463; NCAIM B0.1945; NCIB 10809; NCIMB 10809; NRRL B-14836 | JBAFVK000000000 | PP328871 | 67.6% | 217.6 Kb | 77 | 4473 | 46 | 3 | 5.0 Mb | 59x |
| Xanthobacter autotrophicus 7C SF | Xanthobacter autotrophicus 7C SF | X | **DSM 2267** | JBAFVJ000000000 | PP328866 | 67.6% | 327.5 Kb | 67 | 4477 | 46 | 3 | 5.0 Mb | 64x |
| Xanthobacter viscosus 7d (*classified as Xanthobacter autotrophicus in this study) | Xanthobacter viscosus 7d | X | ATCC BAA-298; CIP 108462; **DSM 21355**; KCTC 12306; VKM B-2253 | JBAFVI000000000 & Gp0538788 (JGI) | PP328865 | 67.6% | 381.1 Kb | 56 | 4476 | 46 | 3 | 5.0 Mb | 197x |

| | | | | | | | | | | | | | |
|---|---|---|---|---|---|---|---|---|---|---|---|---|---|
| Xanthobacter autotrophicus CCUG 44692 | Xanthobacter autotrophicus CCUG 44692 | X | **CCUG 44692** | JBAFVL000000000 | PP328853 | 67.5% | 286.6 Kb | 80 | 4699 | 48 | 3 | 5.2 Mb | 193x |
| Xanthobacter autotrophicus GZ29 | Xanthobacter autotrophicus GZ29 | X | **DSM 1393**; LMG 7044 | JBAFVQ000000000 | PP328856 | 67.6% | 227.5 Kb | 65 | 4650 | 46 | 4 | 5.1 Mb | 57x |
| Xanthobacter autotrophicus JW33 | Xanthobacter autotrophicus JW33 | X | **DSM 1618**; IFO 14758; NBRC 14758; NCIMB 12468 | JBAFVN000000000 | PP328863 | 67.3% | 197.9 Kb | 152 | 5017 | 48 | 3 | 5.5 Mb | 56x |
| Xanthobacter autotrophicus NCIMB 11171 | Xanthobacter autotrophicus NCIMB 11171 | X | LMD 90.18; NCCB 90018; **NCIMB 11171** | JBAFVM000000000 | PP328877 | 67.5% | 323 Kb | 100 | 4739 | 47 | 3 | 5.2 Mb | 236x |
| Xanthobacter autotrophicus V0C-4 | - | X | DSM | JBAFVP000000000 | PP328880 | 67.2% | 364.6 Kb | 107 | 5093 | 50 | 4 | 5.6 Mb | 122x |
| Xanthobacter cornucopiae V4C-4 | - | X | ATCC; DSM | JBAFWM000000000 | PP328896 | 69.2% | 280.2 Kb | 91 | 4326 | 46 | 3 | 5.0 Mb | 205x |
| Xanthobacter flavus 301 | Xanthobacter flavus 301 | X | ATCC 35867; BCRC 12271; CCM 4469; CCRC 12271; CIP 105434; **DSM 338**; IFO 14759; JCM 1204; LMG 7045; NBRC 14759; NCAIM B.01946; NCIB 10071; NCIMB 10071; NRRL B-14838; VKM B-2106 | JBAFUM000000000 | PP328868 | 67.9% | 505.5 Kb | 57 | 5387 | 46 | 4 | 5.8 Mb | 142x |
| Xanthobacter flavus R-10 | Xanthobacter sp. R-10 | X | **DSM 14521** | JBAFUK000000000 | PP328861 | 67.8% | 363 Kb | 70 | 5505 | 48 | 3 | 5.8 Mb | 176x |
| Xanthobacter flavus SS-20 | Xanthobacter sp. SS-20 | X | IFO 15493; **NBRC 15493** | JBAFUQ000000000 | PP328875 | 68.0% | 868.2 Kb | 22 | 4951 | 45 | 3 | 5.4 Mb | 159x |
| Xanthobacter flavus SS-22 | Xanthobacter sp. SS-22 | X | IFO 15494; **NRBC 15494** | JBAFUP000000000 | PP328876 | 68.0% | 784.2 Kb | 14 | 4918 | 46 | 3 | 5.4 Mb | 164x |
| Xanthobacter flavus V2C-3 | - | X | DSM | JBAFUN000000000 | PP328887 | 67.8% | 273.5 Kb | 74 | 5451 | 45 | 3 | 5.9 Mb | 158x |
| Xanthobacter flavus V4C-10 | - | X | DSM | JBAFUJ000000000 | PP328895 | 67.9% | 450.9 Kb | 40 | 5291 | 46 | 3 | 5.7 Mb | 116x |
| Xanthobacter flavus V4C-7 | - | X | DSM | JBAFUL000000000 | PP328897 | 68.1% | 393.1 Kb | 47 | 4896 | 46 | 3 | 5.3 Mb | 117x |
| Xanthobacter flavus V4C-9 | - | X | DSM | JBAFUO000000000 | PP328899 | 67.8% | 283.3 Kb | 59 | 5476 | 46 | 3 | 5.8 Mb | 121x |
| Xanthobacter lutulentifluminis V3C-3 | - | X | ATCC; DSM | JBAFVU000000000 | PP328893 | 69.8% | 421.9 Kb | 41 | 4398 | 49 | 3 | 4.8 Mb | 165x |
| Xanthobacter nonsaccharivorans 14g | Xanthobacter autotrophicus 14g | X | CIP 105432; **DSM 431**; IAM 12635; JCM 1201; NCIB 10811; NCIMB 10811 | JBAFVV000000000 | PP328870 | 68.6% | 652.7 Kb | 18 | 4824 | 46 | 3 | 5.3 Mb | 56x |
| Xanthobacter oligotrophicus 23A | Xanthobacter autotrophicus 23A | X | CIP 105433; **DSM 685**; JCM 1203 | JBAFVH000000000 | PP328874 | 68.1% | 374.5 Kb | 56 | 4753 | 46 | 3 | 5.2 Mb | 55x |
| Xanthobacter pseudotagetidis KA | Xanthobacter tagetidis KA | X | **DSM 11602;** KCTC | JBAFVY000000000 | PP328855 | 69.7% | 254.8 Kb | 41 | 4303 | 47 | 3 | 4.8 Mb | 240x |
| Xanthobacter sediminis V3B-7B | - | X | DSM | JBAFWI000000000 | PP328891 | 68.5% | 181.6 Kb | 87 | 4188 | 43 | 3 | 4.7 Mb | 210x |
| Xanthobacter sediminis V8C-5 | - | X | ATCC; DSM | JBAFWH000000000 | PP328903 | 68.5% | 286.9 Kb | 73 | 4195 | 46 | 3 | 4.7 Mb | 274x |
| Xanthobacter tagetidis A2 | Xanthobacter tagetidis A2 | X | **ATCC 700315** | JBAFVX000000000 | PP328850 | 69.6% | 251.5 Kb | 42 | 4481 | 46 | 3 | 4.9 Mb | 172x |
| Xanthobacter tagetidis TagT2C | Xanthobacter tagetidis TagT2C | X | ATCC 700314; **DSM 11105**; NCIMB 13547 | JBAFVW000000000 | PP328854 | 69.6% | 272.8 Kb | 44 | 4483 | 46 | 3 | 4.9 Mb | 195x |

| | | | | | | | | | | | | | |
|---|---|---|---|---|---|---|---|---|---|---|---|---|---|
| Xanthobacter toluenivorans T101 | Xanthobacter autotrophicus T101 | X | **ATCC 700551** | JBAFVT000000000 | PP328851 | 67.6% | 118.1 Kb | 214 | 4818 | 47 | 3 | 5.2 Mb | 157x |
| Xanthobacter toluenivorans T102 | Xanthobacter autotrophicus T102 | X | **ATCC 700552** | JBAFVS000000000 | PP328852 | 67.6% | 176.7 Kb | 127 | 4977 | 49 | 3 | 5.4 Mb | 181x |
| Xanthobacter variabilis V13B-9B | - | X | DSM | JBAFVZ000000000 | PP328882 | 67.1% | 372.8 Kb | 38 | 3960 | 51 | 9 | 4.7 Mb | 239x |
| Xanthobacter variabilis V13C-7B | - | X | DSM | JBAFWA000000000 | PP328884 | 67.1% | 295.9 Kb | 39 | 4023 | 47 | 3 | 4.7 Mb | 139x |
| Xanthobacter variabilis V4C-8 | - | X | ATCC; DSM | JBAFWB000000000 | PP328898 | 67.2% | 681.3 Kb | 17 | 3867 | 47 | 3 | 4.6 Mb | 189x |
| Xanthobacter versatilis CB6 | Xanthobacter sp. CB6 | X | **DSM 14520** | JBAFVA000000000 | PP328860 | 67.8% | 433.6 Kb | 57 | 4638 | 50 | 3 | 5.1 Mb | 183x |
| Xanthobacter versatilis V1C-2 | - | X | DSM | JBAFVD000000000 | PP328885 | 67.9% | 551.8 Kb | 38 | 4292 | 47 | 3 | 4.8 Mb | 150x |
| Xanthobacter versatilis V1C-6B | - | X | DSM | JBAFVC000000000 | PP328886 | 67.7% | 451.9 Kb | 58 | 4587 | 47 | 3 | 5.1 Mb | 276x |
| Xanthobacter versatilis V2C-9 | - | X | DSM | JBAFVB000000000 | PP328890 | 67.8% | 343.2 Kb | 45 | 4396 | 48 | 5 | 4.9 Mb | 202x |
| Xanthobacter versatilis V3C-2 | - | X | DSM | JBAFVG000000000 | PP328892 | 67.6% | 224.8 Kb | 100 | 4861 | 46 | 3 | 5.3 Mb | 122x |
| Xanthobacter versatilis V3C-4 | - | X | DSM | JBAFVF000000000 | PP328894 | 67.6% | 324.9 Kb | 92 | 4739 | 47 | 3 | 5.2 Mb | 182x |
| Xanthobacter versatilis V7C-1B | - | X | DSM | JBAFVE000000000 | PP328900 | 67.8% | 449.2 Kb | 46 | 4525 | 47 | 3 | 5.0 Mb | 184x |
| Xanthobacter versatilis V7C-4 | - | X | DSM | JBAFUZ000000000 | PP328901 | 67.8% | 554.7 Kb | 56 | 4463 | 47 | 3 | 5.0 Mb | 154x |
| Xanthobacter wiegelii CB2 | Xanthobacter sp. CB2 | X | **DSM 14517** | JBAFUW000000000 | PP328857 | 67.7% | 255.1 Kb | 57 | 5058 | 46 | 3 | 5.4 Mb | 263x |
| Xanthobacter wiegelii NCIMB 11399 | Xanthobacter autotrophicus NCIMB 11399 | X | **NCIMB 11399** | JBAFUX000000000 | PP328878 | 67.8% | 325 Kb | 68 | 4979 | 46 | 3 | 5.3 Mb | 183x |
| Xanthobacter wiegelii RH 10 | Xanthobacter autotrophicus RH 10 | X | CIP 105437; **DSM 597**; JCM 7864; TK 0519 | JBAFUY000000000 | PP328872 | 67.7% | 359.9 Kb | 82 | 5266 | 46 | 3 | 5.6 Mb | 135x |
| Roseixanthobacter finlandensis VTT E-85241 | “Xanthobacter polyaromaticivorans” VTT E-85241 | X | **VTT E-85241**; BIO GISA 20; DSM | JBAFWO000000000 | PP328909 | 65.9% | 443.6 Kb | 42 | 5135 | 50 | 3 | 5.6 Mb | 249x |
| Roseixanthobacter glucoisosaccharinivorans VTT E-85242 | “Xanthobacter polyaromaticivorans” VTT E-85242 | X | **VTT E-85242**; BIO GISA 21; DSM | JBAFWN000000000 | PP328910 | 66.0% | 320.7 Kb | 50 | 4976 | 47 | 3 | 5.4 Mb | 192x |
| Roseixanthobacter liquoris VTT E-85238 | “Xanthobacter polyaromaticivorans” VTT E-85238 | X | **VTT E-85238**; BIO GISA 16; DSM | JBAFWP000000000 | PP328906 | 67.2% | 228.3 Kb | 62 | 4479 | 46 | 2 | 5.0 Mb | 201x |
| Roseixanthobacter pseudopolyaromaticivorans E-85237 | “Xanthobacter polyaromaticivorans” E-85237 | X | **VTT E-85237**; BIO GISA 14 | JBAFWS000000000 | PP328905 | 65.8% | 482.6 Kb | 63 | 5029 | 49 | 3 | 5.4 Mb | 191x |
| Roseixanthobacter pseudopolyaromaticivorans E-85239 | “Xanthobacter polyaromaticivorans E-85239 | X | **VTT E-85239**; BIO GISA 18A | JBAFWR000000000 | PP328907 | 65.7% | 545 Kb | 55 | 5082 | 49 | 3 | 5.5 Mb | 169x |
| Roseixanthobacter pseudopolyaromaticivorans E-85240 | “Xanthobacter polyaromaticivorans” E-85240 | X | **VTT E-85240**; BIO GISA 19; DSM | JBAFWQ000000000 | PP328908 | 65.7% | 497.1 Kb | 68 | 5075 | 47 | 3 | 5.5 Mb | 194x |
| Roseixanthobacter psychrophilus W30 | Xanthobacter sp. W30 | X | **DSM 24535;** KCTC | JBAFWT000000000 | PP328867 | 65.3% | 129.2 Kb | 258 | 5815 | 51 | 3 | 6.0 Mb | 189x |
| Ancylobacter aquaticus DSM 101 | Ancylobacter aquaticus DSM 101 | - | ATCC 25396; CCUG 1820; CCUG 30551; DSM 101; JCM 20518; | GCF_004339465.1 | AB681795 | 67.0% | 1.4 Mb | 11 | 4381 | 46 | 3 | 4.8 Mb | 310x |

| | | | | | | | | | | | | | |
|---|---|---|---|---|---|---|---|---|---|---|---|---|---|
| | | | LMG 4052; NBRC 102453; VKM B-1287 | | | | | | | | | | |
| Ancylobacter defluvii SK15 | Ancylobacter defluvii SK15 | - | CCUG 63806; CCUG 63906; DSM 29296; VKM B-2789 | GCF_018390605.1 | KC243678 | 67.2% | 409.9 Kb | 29 | 4857 | 46 | 3 | 5.3 Mb | 149x |
| Ancylobacter dichloromethanicus DM16 | Ancylobacter dichloromethanicus DM16 | - | DSM 21507; VKM B-2484 | GCF_018390645.1 | EU589386 | 67.7% | 243.9 Kb | 64 | 4741 | 46 | 6 | 5.1 Mb | 129x |
| Ancylobacter koreensis Jip08 | Ancylobacter koreensis Jip08 | - | DSM 18406; IAM 15215; JCM 21669; KCTC 12212; NBRC 100963 | GCF_023016525.1 | NR_041013.1 | 68.9% | 616.9 Kb | 19 | 3717 | 48 | 6 | 4.1 Mb | 195x |
| Ancylobacter lacus F30L | Ancylobacter lacus F30L | - | DSM 106329; VKM B-3280 | GCF_018390635.1 | NR_180559.1 | 69.7% | 136.6 Kb | 90 | 4186 | 47 | 3 | 4.6 Mb | 122x |
| Ancylobacter oerskovii NS05 | Ancylobacter oerskovii NS05 | - | CCM 7435; DSM 18746 | GCF_018390555.1 | AM778407 | 68.2% | 416.9 Kb | 46 | 4991 | 50 | 3 | 5.5 Mb | 110x |
| Ancylobacter polymorphus ZM13 | Ancylobacter polymorphus ZM13 | - | - | GCF_022836935.1 | NR_042795.1 | 67.2% | 4.4 Mb | 6 | 4841 | 51 | 6 | 5.2 Mb | 99x |
| Ancylobacter pratisalsi E130 | Ancylobacter pratisalsi E130 | - | DSM 102029; LMG 29367 | GCF_010669125.1 | KX021302 | 65.9% | 4.6 Mb | 2 | 4375 | 49 | 6 | 4.8 Mb | 80x |
| Ancylobacter rudongensis CGMCC 1.1761 | Ancylobacter rudongensis CGMCC 1.1761 | - | DSM 17131; JCM 11671 | GCF_900100155.1 | AY056830 | 67.9% | 4.5 Mb | 18 | 4121 | 46 | 3 | 4.5 Mb | 390x |
| Ancylobacter sonchi Osot | Ancylobacter sonchi Osot | - | DSM 106440; JCM 32039; VKM B-3145 | GCF_018390695.1 | KY492736 | 67.6% | 628.8 Kb | 47 | 5286 | 48 | 4 | 5.8 Mb | 227x |
| Aquabacter caverane Sn-9-2 | Aquabacter caverane Sn-9-2 | - | CCTCC AB 2018270; KCTC 62308 | GCF_003993795.1 | MF958452 | 67.5% | 512.3 Kb | 19 | 3960 | 47 | 5 | 4.5 Mb | 100x |
| Aquabacter spiritensis SPL-1 | Aquabacter spiritensis SPL-1 | - | ATCC 43981; DSM 9035; LMG 8611 | GCF_004346185.1 | FR733686 | 67.6% | 274.7 Kb | 52 | 4600 | 46 | 3 | 5.2 Mb | 290x |
| Azorhizobium caulinodans ORS 571 | Azorhizobium caulinodans ORS 571 | - | ATCC 43989; CCUG 26647; DSM 5975; IFO 14845; JCM 20966; LMG 6465; NBRC 14845 | GCF_000010525.1 | AB680677 | 67.3% | 5.4 Mb | 1 | 4768 | 53 | 9 | 5.4 Mb | - |
| Azorhizobium doebereinerae UFLA1-100 | Azorhizobium doebereinerae UFLA1-100 | - | BR 5401; DSM 18977; LMG 9993; SEMIA 6401 | GCF_000473085.1 | NR_041839.1 | 68.9% | 174.3 Kb | 104 | 5246 | 48 | 8 | 5.8 Mb | - |
| Azorhizobium oxalatiphilum NS12 | Azorhizobium oxalatiphilum NS12 | - | CCM 7897; DSM 18749 | GCF_014635325.1 | NR_108517.1 | 66.6% | 544.7 Kb | 27 | 5642 | 48 | 3 | 6.4 Mb | 97x |
| Pseudoxanthobacter soli CC4 | Pseudoxanthobacter soli CC4 | - | CIP 109513; DSM 19599 | GCF_900148505.1 | EF465533 | 67.8% | 567.2 Kb | 27 | 4306 | 47 | 9 | 5.2 Mb | 374x |
| Xanthobacter aminoxidans 126 | Xanthobacter sp. 126 | - | - | GCF_000526175.1 | - | 67.8% | 5.5 Mb | 1 | 5033 | 50 | 6 | 5.5 Mb | - |
| Xanthobacter aminoxidans 91 | Xanthobacter sp. 91 | - | - | GCF_000702385.1 | - | 68.1% | 5.3 Mb | 1 | 4885 | 48 | 6 | 5.3 Mb | - |
| Xanthobacter dioxanivorans YN2 | Xanthobacter dioxanivorans YN2 | - | CGMCC 1.19031; JCM 34666 | GCF_016807805.1 | MZ851448 | 67.9% | 6.3 Mb | 5 | 6012 | 49 | 6 | 6.7 Mb | 221x |
| Xanthobacter flavus GJ10 | Xanthobacter autotrophicus GJ10 | - | ATCC 43050; CCM 4398; **DSM 3874**; NCAIM B.01957; NRRL B-14837 | GCF_020683165.1 | - | 67.6% | 5.2 Mb | 1 | 4862 | 42 | 3 | 5.2 Mb | 2.5x |
| Xanthobacter flavus YC-JY1 | Xanthobacter sp. YC-JY1 | - | - | GCF_021730305.1 | - | 68.0% | 5.4 Mb | 1 | 4943 | 48 | 6 | 5.4 Mb | 341x |
| Xanthobacter flavus SG618 | Xanthobacter sp. SG618 | - | - | GCF_012932745.1 | - | 68.2% | 537 Kb | 15 | 4747 | 46 | 3 | 5.2 Mb | 288x |

| | | | | | | | | | | | | | |
|---|---|---|---|---|---|---|---|---|---|---|---|---|---|
| “Xanthobacter gandavensis” NM-25 | “Xanthobacter gandavensis” NM-25 | - | - | GCF_022138605.1 | - | 67.8% | 625.6 Kb | 18 | 4248 | 48 | 6 | 4.8 Mb | 65x |
| “Xanthobacter gandavensis” NFM-26 | “Xanthobacter gandaensis” NFM-26 | - | - | GCF_022138625.1 | MT623481.1 | 67.8% | 611.7 Kb | 20 | 4249 | 46 | 3 | 4.8 Mb | 61x |
| Xanthobacter oligotrophicus 29k | Xanthobacter oligotrophicus 29k | - | KCTC 72777; VKM B-3453 | GCF_008364685.1 | NR_181626.1 | 67.9% | 342 Kb | 35 | 4688 | 47 | 4 | 5.3 Mb | 100.8x |
| Xanthobacter oligotrophicus BF-7S | Xanthobacter oligotrophicus BF-7S | - | - | GCF_022213975.1 | | 67.9% | 458.1 Kb | 31 | 4670 | 49 | 4 | 5.3 Mb | 286.8x |
| Xanthobacter sp. AM3 | Xanthobacter autotrophicus AM3 | - | - | GCF_030015065.1 | - | 68.0% | 31.1 Kb | 594 | 4672 | 52 | 6 | 5 Mb | 30x |
| Xanthobacter sp. AM5 | Xanthobacter autotrophicus AM5 | - | - | GCF_029991245.1 | - | 67.9% | 27.6 Kb | 495 | 4494 | 50 | 5 | 5 Mb | 30x |
| Xanthobacter sp. SG518 | Xanthobacter sp. SG518 | - | - | Gp0393885 (JGI) | - | 68.0% | - | - | - | - | - | - | - |
| Xanthobacter sp. SG563 | Xanthobacter sp. SG563 | - | - | Gp0393886 (JGI) | - | 68.0% | - | - | - | - | - | - | - |
| Xanthobacter variabilis NM-44 | Xanthobacter variabilis NM-44 | - | - | GCF_021401105.1 | - | 66.8% | 306.1 Kb | 38 | 4255 | 48 | 4 | 5 Mb | 74x |
| Xanthobacter variabilis NM-59 | Xanthobacter variabilis NM-59 | - | - | GCF_021401125.1 | - | 66.8% | 304.8 Kb | 36 | 4250 | 48 | 4 | 5 Mb | 66x |
| Xanthobacter variabilis R2A-8 | Xanthobacter variabilis R2A-8 | - | - | GCF_021401165.1 | - | 66.8% | 196.1 Kb | 49 | 4252 | 48 | 4 | 5 Mb | 77x |
| Xanthobacter variabilis NFM-97 | Xanthobacter variabilis NFM-97 | - | - | GCF_021401185.1 | - | 66.8% | 277 Kb | 42 | 4407 | 48 | 4 | 5.1 Mb | 69x |
| Xanthobacter variabilis NFM-89 | Xanthobacter variabilis NFM-89 | - | - | GCF_021401225.1 | - | 66.7% | 701.8 Kb | 32 | 4433 | 50 | 6 | 5.1 Mb | 53x |
| Xanthobacter variabilis NFM-59 | Xanthobacter variabilis NFM-59 | - | - | GCF_021401255.1 | - | 66.8% | 305.9 Kb | 37 | 4250 | 48 | 4 | 5 Mb | 70x |
| Xanthobacter variabilis NFH-94 | Xanthobacter variabilis NFH-94 | - | - | GCF_021401295.1 | - | 66.8% | 283.1 Kb | 41 | 4257 | 48 | 4 | 5 Mb | 70x |
| Xanthobacter variabilis NFH-6 | Xanthobacter variabilis NFH-6 | - | - | GCF_021401325.1 | - | 66.7% | 282.8 Kb | 34 | 4434 | 48 | 4 | 5.1 Mb | 65x |
| Xanthobacter variabilis NFH-44 | Xanthobacter variabilis NFH-44 | - | - | GCF_021401365.1 | - | 66.8% | 304.7 Kb | 39 | 4256 | 48 | 4 | 5 Mb | 79x |
| Xanthobacter versatilis Py2 | Xanthobacter autotrophicus Py2 | - | **ATCC BAA-1158**; DSM | GCF_000017645.1 | NR_074255.1 | 67.3% | 5.3 Mb | 2 | 5076 | 48 | 6 | 5.6 Mb | - |
| Xanthobacter wiegelii WS4821 | Xanthobacter flavus WS4821 | - | - | GCF_017875275.1 | - | 67.8% | 5.4 Mb | 1 | 4906 | 48 | 6 | 5.4 Mb | 173x |
| “Xanthobacter polyaromaticivorans” 127W | “Xanthobacter polyaromaticivorans” 127W | - | - | - | AB106864 | - | - | - | - | - | - | - | - |
| “Xanthobacter xylophilus” Z-0055 | “Xanthobacter xylophilus” Z-0055 | - | VKM B-2535 | - | FJ882073 | - | - | - | - | - | - | - | - |

**Supplementary Figure 1. The evolutionary history of all Xanthobacter and Roseixanthobacter gen. nov. strains inferred using the neighbor-joining method with 120 core, conserved, single-copy bacterial genes indicates robust evolutionary groupings.** The phylogenomic tree was generated using MEGA-X with the neighbor-joining method with the partial deletion (95% cutoff) option and was bootstrapped 1000 times (with bootstrap percentages shown next to the branch). Type strains are underlined and genomes that we sequenced in this study are bolded. Related genera Azorhizobium, Aquabacter, and Ancylobacter are included. *Pseudoxanthobacter soli* CC4$^T$ is used as an outgroup. The scale bar represents 0.05 amino acid substitutions per site as computed using the Poisson correction method.

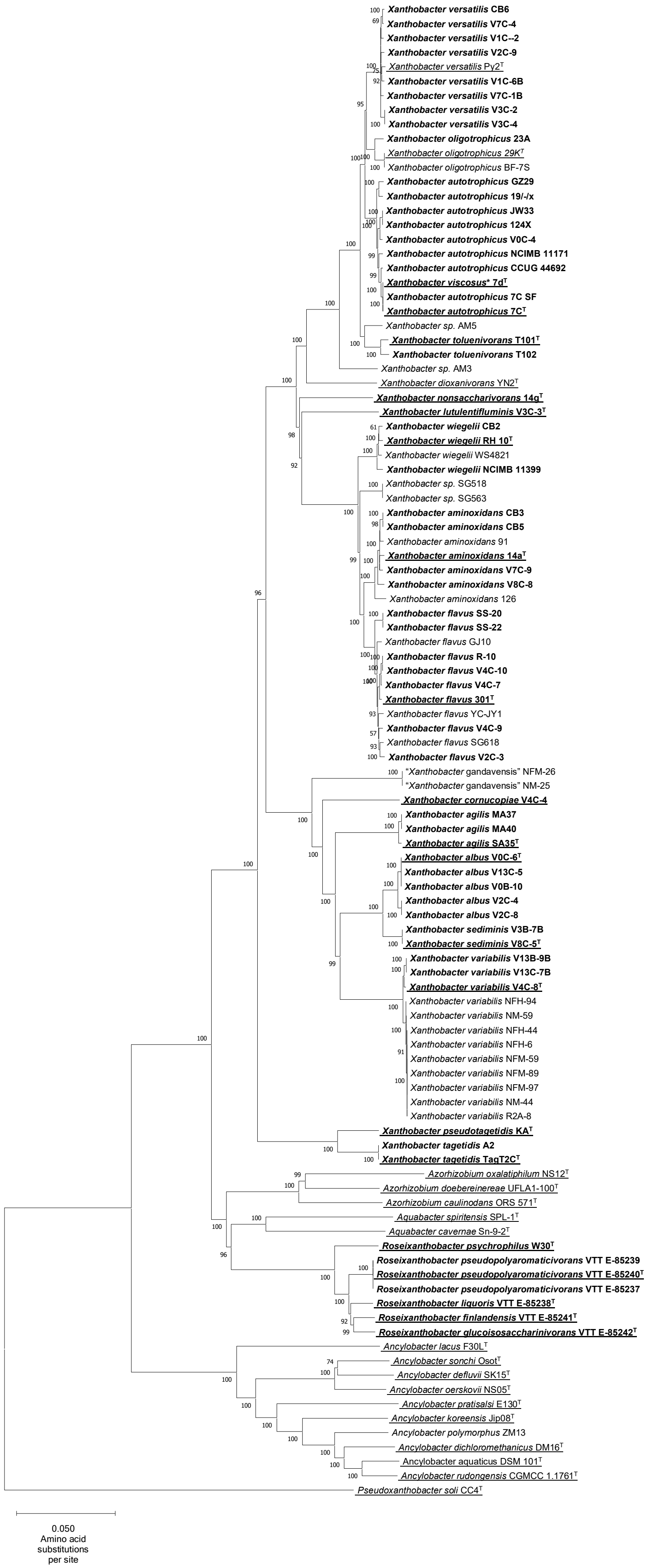

Xanthobacter versatilis CB6
Xanthobacter versatilis V7C-4
Xanthobacter versatilis V1C--2
Xanthobacter versatilis V2C-9
Xanthobacter versatilis Py2$^{T}$
Xanthobacter versatilis V1C-6B
Xanthobacter versatilis V7C-1B
Xanthobacter versatilis V3C-2
Xanthobacter versatilis V3C-4
Xanthobacter oligotrophicus 23A
Xanthobacter oligotrophicus 29K$^{T}$
Xanthobacter oligotrophicus BF-7S
Xanthobacter autotrophicus GZ29
Xanthobacter autotrophicus 19/-/x
Xanthobacter autotrophicus JW33
Xanthobacter autotrophicus 124X
Xanthobacter autotrophicus V0C-4
Xanthobacter autotrophicus NCIMB 11171
Xanthobacter autotrophicus CCUG 44692
Xanthobacter viscosus* 7d$^{T}$
Xanthobacter autotrophicus 7C SF
Xanthobacter autotrophicus 7C$^{T}$
Xanthobacter sp. AM5
Xanthobacter toluenivorans T101$^{T}$
Xanthobacter toluenivorans T102
Xanthobacter sp. AM3
Xanthobacter dioxanivorans YN2$^{T}$
Xanthobacter nonsaccharivorans 14g$^{T}$
Xanthobacter lutulentifluminis V3C-3$^{T}$
Xanthobacter wiegelii CB2
Xanthobacter wiegelii RH 10$^{T}$
Xanthobacter wiegelii WS4821
Xanthobacter wiegelii NCIMB 11399
Xanthobacter sp. SG518
Xanthobacter sp. SG563
Xanthobacter aminoxidans CB3
Xanthobacter aminoxidans CB5
Xanthobacter aminoxidans 91
Xanthobacter aminoxidans 14a$^{T}$
Xanthobacter aminoxidans V7C-9
Xanthobacter aminoxidans V8C-8
Xanthobacter aminoxidans 126
Xanthobacter flavus SS-20
Xanthobacter flavus SS-22
Xanthobacter flavus GJ10
Xanthobacter flavus R-10
Xanthobacter flavus V4C-10
Xanthobacter flavus V4C-7
Xanthobacter flavus 301$^{T}$
Xanthobacter flavus YC-JY1
Xanthobacter flavus V4C-9
Xanthobacter flavus SG618
Xanthobacter flavus V2C-3
"Xanthobacter gandavensis" NFM-26
"Xanthobacter gandavensis" NM-25
Xanthobacter cornucopiae V4C-4
Xanthobacter agilis MA37
Xanthobacter agilis MA40
Xanthobacter agilis SA35$^{T}$
Xanthobacter albus V0C-6$^{T}$
Xanthobacter albus V13C-5
Xanthobacter albus V0B-10
Xanthobacter albus V2C-4
Xanthobacter albus V2C-8
Xanthobacter sediminis V3B-7B
Xanthobacter sediminis V8C-5$^{T}$
Xanthobacter variabilis V13B-9B
Xanthobacter variabilis V13C-7B
Xanthobacter variabilis V4C-8$^{T}$
Xanthobacter variabilis NFH-94
Xanthobacter variabilis NM-59
Xanthobacter variabilis NFH-44
Xanthobacter variabilis NFH-6
Xanthobacter variabilis NFM-59
Xanthobacter variabilis NFM-89
Xanthobacter variabilis NFM-97
Xanthobacter variabilis NM-44
Xanthobacter variabilis R2A-8
Xanthobacter pseudotagetidis KA$^{T}$
Xanthobacter tagetidis A2
Xanthobacter tagetidis TagT2C$^{T}$
Azorhizobium oxalatiphilum NS12$^{T}$
Azorhizobium doebereinereae UFLA1-100$^{T}$
Azorhizobium caulinodans ORS 571$^{T}$
Aquabacter spiritensis SPL-1$^{T}$
Aquabacter cavernae Sn-9-2$^{T}$
Roseixanthobacter psychrophilus W30$^{T}$
Roseixanthobacter pseudopolyaromaticivorans VTT E-85239
Roseixanthobacter pseudopolyaromaticivorans VTT E-85240$^{T}$
Roseixanthobacter pseudopolyaromaticivorans VTT E-85237
Roseixanthobacter liquoris VTT E-85238$^{T}$
Roseixanthobacter finlandensis VTT E-85241$^{T}$
Roseixanthobacter glucoisosaccharinivorans VTT E-85242$^{T}$
Ancylobacter lacus F30L$^{T}$
Ancylobacter sonchi Osot$^{T}$
Ancylobacter defluvii SK15$^{T}$
Ancylobacter oerskovii NS05$^{T}$
Ancylobacter pratisalsi E130$^{T}$
Ancylobacter koreensis Jip08$^{T}$
Ancylobacter polymorphus ZM13
Ancylobacter dichloromethanicus DM16$^{T}$
Ancylobacter aquaticus DSM 101$^{T}$
Ancylobacter rudongensis CGMCC 1.1761$^{T}$
Pseudoxanthobacter soli CC4$^{T}$
0.050
Amino acid
substitutions
per site

**Supplementary Figure 2. The evolutionary history of all Xanthobacter and Roseixanthobacter gen. nov. strains inferred using the maximum parsimony method with 120 core, conserved, single-copy bacterial genes indicates robust evolutionary groupings.** The phylogenomic tree was generated using MEGA-X with the maximum parsimony method with the partial deletion (95% cutoff) option and was bootstrapped 1000 times (with bootstrap percentages shown next to the branch). Type strains are underlined and genomes that we sequenced in this study are bolded. Related genera Azorhizobium, Aquabacter, and Ancylobacter are included. *Pseudoxanthobacter soli* CC4$^{T}$ is used as an outgroup. Branch lengths could not be calculated due to computational constraints (large number of sequences each with large sequence length).

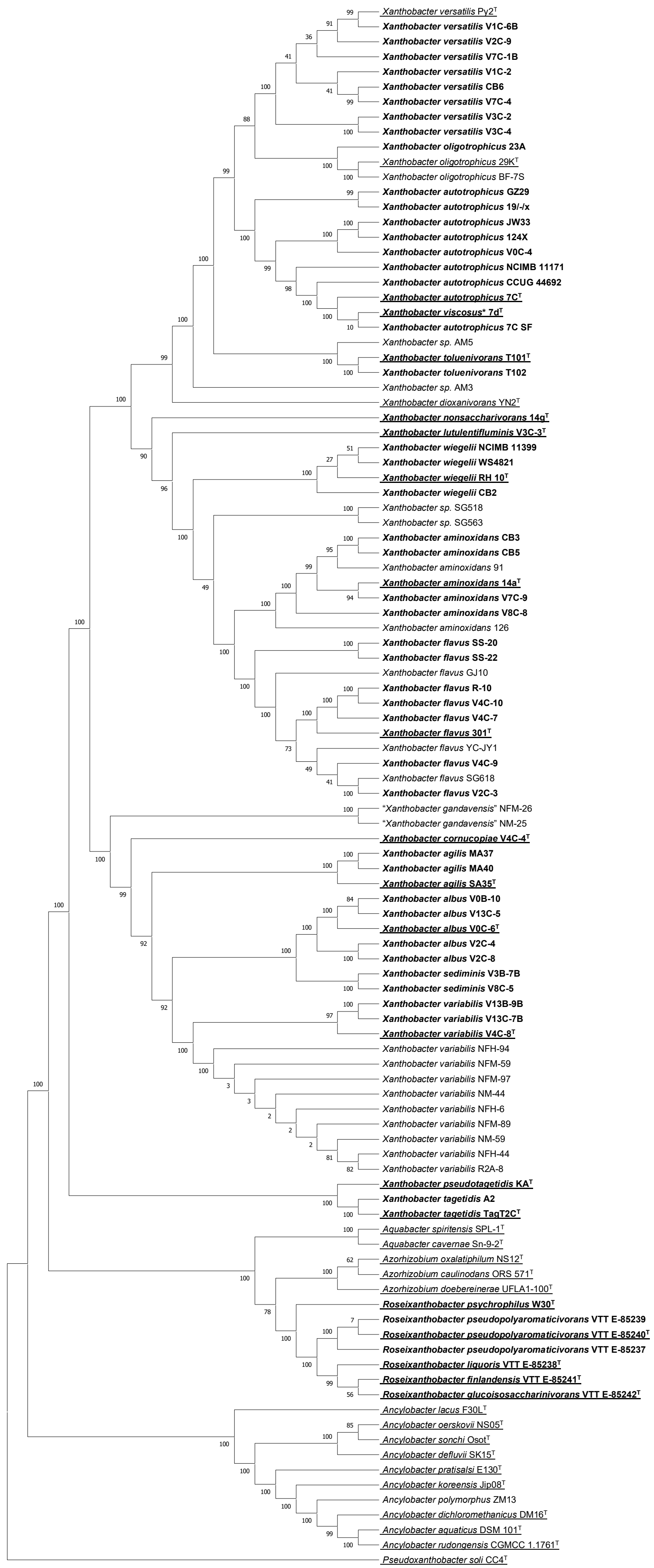
Xanthobacter versatilis Py2$^{T}$
Xanthobacter versatilis V1C-6B
Xanthobacter versatilis V2C-9
Xanthobacter versatilis V7C-1B
Xanthobacter versatilis V1C-2
Xanthobacter versatilis CB6
Xanthobacter versatilis V7C-4
Xanthobacter versatilis V3C-2
Xanthobacter versatilis V3C-4
Xanthobacter oligotrophicus 23A
Xanthobacter oligotrophicus 29K$^{T}$
Xanthobacter oligotrophicus BF-7S
Xanthobacter autotrophicus GZ29
Xanthobacter autotrophicus 19/-/x
Xanthobacter autotrophicus JW33
Xanthobacter autotrophicus 124X
Xanthobacter autotrophicus V0C-4
Xanthobacter autotrophicus NCIMB 11171
Xanthobacter autotrophicus CCUG 44692
Xanthobacter autotrophicus 7C$^{T}$
Xanthobacter viscosus* 7d$^{T}$
Xanthobacter autotrophicus 7C SF
Xanthobacter sp. AM5
Xanthobacter toluenivorans T101$^{T}$
Xanthobacter toluenivorans T102
Xanthobacter sp. AM3
Xanthobacter dioxanivorans YN2$^{T}$
Xanthobacter nonsaccharivorans 14g$^{T}$
Xanthobacter lutulentifluminis V3C-3$^{T}$
Xanthobacter wiegelii NCIMB 11399
Xanthobacter wiegelii WS4821
Xanthobacter wiegelii RH 10$^{T}$
Xanthobacter wiegelii CB2
Xanthobacter sp. SG518
Xanthobacter sp. SG563
Xanthobacter aminoxidans CB3
Xanthobacter aminoxidans CB5
Xanthobacter aminoxidans 91
Xanthobacter aminoxidans 14a$^{T}$
Xanthobacter aminoxidans V7C-9
Xanthobacter aminoxidans V8C-8
Xanthobacter aminoxidans 126
Xanthobacter flavus SS-20
Xanthobacter flavus SS-22
Xanthobacter flavus GJ10
Xanthobacter flavus R-10
Xanthobacter flavus V4C-10
Xanthobacter flavus V4C-7
Xanthobacter flavus 301$^{T}$
Xanthobacter flavus YC-JY1
Xanthobacter flavus V4C-9
Xanthobacter flavus SG618
Xanthobacter flavus V2C-3
"Xanthobacter gandavensis" NFM-26
"Xanthobacter gandavensis" NM-25
Xanthobacter cornucopiae V4C-4$^{T}$
Xanthobacter agilis MA37
Xanthobacter agilis MA40
Xanthobacter agilis SA35$^{T}$
Xanthobacter albus V0B-10
Xanthobacter albus V13C-5
Xanthobacter albus V0C-6$^{T}$
Xanthobacter albus V2C-4
Xanthobacter albus V2C-8
Xanthobacter sediminis V3B-7B
Xanthobacter sediminis V8C-5
Xanthobacter variabilis V13B-9B
Xanthobacter variabilis V13C-7B
Xanthobacter variabilis V4C-8$^{T}$
Xanthobacter variabilis NFH-94
Xanthobacter variabilis NFM-59
Xanthobacter variabilis NFM-97
Xanthobacter variabilis NM-44
Xanthobacter variabilis NFH-6
Xanthobacter variabilis NFM-89
Xanthobacter variabilis NM-59
Xanthobacter variabilis NFH-44
Xanthobacter variabilis R2A-8
Xanthobacter pseudotagetidis KA$^{T}$
Xanthobacter tagetidis A2
Xanthobacter tagetidis TagT2C$^{T}$
Aquabacter spiritensis SPL-1$^{T}$
Aquabacter cavernae Sn-9-2$^{T}$
Azorhizobium oxalatiphilum NS12$^{T}$
Azorhizobium caulinodans ORS 571$^{T}$
Azorhizobium doebereinerae UFLA1-100$^{T}$
Roseixanthobacter psychrophilus W30$^{T}$
Roseixanthobacter pseudopolyaromaticivorans VTT E-85239
Roseixanthobacter pseudopolyaromaticivorans VTT E-85240$^{T}$
Roseixanthobacter pseudopolyaromaticivorans VTT E-85237
Roseixanthobacter liquoris VTT E-85238$^{T}$
Roseixanthobacter finlandensis VTT E-85241$^{T}$
Roseixanthobacter glucoisosaccharinivorans VTT E-85242$^{T}$
Ancylobacter lacus F30L$^{T}$
Ancylobacter oerskovii NS05$^{T}$
Ancylobacter sonchi Osot$^{T}$
Ancylobacter defluvii SK15$^{T}$
Ancylobacter pratisalsi E130$^{T}$
Ancylobacter koreensis Jip08$^{T}$
Ancylobacter polymorphus ZM13
Ancylobacter dichloromethanicus DM16$^{T}$
Ancylobacter aquaticus DSM 101$^{T}$
Ancylobacter rudongensis CGMCC 1.1761$^{T}$
Pseudoxanthobacter soli CC4$^{T}$

**Supplementary Figure 3. The evolutionary history of Xanthobacter and Roseixanthobacter gen. nov. species type strains inferred using the maximum likelihood method with 16S rRNA genes cannot resolve genus level boundaries between Xanthobacter, Roseixanthobacter, Azorhizobium, and Aquabacter.** The phylogenetic tree was generated using MEGA-X with the K2+G(5)+I maximum likelihood model with the partial deletion (95% cutoff) option and was bootstrapped 1000 times (with bootstrap percentages shown next to the branch). Novel species described in this paper are bolded while genome sequence accessions that we sequenced in this study are underlined. Related genera Azorhizobium, Aquabacter, and Ancylobacter are included. *Pseudoxanthobacter soli* CC4$^T$ is used as an outgroup. The scale bar represents 0.02 nucleotide substitutions per site.

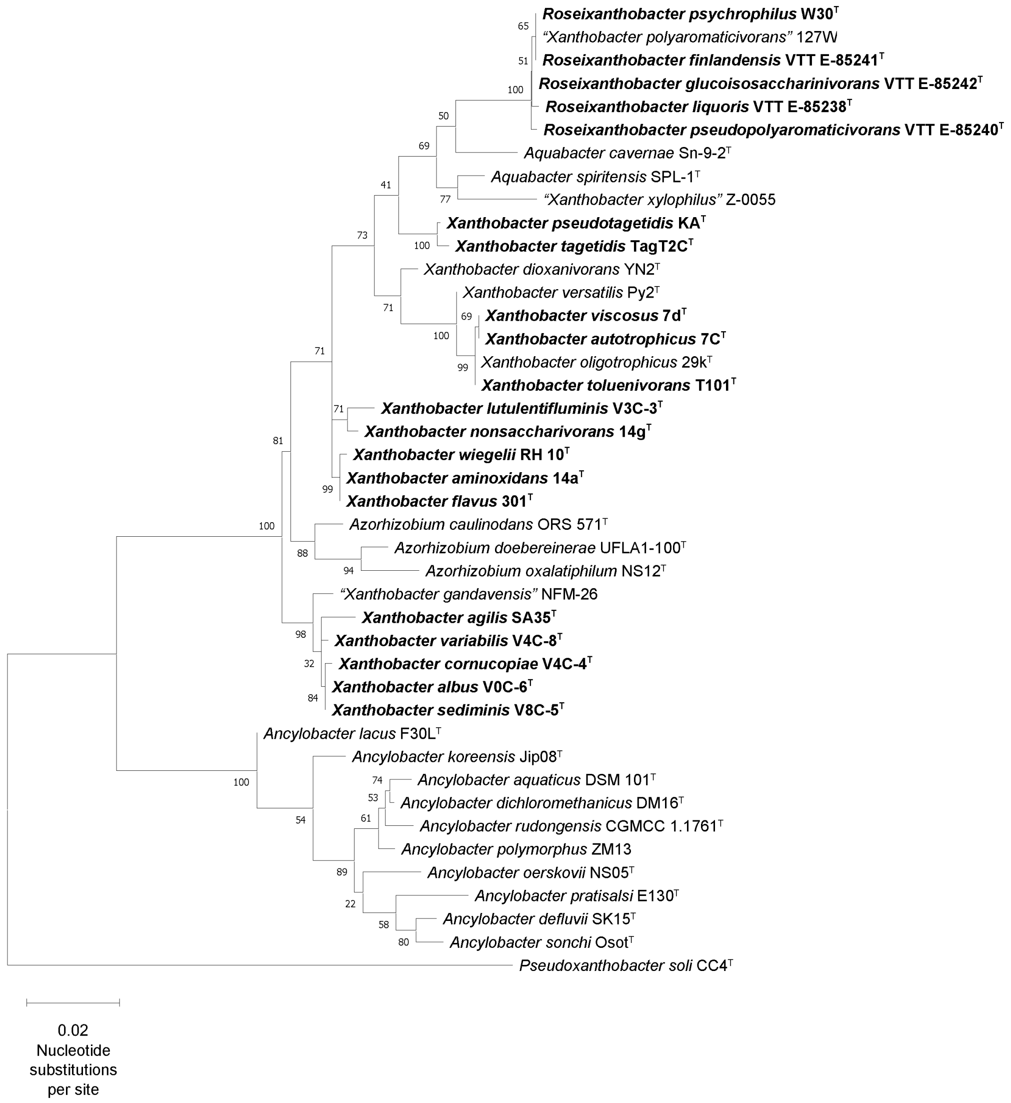

**Supplementary Figure 4. The evolutionary history of Xanthobacter and Roseixanthobacter gen. nov. species type strains inferred using the neighbor-joining method with 16S rRNA genes cannot resolve genus level boundaries between Xanthobacter, Roseixanthobacter, Azorhizobium, and Aquabacter.** The phylogenetic tree was generated using MEGA-X with the neighbor-joining method with the partial deletion (95% cutoff) option and was bootstrapped 1000 times (with bootstrap percentages shown next to the branch). Novel species described in this paper are bolded while genome sequence accessions that we sequenced in this study are underlined. Related genera Azorhizobium, Aquabacter, and Ancylobacter are included. *Pseudoxanthobacter soli* CC4$^{T}$ is used as an outgroup. The scale bar represents 0.01 nucleotide substitutions per site as computed by the maximum composite likelihood method.

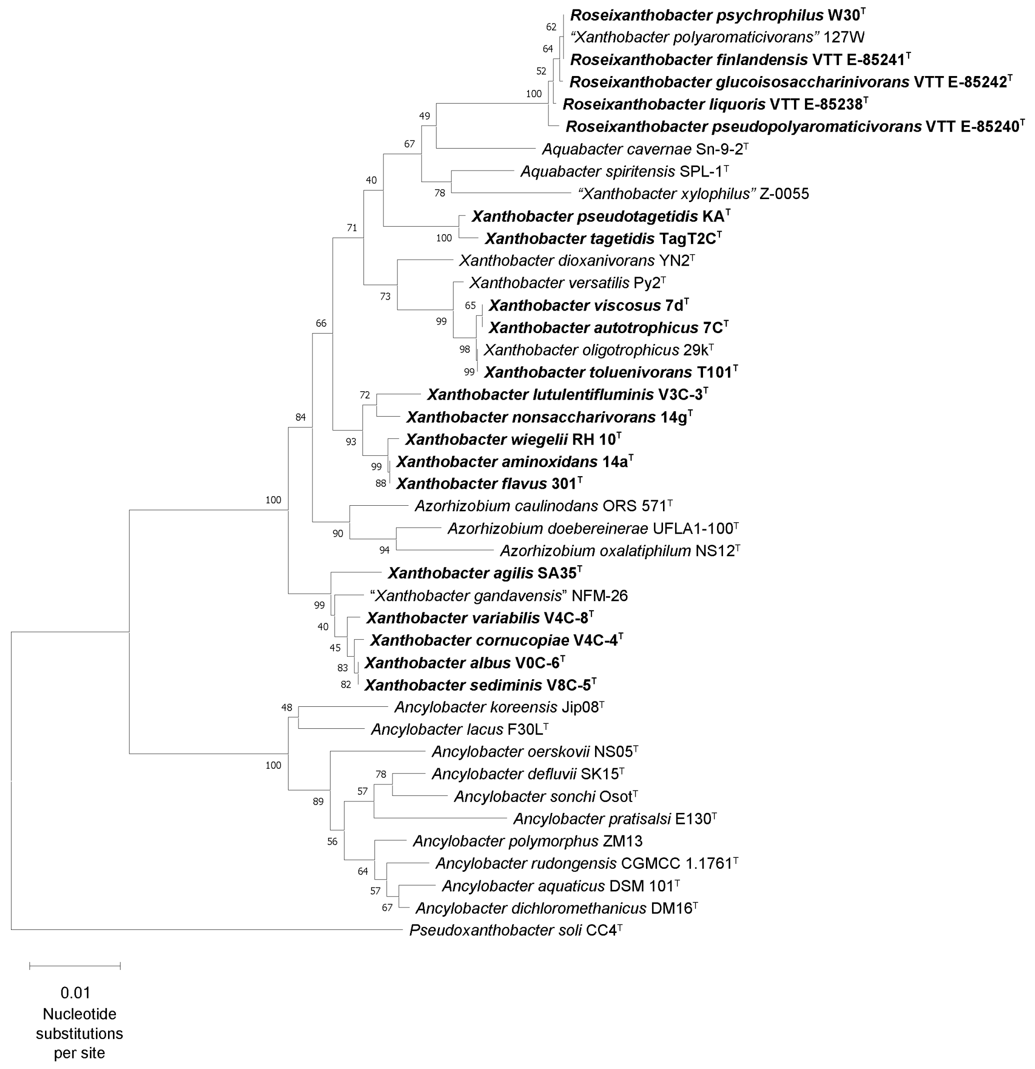

**Supplementary Figure 5. The evolutionary history of Xanthobacter and Roseixanthobacter gen. nov. species type strains inferred using the maximum parsimony method with 16S rRNA genes cannot resolve genus level boundaries between Xanthobacter, Roseixanthobacter, Azorhizobium, and Aquabacter.** The phylogenetic tree was generated using MEGA-X with the maximum parsimony method with the partial deletion (95% cutoff) option and was bootstrapped 1000 times (with bootstrap percentages shown next to the branch). Novel species described in this paper are bolded while genome sequence accessions that we sequenced in this study are underlined.  Related genera Azorhizobium, Aquabacter, and Ancylobacter are included. *Pseudoxanthobacter soli* CC4$^T$ is used as an outgroup. The scale bar is in units of the number of changes over the whole sequence and branch lengths were calculated using the average pathway method.

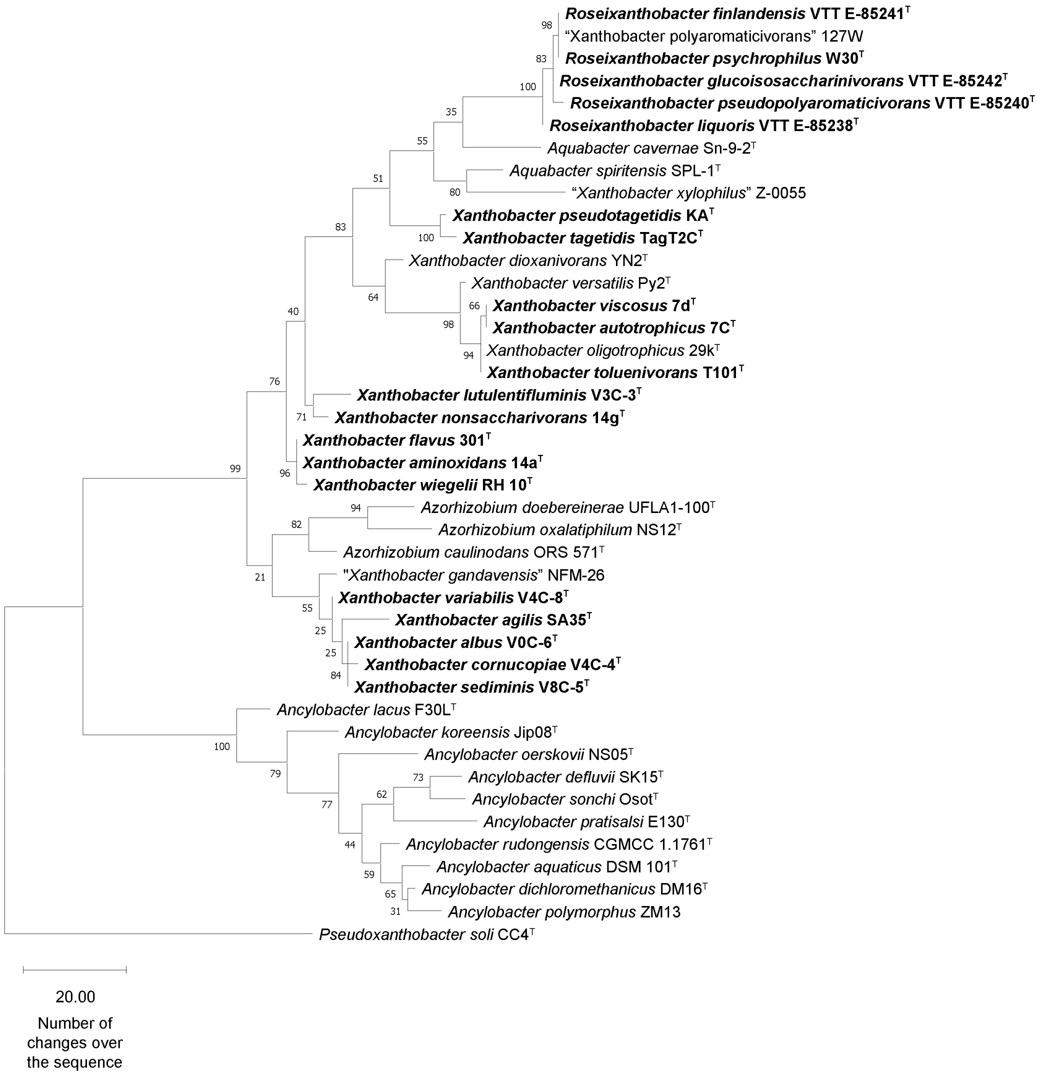

**Supplementary Table 3 – Xanthobacter and Roseixanthobacter gen. nov. strains can be separated into species clusters using average nucleotide identity values (ANI) calculated using FastANI with a 95% species level threshold.** Values above the 95% ANI score species threshold are colored in blue while those below the species threshold are colored in red.

| | Xanthobacter versatilis CB6 | Xanthobacter versatilis V7C-4 | Xanthobacter versatilis V1C-2 | Xanthobacter versatilis V2C-9 | Xanthobacter versatilis Py2 | Xanthobacter versatilis V1C-6B | Xanthobacter versatilis V7C-1B | Xanthobacter versatilis V3C-2 | Xanthobacter versatilis V3C-4 | Xanthobacter oligotrophicus 23A | Xanthobacter oligotrophicus 29k | Xanthobacter oligotrophicus BF-7S | Xanthobacter autotrophicus GZ29 | Xanthobacter autotrophicus 19/-/x | Xanthobacter autotrophicus JW33 | Xanthobacter autotrophicus 124X | Xanthobacter autotrophicus V0C-4 | Xanthobacter autotrophicus NCIMB 11171 | Xanthobacter autotrophicus CCUG 44692 | Xanthobacter viscosus 7d | Xanthobacter autotrophicus 7C SF | Xanthobacter autotrophicus 7C | Xanthobacter sp. AM5 | Xanthobacter toluenivorans T101 | Xanthobacter toluenivorans T102 | Xanthobacter sp. AM3 | Xanthobacter dioxanivorans YN2 | Xanthobacter nonsaccharivorans 14g | Xanthobacter lutulentifluminis V3C-3 | Xanthobacter wiegelii CB2 | Xanthobacter wiegelii RH 10 | Xanthobacter wiegelii WS4821 | Xanthobacter wiegelii NCIMB 11399 | Xanthobacter sp. SG518 | Xanthobacter sp. SG563 | Xanthobacter aminoxidans CB3 | Xanthobacter aminoxidans CB5 | Xanthobacter aminoxidans 91 | Xanthobacter aminoxidans 14a | Xanthobacter aminoxidans V7C-9 | Xanthobacter aminoxidans V8C-8 | Xanthobacter aminoxidans 126 | Xanthobacter flavus SS-20 | Xanthobacter flavus SS-22 | Xanthobacter flavus GJ10 | Xanthobacter flavus R-10 | Xanthobacter flavus V4C-10 | Xanthobacter flavus V4C-7 | Xanthobacter flavus 301 | Xanthobacter flavus YC-JY1 | Xanthobacter flavus V4C-9 | Xanthobacter flavus SG618 | Xanthobacter flavus V2C-3 |
|---|---|---|---|---|---|---|---|---|---|---|---|---|---|---|---|---|---|---|---|---|---|---|---|---|---|---|---|---|---|---|---|---|---|---|---|---|---|---|---|---|---|---|---|---|---|---|---|---|---|---|---|---|---|
| Xanthobacter versatilis CB6 | 100 | 98.7791 | 98.3348 | 98.6853 | 98.504 | 98.4613 | 98.5452 | 98.6096 | 98.6155 | 93.3383 | 93.777 | 93.7789 | 92.2533 | 92.3843 | 92.3769 | 92.2938 | 92.3399 | 92.2438 | 92.2846 | 92.3955 | 92.4018 | 92.3762 | 91.9983 | 91.3291 | 91.4274 | 90.3217 | 85.6049 | 85.2661 | 85.9873 | 84.7199 | 85.2375 | 85.3333 | 84.8968 | 84.732 | 84.7585 | 85.0669 | 85.1043 | 85.0811 | 85.1097 | 85.607 | 85.2889 | 84.9969 | 85.027 | 85.0274 | 85.3879 | 84.9221 | 85.4379 | 84.975 | 85.3198 | 85.1487 | 85.0049 | 84.9962 | 85.6056 |
| Xanthobacter versatilis V7C-4 | 98.7609 | 100 | 98.4776 | 98.7078 | 98.394 | 98.6065 | 98.4715 | 98.3879 | 98.3798 | 93.4287 | 93.5824 | 93.5848 | 92.2294 | 92.3788 | 92.286 | 92.3463 | 92.4522 | 92.3542 | 92.306 | 92.3456 | 92.3533 | 92.3783 | 91.9209 | 91.3018 | 91.3083 | 90.1815 | 85.5734 | 85.215 | 85.9388 | 84.7391 | 85.0919 | 85.0184 | 84.7195 | 84.6797 | 84.6817 | 85.0344 | 84.9897 | 84.9682 | 85.1023 | 85.2531 | 85.0551 | 84.9484 | 85.1442 | 85.0467 | 85.2356 | 85.0207 | 85.289 | 84.9472 | 85.2259 | 85.0318 | 84.9794 | 84.9667 | 85.2557 |
| Xanthobacter versatilis V1C-2 | 98.3954 | 98.5477 | 100 | 98.3637 | 98.2724 | 98.4889 | 98.4226 | 98.2195 | 98.3551 | 93.609 | 93.4382 | 93.445 | 92.2749 | 92.4398 | 92.3267 | 92.3784 | 92.4156 | 92.4584 | 92.4085 | 92.3632 | 92.3712 | 92.3423 | 92.0599 | 91.2959 | 91.3961 | 90.0376 | 85.4694 | 85.1981 | 85.8273 | 84.7266 | 84.7037 | 84.7569 | 84.7816 | 84.7089 | 84.7535 | 85.0778 | 85.0643 | 85.0273 | 85.6705 | 85.0315 | 84.9189 | 84.9309 | 85.116 | 85.0171 | 85.1276 | 85.0136 | 84.987 | 84.9112 | 84.9581 | 85.0033 | 84.9981 | 85.0436 | 84.9588 |
| Xanthobacter versatilis V2C-9 | 98.6304 | 98.6017 | 98.3872 | 100 | 98.284 | 98.4235 | 98.4091 | 98.2719 | 98.2368 | 93.3618 | 93.7342 | 93.7765 | 92.1821 | 92.3748 | 92.2455 | 92.1942 | 92.3557 | 92.1729 | 92.2044 | 92.3466 | 92.3731 | 92.4147 | 91.7183 | 91.2962 | 91.1541 | 90.2653 | 85.5599 | 85.2302 | 85.9621 | 84.7767 | 85.105 | 85.3375 | 84.8282 | 84.9285 | 84.9223 | 85.1051 | 85.1198 | 85.1423 | 85.2326 | 85.6282 | 85.1376 | 85.0645 | 85.1307 | 85.1241 | 85.2789 | 85.0238 | 85.5015 | 84.9708 | 85.3018 | 85.1685 | 85.1504 | 85.0838 | 85.5954 |
| Xanthobacter versatilis Py2 | 98.3834 | 98.3095 | 98.2263 | 98.2452 | 100 | 98.195 | 98.1858 | 98.1299 | 98.0919 | 93.2305 | 93.527 | 93.5469 | 91.951 | 92.225 | 92.0571 | 92.2047 | 92.1507 | 92.1632 | 92.0745 | 92.2876 | 92.2553 | 92.2639 | 91.6397 | 91.1784 | 91.3114 | 90.0981 | 85.7421 | 85.1833 | 86.0082 | 85.3175 | 85.39 | 85.1426 | 84.8399 | 84.7268 | 84.6799 | 84.9205 | 84.8914 | 85.0629 | 84.9976 | 85.3692 | 84.9923 | 84.9104 | 85.0662 | 84.9966 | 85.0948 | 85.2069 | 85.2513 | 85.0074 | 85.3447 | 84.9736 | 85.0613 | 85.0076 | 85.3353 |
| Xanthobacter versatilis V1C-6B | 98.4151 | 98.6214 | 98.4953 | 98.374 | 98.245 | 100 | 98.4061 | 98.2915 | 98.315 | 93.4533 | 93.5852 | 93.5938 | 92.2177 | 92.3448 | 92.3167 | 92.2923 | 92.3594 | 92.251 | 92.4178 | 92.4113 | 92.3536 | 92.3637 | 91.841 | 91.2758 | 91.3569 | 90.1989 | 85.4599 | 85.1864 | 85.8566 | 84.8147 | 84.9829 | 85.0946 | 84.8026 | 84.7506 | 84.7725 | 85.0085 | 85.0092 | 85.0653 | 85.132 | 85.2774 | 85.0399 | 84.8477 | 84.9342 | 84.9496 | 85.5674 | 85.0384 | 85.1358 | 84.9322 | 85.2401 | 85.0339 | 84.942 | 84.9649 | 85.2673 |
| Xanthobacter versatilis V7C-1B | 98.4641 | 98.4084 | 98.3631 | 98.4067 | 98.2861 | 98.4085 | 100 | 98.2248 | 98.2352 | 93.2724 | 93.7066 | 93.7415 | 92.2439 | 92.354 | 92.3704 | 92.214 | 92.3118 | 92.2936 | 92.1979 | 92.3546 | 92.3347 | 92.3069 | 91.6899 | 91.2714 | 91.1855 | 90.359 | 85.4725 | 85.2846 | 86.2211 | 84.7894 | 85.0907 | 85.4816 | 84.773 | 84.8058 | 84.8504 | 85.091 | 85.0612 | 85.0687 | 85.0726 | 85.4877 | 84.9462 | 84.9105 | 85.159 | 85.057 | 85.1662 | 85.0056 | 85.5022 | 84.9225 | 85.4207 | 85.1882 | 85.1289 | 84.9805 | 85.5428 |
| Xanthobacter versatilis V3C-2 | 98.6202 | 98.4157 | 98.2428 | 98.3274 | 98.1298 | 98.3068 | 98.2987 | 100 | 99.9677 | 93.2574 | 93.6129 | 93.5949 | 92.106 | 92.331 | 92.1414 | 92.1232 | 92.3563 | 92.1714 | 92.1134 | 92.2406 | 92.28 | 92.2264 | 91.7249 | 91.1712 | 91.1943 | 90.0776 | 85.6514 | 85.3805 | 85.8721 | 84.8197 | 85.0989 | 85.1947 | 84.7827 | 84.7566 | 84.739 | 85.096 | 85.1463 | 85.2051 | 85.1348 | 85.5542 | 85.0372 | 85.0602 | 84.9939 | 85.0483 | 85.2722 | 84.9774 | 85.4011 | 85.0049 | 85.4722 | 85.1537 | 84.9891 | 84.9907 | 85.4593 |
| Xanthobacter versatilis V3C-4 | 98.5352 | 98.38 | 98.3046 | 98.2691 | 98.1801 | 98.2764 | 98.2865 | 99.9211 | 100 | 93.2851 | 93.5287 | 93.5884 | 92.1469 | 92.2861 | 92.1377 | 92.1937 | 92.3228 | 92.2268 | 92.2071 | 92.3072 | 92.3156 | 92.3159 | 91.7246 | 91.3203 | 91.2689 | 90.0344 | 85.4155 | 85.1799 | 85.9189 | 84.6949 | 85.0917 | 85.0826 | 84.7601 | 84.739 | 84.7116 | 85.0187 | 85.0547 | 85.133 | 85.1456 | 85.4679 | 85.0241 | 85.0921 | 85.0008 | 84.9816 | 85.2372 | 84.9203 | 85.2242 | 84.9017 | 85.3368 | 85.0597 | 85.0197 | 84.9245 | 85.4372 |
| Xanthobacter oligotrophicus 23A | 93.229 | 93.3497 | 93.5475 | 93.2843 | 93.1956 | 93.3578 | 93.2986 | 93.2709 | 93.2769 | 100 | 96.3708 | 96.3786 | 92.4925 | 92.7055 | 92.3228 | 92.4582 | 92.4431 | 92.5584 | 92.6357 | 92.5467 | 92.5736 | 92.6164 | 92.5538 | 91.6923 | 91.928 | 90.4215 | 85.6856 | 85.2611 | 85.8252 | 84.6319 | 84.8085 | 84.7836 | 84.7386 | 84.7163 | 84.7305 | 85.0959 | 85.0882 | 85.0407 | 85.0767 | 85.1904 | 84.9513 | 84.8859 | 85.0686 | 85.0394 | 85.0693 | 84.9719 | 84.9564 | 84.9884 | 84.904 | 84.9167 | 84.8795 | 85.0877 | 85.0549 |
| Xanthobacter oligotrophicus 29k | 93.6583 | 93.4668 | 93.4142 | 93.6324 | 93.398 | 93.5408 | 93.6654 | 93.4458 | 93.3912 | 96.3421 | 100 | 99.994 | 92.696 | 92.8426 | 92.5492 | 92.5386 | 92.5748 | 92.5593 | 92.5586 | 92.7097 | 92.6801 | 92.6859 | 92.6597 | 91.6922 | 91.9261 | 90.7471 | 85.6728 | 85.1756 | 85.922 | 84.7434 | 85.2321 | 85.2737 | 84.873 | 84.6924 | 84.7249 | 85.076 | 85.0582 | 85.1507 | 85.2126 | 85.5375 | 85.1016 | 84.8594 | 84.965 | 84.9658 | 85.1077 | 85.0869 | 85.4366 | 85.0324 | 85.4823 | 84.9802 | 84.9895 | 84.9924 | 85.4588 |
| Xanthobacter oligotrophicus BF-7S | 93.6386 | 93.5624 | 93.3677 | 93.6883 | 93.4713 | 93.6127 | 93.6914 | 93.5456 | 93.5234 | 96.4313 | 99.9911 | 100 | 92.6302 | 92.782 | 92.5329 | 92.5116 | 92.612 | 92.5736 | 92.6096 | 92.6322 | 92.647 | 92.6711 | 92.6564 | 91.8108 | 92.0055 | 90.6692 | 85.6549 | 85.3236 | 85.9673 | 84.7559 | 85.2376 | 85.3005 | 84.925 | 84.7247 | 84.7261 | 85.1082 | 85.1174 | 85.1939 | 85.212 | 85.4885 | 85.1516 | 84.9142 | 84.992 | 85.0396 | 85.2537 | 85.087 | 85.4653 | 85.1518 | 85.6284 | 85.1052 | 85.0945 | 85.1089 | 85.4782 |
| Xanthobacter autotrophicus GZ29 | 92.1913 | 92.2094 | 92.3369 | 92.2383 | 92.0837 | 92.2106 | 92.2529 | 92.1639 | 92.1942 | 92.5236 | 92.7129 | 92.7353 | 100 | 99.0627 | 97.4733 | 97.5952 | 97.4891 | 97.4563 | 97.6026 | 97.4361 | 97.438 | 97.4349 | 91.5987 | 91.4589 | 91.3907 | 89.7915 | 85.2929 | 85.0002 | 85.7517 | 84.391 | 84.6427 | 84.5198 | 84.4886 | 84.4599 | 84.4621 | 84.8358 | 84.7329 | 84.8986 | 84.8323 | 84.9445 | 84.711 | 84.6526 | 84.7759 | 84.7775 | 84.8324 | 84.6163 | 84.7737 | 84.6786 | 85.1837 | 84.8277 | 84.7243 | 84.7156 | 84.992 |
| Xanthobacter autotrophicus 19/-/x | 92.3423 | 92.3961 | 92.3702 | 92.4123 | 92.3334 | 92.3305 | 92.3713 | 92.362 | 92.3884 | 92.7637 | 92.8789 | 92.913 | 99.0418 | 100 | 97.7622 | 97.751 | 97.763 | 97.7047 | 97.6431 | 97.6037 | 97.6315 | 97.634 | 91.8021 | 91.468 | 91.5459 | 89.9609 | 85.4177 | 85.0535 | 85.5745 | 84.5282 | 84.5882 | 84.6377 | 84.5745 | 84.4232 | 84.4706 | 84.8207 | 84.7919 | 84.9766 | 84.981 | 84.9419 | 84.8368 | 84.6351 | 84.7547 | 84.7633 | 84.7054 | 84.6195 | 84.7076 | 84.7656 | 84.7926 | 84.8394 | 84.7605 | 84.6936 | 84.8144 |
| Xanthobacter autotrophicus JW33 | 92.4213 | 92.3503 | 92.2366 | 92.3267 | 92.0957 | 92.3191 | 92.4401 | 92.1239 | 92.2036 | 92.485 | 92.6135 | 92.6562 | 97.5241 | 97.7427 | 100 | 99.4116 | 98.6573 | 98.0267 | 98.0033 | 98.2428 | 98.2485 | 98.2472 | 91.5901 | 91.7899 | 91.6977 | 89.9427 | 85.3717 | 84.9564 | 85.882 | 84.7864 | 85.3575 | 84.7993 | 84.6795 | 84.5401 | 84.5399 | 84.8024 | 84.8505 | 84.9787 | 84.9293 | 85.0346 | 84.7676 | 84.7706 | 84.7913 | 84.7605 | 85.1872 | 84.8521 | 84.8286 | 84.7427 | 85.075 | 84.9307 | 84.7225 | 84.8979 | 84.9982 |
| Xanthobacter autotrophicus 124X | 92.2363 | 92.3762 | 92.3237 | 92.2355 | 92.1321 | 92.2605 | 92.289 | 92.1838 | 92.2104 | 92.479 | 92.5149 | 92.5674 | 97.5778 | 97.7087 | 99.3095 | 100 | 98.8364 | 97.8478 | 98.2042 | 98.132 | 98.1594 | 98.1621 | 91.5777 | 91.6807 | 91.5484 | 89.8772 | 85.3825 | 84.8688 | 85.7069 | 84.5347 | 84.8843 | 84.5587 | 84.5134 | 84.5711 | 84.511 | 84.8839 | 84.8121 | 84.8812 | 84.9554 | 84.9093 | 84.8017 | 84.5873 | 84.726 | 84.8107 | 85.1639 | 84.8766 | 84.9282 | 84.8119 | 85.0986 | 84.7511 | 84.6606 | 84.801 | 84.9534 |
| Xanthobacter autotrophicus V0C-4 | 92.2377 | 92.5456 | 92.2951 | 92.1995 | 92.0501 | 92.2931 | 92.3482 | 92.2481 | 92.3852 | 92.5183 | 92.6953 | 92.651 | 97.4598 | 97.8042 | 98.6152 | 98.8095 | 100 | 97.8517 | 98.0544 | 98.1982 | 98.1797 | 98.2254 | 91.5026 | 91.4281 | 91.2509 | 89.9785 | 85.2244 | 85.1023 | 85.6029 | 84.3747 | 84.5967 | 84.6634 | 84.4683 | 84.431 | 84.3893 | 84.6584 | 84.7104 | 84.8673 | 84.8331 | 84.9854 | 84.793 | 84.5818 | 84.6614 | 84.6721 | 85.2736 | 84.6144 | 84.7733 | 84.5307 | 84.8354 | 84.715 | 84.561 | 84.6471 | 84.7866 |
| Xanthobacter autotrophicus NCIMB 11171 | 92.2839 | 92.3068 | 92.3949 | 92.2274 | 92.2156 | 92.2289 | 92.2456 | 92.2175 | 92.1731 | 92.541 | 92.5691 | 92.6409 | 97.4294 | 97.6732 | 98.0491 | 97.9277 | 98.021 | 100 | 98.5882 | 98.7349 | 98.7285 | 98.7158 | 91.9099 | 91.6613 | 91.3707 | 89.8953 | 85.4169 | 85.0715 | 85.6375 | 84.5826 | 84.5779 | 84.4858 | 84.5007 | 84.4682 | 84.448 | 84.8332 | 84.7758 | 84.9032 | 84.8494 | 84.7456 | 84.687 | 84.5823 | 84.57 | 84.5071 | 84.7004 | 84.7244 | 84.6522 | 84.5352 | 84.6467 | 84.6048 | 84.6012 | 84.7454 | 84.6603 |
| Xanthobacter autotrophicus CCUG 44692 | 92.2213 | 92.2872 | 92.3364 | 92.1798 | 92.0577 | 92.3411 | 92.2313 | 92.0508 | 92.1313 | 92.5805 | 92.5671 | 92.634 | 97.6341 | 97.6139 | 97.9919 | 98.2905 | 98.0521 | 98.6744 | 100 | 99.1892 | 99.1649 | 99.1827 | 91.7775 | 91.2274 | 91.5328 | 89.9415 | 85.3134 | 85.1567 | 85.6959 | 84.4198 | 84.4376 | 84.4785 | 84.5052 | 84.5406 | 84.5433 | 84.6733 | 84.7566 | 84.7971 | 84.9461 | 84.814 | 84.6571 | 84.6631 | 84.5979 | 84.517 | 84.7824 | 84.7407 | 84.7234 | 84.7243 | 84.5829 | 84.6611 | 84.7282 | 84.7517 | 84.6921 |
| Xanthobacter viscosus 7d | 92.2875 | 92.3439 | 92.3331 | 92.3853 | 92.2654 | 92.3367 | 92.4222 | 92.2599 | 92.3745 | 92.5633 | 92.7461 | 92.885 | 97.4201 | 97.6231 | 98.1604 | 98.1432 | 98.2428 | 98.725 | 99.2338 | 100 | 99.9945 | 99.9955 | 91.7951 | 91.3901 | 91.2955 | 90.069 | 85.3628 | 85.0201 | 85.7236 | 84.479 | 84.7266 | 84.6571 | 84.4567 | 84.499 | 84.5392 | 84.8247 | 84.8016 | 84.9733 | 84.9569 | 85.0153 | 84.8003 | 84.7054 | 84.6755 | 84.615 | 84.9152 | 84.7305 | 84.8394 | 84.6089 | 84.7859 | 84.8218 | 84.6505 | 84.7702 | 84.8995 |
| Xanthobacter autotrophicus 7C SF | 92.3513 | 92.4449 | 92.3689 | 92.4139 | 92.2456 | 92.3904 | 92.3797 | 92.2575 | 92.3823 | 92.5832 | 92.7576 | 92.8159 | 97.429 | 97.631 | 98.1154 | 98.1341 | 98.2089 | 98.7847 | 99.229 | 99.9976 | 100 | 99.9931 | 91.7652 | 91.4056 | 91.3978 | 90.0075 | 85.3045 | 84.9665 | 85.6839 | 84.526 | 84.7177 | 84.7513 | 84.5039 | 84.5804 | 84.5497 | 84.8503 | 84.8508 | 84.9032 | 85.019 | 85.0354 | 84.7001 | 84.7299 | 84.6644 | 84.6831 | 84.9093 | 84.6714 | 84.8814 | 84.6954 | 84.7553 | 84.7476 | 84.737 | 84.7598 | 84.9093 |
| Xanthobacter autotrophicus 7C | 92.3252 | 92.45 | 92.402 | 92.4328 | 92.3083 | 92.3873 | 92.395 | 92.2798 | 92.3867 | 92.6039 | 92.7962 | 92.8945 | 97.4138 | 97.6246 | 98.1441 | 98.1346 | 98.2037 | 98.7438 | 99.217 | 99.9977 | 99.997 | 100 | 91.8056 | 91.4237 | 91.4234 | 90.0172 | 85.3183 | 84.9977 | 85.6654 | 84.5026 | 84.738 | 84.7612 | 84.5202 | 84.5339 | 84.4695 | 84.8488 | 84.8172 | 84.9344 | 84.9699 | 85.0706 | 84.7546 | 84.7419 | 84.6856 | 84.7103 | 85.0004 | 84.6617 | 84.878 | 84.7079 | 84.664 | 84.7782 | 84.6905 | 84.7967 | 84.9211 |
| Xanthobacter sp. AM5 | 91.9866 | 91.9836 | 92.0838 | 91.9372 | 91.7654 | 91.994 | 91.9271 | 91.91 | 91.8899 | 92.6646 | 92.7016 | 92.7052 | 91.7134 | 91.8096 | 91.6542 | 91.584 | 91.6444 | 91.9453 | 91.8073 | 91.7266 | 91.7153 | 91.7371 | 100 | 93.6026 | 93.4252 | 91.1402 | 85.9518 | 85.3764 | 85.8159 | 84.8606 | 84.7385 | 84.9389 | 84.8374 | 84.8959 | 84.9511 | 85.2112 | 85.1635 | 85.3059 | 85.3269 | 85.2443 | 85.1147 | 85.0591 | 84.9515 | 84.9503 | 84.9723 | 85.1309 | 85.0195 | 85.0971 | 85.118 | 85.0056 | 85.5524 | 85.0891 | 84.9134 |
| Xanthobacter toluenivorans T101 | 91.2433 | 91.3065 | 91.421 | 91.2326 | 91.1313 | 91.3017 | 91.3286 | 91.1689 | 91.1848 | 91.7301 | 91.82 | 91.9027 | 91.4281 | 91.5104 | 91.6174 | 91.6349 | 91.3858 | 91.557 | 91.3215 | 91.3877 | 91.3315 | 91.3717 | 93.4816 | 100 | 97.6009 | 90.2527 | 85.709 | 85.0972 | 85.8852 | 84.7991 | 85.2649 | 84.7315 | 84.6367 | 84.6853 | 84.7162 | 84.8953 | 84.8619 | 84.8376 | 85.0489 | 84.8904 | 84.7779 | 84.6126 | 84.6212 | 84.6979 | 85.3821 | 84.7011 | 84.7899 | 84.8306 | 84.8699 | 84.7474 | 84.6832 | 84.7082 | 84.78 |
| Xanthobacter toluenivorans T102 | 91.3362 | 91.2781 | 91.3766 | 91.2632 | 91.243 | 91.3578 | 91.2276 | 91.1905 | 91.2672 | 91.8821 | 91.9493 | 92.0433 | 91.4194 | 91.5826 | 91.6141 | 91.5911 | 91.2639 | 91.3782 | 91.5038 | 91.4335 | 91.4193 | 91.4791 | 93.3833 | 97.5654 | 100 | 90.1917 | 85.8596 | 85.0308 | 85.7934 | 85.1416 | 85.9481 | 84.8017 | 84.8271 | 84.6484 | 84.6116 | 84.8901 | 84.9211 | 85.0312 | 85.0562 | 85.1202 | 84.9867 | 84.712 | 84.6285 | 84.7559 | 85.2749 | 85.3258 | 84.8265 | 84.8602 | 84.9627 | 84.8966 | 84.7913 | 84.6534 | 84.8497 |
| Xanthobacter sp. AM3 | 90.4126 | 90.1839 | 90.1187 | 90.3703 | 90.0864 | 90.2361 | 90.3434 | 90.1956 | 90.1923 | 90.4881 | 90.8293 | 90.9031 | 89.9523 | 90.0348 | 89.9184 | 89.8525 | 89.9783 | 89.9227 | 89.9298 | 89.9366 | 89.9357 | 89.9748 | 91.3629 | 90.4248 | 90.3069 | 100 | 86.0969 | 85.5782 | 86.205 | 84.9901 | 85.4234 | 85.5484 | 84.9502 | 84.9535 | 84.9342 | 85.2481 | 85.234 | 85.4204 | 85.3483 | 85.7284 | 85.239 | 85.1592 | 85.1515 | 85.1026 | 85.3016 | 85.2729 | 85.685 | 85.1589 | 85.49 | 85.1204 | 85.6994 | 85.054 | 85.5031 |
| Xanthobacter dioxanivorans YN2 | 85.5121 | 85.4425 | 85.3754 | 85.4888 | 85.6616 | 85.4305 | 85.44 | 85.4985 | 85.3042 | 85.4387 | 85.5161 | 85.6658 | 85.0728 | 85.2198 | 85.2352 | 85.2495 | 85.1223 | 85.3019 | 85.2342 | 85.2649 | 85.2253 | 85.2437 | 85.966 | 85.5869 | 85.8103 | 85.8439 | 100 | 85.8655 | 85.5632 | 84.7829 | 84.6579 | 84.6201 | 84.4772 | 84.7127 | 84.6946 | 85.1563 | 85.1964 | 84.9626 | 85.0363 | 84.885 | 84.9064 | 84.6661 | 84.8973 | 84.9177 | 85.0953 | 85.1194 | 85.026 | 85.0932 | 84.9612 | 84.8247 | 85.0582 | 85.0927 | 85.0553 |
| Xanthobacter nonsaccharivorans 14g | 85.212 | 85.217 | 85.143 | 85.154 | 85.1889 | 85.1556 | 85.2101 | 85.2203 | 85.112 | 85.2211 | 85.2346 | 85.2905 | 84.8694 | 84.9963 | 84.871 | 84.8893 | 85.0438 | 84.9978 | 85.0806 | 85.0008 | 85.0508 | 85.0083 | 85.4318 | 85.0797 | 85.0976 | 85.3977 | 85.9085 | 100 | 85.5569 | 84.77 | 84.871 | 84.7723 | 84.5426 | 84.8346 | 84.8809 | 85.1155 | 85.1382 | 85.1632 | 85.0697 | 85.2431 | 85.0142 | 85.0508 | 85.1539 | 85.1124 | 84.9931 | 85.0935 | 85.174 | 85.1126 | 85.3111 | 85.0247 | 85.2073 | 85.0097 | 85.1408 |
| Xanthobacter lutulentifluminis V3C-3 | 85.9177 | 85.7988 | 85.794 | 86.0222 | 86.0067 | 86.0508 | 86.0415 | 85.9673 | 85.9513 | 85.9363 | 85.9654 | 85.9564 | 85.7046 | 85.5717 | 85.9057 | 85.638 | 85.5897 | 85.5674 | 85.6213 | 85.5131 | 85.5273 | 85.5332 | 85.8209 | 85.8186 | 85.8047 | 86.1218 | 85.5287 | 85.6906 | 100 | 85.358 | 85.3501 | 85.4024 | 85.4567 | 85.4191 | 85.4134 | 85.6534 | 85.7262 | 85.6947 | 85.6907 | 85.7402 | 85.567 | 85.609 | 85.6162 | 85.6183 | 85.8327 | 85.6361 | 85.6609 | 85.623 | 85.8797 | 85.6477 | 85.5904 | 85.6504 | 85.9823 |
| [illegible] | [illegible] | [illegible] | [illegible] | [illegible] | [illegible] | [illegible] | [illegible] | [illegible] | [illegible] | [illegible] | [illegible] | [illegible] | [illegible] | [illegible] | [illegible] | [illegible] | [illegible] | [illegible] | [illegible] | [illegible] | [illegible] | [illegible] | [illegible] | [illegible] | [illegible] | [illegible] | [illegible] | [illegible] | [illegible] | [illegible] | [illegible] | [illegible] | [illegible] | [illegible] | [illegible] | [illegible] | [illegible] | [illegible] | [illegible] | [illegible] | [illegible] | [illegible] | [illegible] | [illegible] | [illegible] | [illegible] | [illegible] | [illegible] | [illegible] | [illegible] | [illegible] | [illegible] | [illegible] |

**Supplementary Table 4 –Xanthobacter and Roseixanthobacter gen. nov. strains can be largely separated into our defined species clusters using digital DNA-DNA Hybridization (dDDH) values calculated using the Genome-to-Genome Distance Calculator (v3) with a 70% species level threshold.** Values above the 70% dDDH score species threshold are colored in blue while those below the species threshold are colored in red. With dDDH values alone, unlike with average nucleotide value thresholds at 95%, strain 23A is no longer within the species level boundary for *X. oligotrophicus*, and strains V8C-8 and 126 are no longer within the species level boundary for *X. aminoxidans*.

| | Xanthobacter versatilis CB6 | Xanthobacter versatilis V7C-4 | Xanthobacter versatilis V1C-2 | Xanthobacter versatilis V2C-9 | Xanthobacter versatilis Py2 | Xanthobacter versatilis V1C-6B | Xanthobacter versatilis V7C-1B | Xanthobacter versatilis V3C-2 | Xanthobacter oligotrophicus 23A | Xanthobacter oligotrophicus 29k | Xanthobacter oligotrophicus BF-7S | Xanthobacter autotrophicus GZ29 | Xanthobacter autotrophicus 19/-/x | Xanthobacter autotrophicus JW33 | Xanthobacter autotrophicus 124X | Xanthobacter autotrophicus V0C-4 | Xanthobacter autotrophicus NCIMB 11171 | Xanthobacter autotrophicus CCUG 44692 | Xanthobacter viscosus 7d | Xanthobacter autotrophicus 7C SF | Xanthobacter autotrophicus 7C | Xanthobacter sp. AM5 | Xanthobacter toluenivorans T101 | Xanthobacter toluenivorans T102 | Xanthobacter sp. AM3 | Xanthobacter dioxanivorans YN2 | Xanthobacter nonsaccharivorans 14g | Xanthobacter lutulentifluminis V3C-3 | Xanthobacter wiegelii CB2 | Xanthobacter wiegelii RH 10 | Xanthobacter wiegelii WS4821 | Xanthobacter wiegelii NCIMB 11399 |
|---|---|---|---|---|---|---|---|---|---|---|---|---|---|---|---|---|---|---|---|---|---|---|---|---|---|---|---|---|---|---|---|---|
| Xanthobacter versatilis CB6 | 100 | 88.7 | 85.7 | 87.8 | 85 | 85 | 85.9 | 87.5 | 47.4 | 48.7 | 48.7 | 42.9 | 43.4 | 43.1 | 43 | 43.2 | 42.9 | 43.2 | 43.4 | 43.4 | 43.4 | 42.3 | 39.4 | 39.8 | 36.6 | 26.8 | 25.9 | 26.3 | 25 | 26.1 | 26 | 25.2 |
| Xanthobacter versatilis V7C-4 | 88.7 | 100 | 86.8 | 88.3 | 84.7 | 88 | 86.3 | 85.4 | 48.1 | 48.8 | 48.8 | 42.9 | 43.5 | 43.1 | 43.3 | 44 | 43.4 | 43.4 | 43.5 | 43.5 | 43.5 | 42.5 | 39.6 | 39.7 | 36.2 | 26.8 | 26.1 | 26.3 | 25 | 25.7 | 25.6 | 25 |
| Xanthobacter versatilis V1C-2 | 85.7 | 86.8 | 100 | 86.3 | 84.9 | 86.4 | 86.3 | 85.2 | 48.9 | 48 | 48 | 43.1 | 43.6 | 43.1 | 43.4 | 43.5 | 43.5 | 43.6 | 43.6 | 43.6 | 43.6 | 42.5 | 39.6 | 39.8 | 35.8 | 26.6 | 26 | 26.1 | 24.8 | 25 | 24.9 | 24.8 |
| Xanthobacter versatilis V2C-9 | 87.8 | 88.3 | 86.3 | 100 | 83.9 | 85.8 | 86.2 | 84.3 | 47.6 | 48.9 | 48.9 | 42.9 | 43.5 | 42.9 | 42.8 | 43.1 | 43 | 43.2 | 43.5 | 43.5 | 43.5 | 42.3 | 39.6 | 39.4 | 36.6 | 26.7 | 26 | 26.3 | 25 | 25.9 | 26 | 25 |
| Xanthobacter versatilis Py2 | 85 | 84.7 | 84.9 | 83.9 | 100 | 83.3 | 83.2 | 83 | 47.3 | 48.2 | 48.1 | 42.4 | 43.3 | 43 | 42.8 | 43 | 43 | 42.6 | 43.2 | 43.2 | 43.2 | 41.8 | 39.1 | 39.6 | 36 | 27.1 | 25.9 | 26.5 | 26.1 | 26.5 | 25.7 | 24.9 |
| Xanthobacter versatilis V1C-6B | 85 | 88 | 86.4 | 85.8 | 83.3 | 100 | 85.6 | 84.2 | 48.2 | 48.7 | 48.6 | 43.1 | 43.4 | 43.2 | 43.1 | 43.4 | 43.4 | 43.5 | 43.5 | 43.5 | 43.5 | 42.4 | 39.6 | 39.8 | 36 | 26.7 | 25.9 | 26.5 | 25 | 25.7 | 25.6 | 25 |
| Xanthobacter versatilis V7C-1B | 85.9 | 86.3 | 86.3 | 86.2 | 83.2 | 85.6 | 100 | 83.7 | 47.7 | 49.2 | 49.2 | 43.1 | 43.5 | 43.4 | 42.9 | 43.3 | 43.1 | 43.3 | 43.4 | 43.4 | 43.5 | 42.2 | 39.5 | 39.4 | 36.5 | 26.8 | 26 | 26.4 | 25 | 25.8 | 26.2 | 24.9 |
| Xanthobacter versatilis V3C-2 | 87.5 | 85.4 | 85.2 | 84.3 | 83 | 84.2 | 83.7 | 100 | 47.6 | 48.5 | 48.5 | 42.6 | 43.4 | 42.8 | 42.5 | 43.5 | 42.7 | 42.8 | 43.4 | 43.3 | 43.4 | 42.1 | 39.1 | 39.1 | 36.3 | 26.9 | 26.1 | 26.3 | 25.1 | 26 | 25.8 | 25 |
| Xanthobacter oligotrophicus 23A | 47.4 | 48.1 | 48.9 | 47.6 | 47.3 | 48.2 | 47.7 | 47.6 | 100 | 66.3 | 66.3 | 44.4 | 45 | 43.4 | 43.8 | 44 | 44.3 | 44.7 | 44.6 | 44.6 | 44.6 | 44.9 | 41.5 | 41.7 | 36.8 | 26.7 | 26.2 | 26.1 | 24.8 | 24.9 | 24.9 | 24.9 |
| Xanthobacter oligotrophicus 29k | 48.7 | 48.8 | 48 | 48.9 | 48.2 | 48.7 | 49.2 | 48.5 | 66.3 | 100 | 100 | 44.7 | 45.3 | 44 | 44 | 44.4 | 44.4 | 44.6 | 44.9 | 44.9 | 44.9 | 45 | 41.5 | 41.8 | 37.6 | 26.9 | 26.1 | 26.2 | 24.9 | 25.7 | 25.9 | 25 |
| Xanthobacter oligotrophicus BF-7S | 48.7 | 48.8 | 48 | 48.9 | 48.1 | 48.6 | 49.2 | 48.5 | 66.3 | 100 | 100 | 44.8 | 45.3 | 44 | 44 | 44.4 | 44.4 | 44.6 | 44.9 | 44.9 | 44.9 | 45 | 41.5 | 41.8 | 37.6 | 26.9 | 26.1 | 26.2 | 24.9 | 25.7 | 25.9 | 24.9 |
| Xanthobacter autotrophicus GZ29 | 42.9 | 42.9 | 43.1 | 42.9 | 42.4 | 43.1 | 43.1 | 42.6 | 44.4 | 44.7 | 44.8 | 100 | 91.2 | 75.3 | 76.5 | 76.1 | 75.8 | 77.3 | 76.2 | 76.2 | 76.3 | 41.8 | 40.2 | 40.3 | 35.7 | 26.7 | 25.9 | 26.3 | 24.8 | 25.1 | 25.1 | 24.9 |
| Xanthobacter autotrophicus 19/-/x | 43.4 | 43.5 | 43.6 | 43.5 | 43.3 | 43.4 | 43.5 | 43.4 | 45 | 45.3 | 45.3 | 91.2 | 100 | 77.7 | 77.7 | 78.1 | 77.3 | 77.8 | 77.8 | 77.9 | 78 | 42 | 40.1 | 40.4 | 35.8 | 26.7 | 26 | 26.1 | 24.9 | 25.1 | 25.1 | 25 |
| Xanthobacter autotrophicus JW33 | 43.1 | 43.1 | 43.1 | 42.9 | 43 | 43.2 | 43.4 | 42.8 | 43.4 | 44 | 44 | 75.3 | 77.7 | 100 | 93.6 | 87.8 | 81.3 | 82.4 | 82.6 | 82.6 | 82.6 | 41.3 | 41.2 | 40.9 | 35.7 | 26.8 | 25.9 | 26.6 | 25.4 | 26.7 | 25.5 | 25.4 |
| Xanthobacter autotrophicus 124X | 43 | 43.3 | 43.4 | 42.8 | 42.8 | 43.1 | 42.9 | 42.5 | 43.8 | 44 | 44 | 76.5 | 77.7 | 93.6 | 100 | 88.7 | 80.2 | 83.7 | 82.1 | 82.1 | 82.2 | 40.9 | 40.8 | 40.7 | 35.6 | 26.7 | 25.8 | 26.3 | 25.2 | 26.1 | 25.4 | 25.1 |
| Xanthobacter autotrophicus V0C-4 | 43.2 | 44 | 43.5 | 43.1 | 43 | 43.4 | 43.3 | 43.5 | 44 | 44.4 | 44.4 | 76.1 | 78.1 | 87.8 | 88.7 | 100 | 81.2 | 81.6 | 82.9 | 82.9 | 83 | 41.5 | 40.5 | 39.8 | 35.8 | 26.5 | 26 | 26.1 | 25 | 25.6 | 25.5 | 25.2 |
| Xanthobacter autotrophicus NCIMB 11171 | 42.9 | 43.4 | 43.5 | 43 | 43 | 43.4 | 43.1 | 42.7 | 44.3 | 44.4 | 44.4 | 75.8 | 77.3 | 81.3 | 80.2 | 81.2 | 100 | 87.6 | 87.9 | 88 | 88 | 42.2 | 40.6 | 40.2 | 35.8 | 26.6 | 25.9 | 25.9 | 25.2 | 25.4 | 25.1 | 25 |
| Xanthobacter autotrophicus CCUG 44692 | 43.2 | 43.4 | 43.6 | 43.2 | 42.6 | 43.5 | 43.3 | 42.8 | 44.7 | 44.6 | 44.6 | 77.3 | 77.8 | 82.4 | 83.7 | 81.6 | 87.6 | 100 | 93.7 | 93.7 | 93.8 | 42.1 | 40.2 | 40.4 | 36 | 26.7 | 26 | 26 | 24.9 | 25.1 | 25.2 | 25.1 |
| Xanthobacter viscosus 7d | 43.4 | 43.5 | 43.6 | 43.5 | 43.2 | 43.5 | 43.4 | 43.4 | 44.6 | 44.9 | 44.9 | 76.2 | 77.8 | 82.6 | 82.1 | 82.9 | 87.9 | 93.7 | 100 | 100 | 100 | 42.1 | 40 | 40 | 36 | 26.7 | 26 | 26 | 25 | 25.4 | 25.5 | 25.1 |
| Xanthobacter autotrophicus 7C SF | 43.4 | 43.5 | 43.6 | 43.5 | 43.2 | 43.5 | 43.4 | 43.3 | 44.6 | 44.9 | 44.9 | 76.2 | 77.9 | 82.6 | 82.1 | 82.9 | 88 | 93.7 | 100 | 100 | 100 | 42 | 40 | 40 | 36 | 26.7 | 26.1 | 26 | 25 | 25.4 | 25.5 | 25.1 |
| Xanthobacter autotrophicus 7C | 43.4 | 43.5 | 43.6 | 43.5 | 43.2 | 43.5 | 43.5 | 43.4 | 44.6 | 44.9 | 44.9 | 76.3 | 78 | 82.6 | 82.2 | 83 | 88 | 93.8 | 100 | 100 | 100 | 42 | 40 | 40 | 36 | 26.7 | 26.1 | 26 | 25 | 25.4 | 25.5 | 25.1 |
| Xanthobacter sp. AM5 | 42.3 | 42.5 | 42.5 | 42.3 | 41.8 | 42.4 | 42.2 | 42.1 | 44.9 | 45 | 45 | 41.8 | 42 | 41.3 | 40.9 | 41.5 | 42.2 | 42.1 | 42.1 | 42 | 42 | 100 | 48.3 | 47.9 | 39.5 | 27.7 | 26.4 | 26.6 | 25.4 | 25.4 | 25.5 | 25.3 |
| Xanthobacter toluenivorans T101 | 39.4 | 39.6 | 39.6 | 39.6 | 39.1 | 39.6 | 39.5 | 39.1 | 41.5 | 41.5 | 41.5 | 40.2 | 40.1 | 41.2 | 40.8 | 40.5 | 40.6 | 40.2 | 40 | 40 | 40 | 48.3 | 100 | 75.4 | 36.8 | 27.4 | 25.9 | 26.6 | 25.5 | 26.7 | 25.5 | 25.4 |
| Xanthobacter toluenivorans T102 | 39.8 | 39.7 | 39.8 | 39.4 | 39.6 | 39.8 | 39.4 | 39.1 | 41.7 | 41.8 | 41.8 | 40.3 | 40.4 | 40.9 | 40.7 | 39.8 | 40.2 | 40.4 | 40 | 40 | 40 | 47.9 | 75.4 | 100 | 36.6 | 27.7 | 26 | 26.6 | 26.4 | 27.9 | 25.5 | 25.6 |
| Xanthobacter sp. AM3 | 36.6 | 36.2 | 35.8 | 36.6 | 36 | 36 | 36.5 | 36.3 | 36.8 | 37.6 | 37.6 | 35.7 | 35.8 | 35.7 | 35.6 | 35.8 | 35.8 | 36 | 36 | 36 | 36 | 39.5 | 36.8 | 36.6 | 100 | 27.4 | 26.5 | 26.8 | 25.5 | 26.5 | 26.6 | 25.5 |
| Xanthobacter dioxanivorans YN2 | 26.8 | 26.8 | 26.6 | 26.7 | 27.1 | 26.7 | 26.8 | 26.9 | 26.7 | 26.9 | 26.9 | 26.7 | 26.7 | 26.8 | 26.7 | 26.5 | 26.6 | 26.7 | 26.7 | 26.7 | 26.7 | 27.7 | 27.4 | 27.7 | 27.4 | 100 | 26.6 | 26 | 25.7 | 25.6 | 25.4 | 25.2 |
| Xanthobacter nonsaccharivorans 14g | 25.9 | 26.1 | 26 | 26 | 25.9 | 25.9 | 26 | 26.1 | 26.2 | 26.1 | 26.1 | 25.9 | 26 | 25.9 | 25.8 | 26 | 25.9 | 26 | 26 | 26.1 | 26.1 | 26.4 | 25.9 | 26 | 26.5 | 26.6 | 100 | 26.2 | 25.5 | 25.6 | 25.3 | 25.3 |
| Xanthobacter lutulentifluminis V3C-3 | 26.3 | 26.3 | 26.1 | 26.3 | 26.5 | 26.5 | 26.4 | 26.3 | 26.1 | 26.2 | 26.2 | 26.3 | 26.1 | 26.6 | 26.3 | 26.1 | 25.9 | 26 | 26 | 26 | 26 | 26.6 | 26.6 | 26.6 | 26.8 | 26 | 26.2 | 100 | 25.6 | 25.7 | 25.7 | 25.5 |
| Xanthobacter wiegelii CB2 | 25 | 25 | 24.8 | 25 | 26.1 | 25 | 25 | 25.1 | 24.8 | 24.9 | 24.9 | 24.8 | 24.9 | 25.4 | 25.2 | 25 | 25.2 | 24.9 | 25 | 25 | 25 | 25.4 | 25.5 | 26.4 | 25.5 | 25.7 | 25.5 | 25.6 | 100 | 86.3 | 86.7 | 87.1 |
| Xanthobacter wiegelii RH 10 | 26.1 | 25.7 | 25 | 25.9 | 26.5 | 25.7 | 25.8 | 26 | 24.9 | 25.7 | 25.7 | 25.1 | 25.1 | 26.7 | 26.1 | 25.6 | 25.4 | 25.1 | 25.4 | 25.4 | 25.4 | 25.4 | 26.7 | 27.9 | 26.5 | 25.6 | 25.6 | 25.7 | 86.3 | 100 | 86.9 | 85.5 |
| Xanthobacter wiegelii WS4821 | 26 | 25.6 | 24.9 | 26 | 25.7 | 25.6 | 26.2 | 25.8 | 24.9 | 25.9 | 25.9 | 25.1 | 25.1 | 25.5 | 25.4 | 25.5 | 25.1 | 25.2 | 25.5 | 25.5 | 25.5 | 25.5 | 25.5 | 25.5 | 26.6 | 25.4 | 25.3 | 25.7 | 86.7 | 86.9 | 100 | 85.1 |
| Xanthobacter wiegelii NCIMB 11399 | 25.2 | 25 | 24.8 | 25 | 24.9 | 25 | 24.9 | 25 | 24.9 | 25 | 24.9 | 24.9 | 25 | 25.4 | 25.1 | 25.2 | 25 | 25.1 | 25.1 | 25.1 | 25.1 | 25.3 | 25.4 | 25.6 | 25.5 | 25.2 | 25.3 | 25.5 | 87.1 | 85.5 | 85.1 | 100 |

| | Xanthobacter versatilis CB6 | Xanthobacter versatilis V7C-4 | Xanthobacter versatilis V1C-2 | Xanthobacter versatilis V2C-9 | Xanthobacter versatilis Py2 | Xanthobacter versatilis V1C-6B | Xanthobacter versatilis V7C-1B | Xanthobacter versatilis V3C-2 | Xanthobacter oligotrophicus 23A | Xanthobacter oligotrophicus 29k | Xanthobacter oligotrophicus BF-7S | Xanthobacter autotrophicus GZ29 | Xanthobacter autotrophicus 19/-/x | Xanthobacter autotrophicus JW33 | Xanthobacter autotrophicus 124X | Xanthobacter autotrophicus V0C-4 | Xanthobacter autotrophicus NCIMB 11171 | Xanthobacter autotrophicus CCUG 44692 | Xanthobacter viscosus 7d | Xanthobacter autotrophicus 7C SF | Xanthobacter autotrophicus 7C | Xanthobacter sp. AM5 |
|---|---|---|---|---|---|---|---|---|---|---|---|---|---|---|---|---|---|---|---|---|---|---|
| Xanthobacter sp. SG518 | 24.9 | 24.8 | 24.7 | 24.8 | 24.9 | 24.8 | 24.8 | 24.9 | 24.8 | 24.8 | 24.8 | 24.8 | 24.7 | 24.9 | 24.9 | 24.8 | 24.8 | 24.8 | 24.7 | 24.7 | 24.8 | 25.2 |
| Xanthobacter sp. SG563 | 24.9 | 24.8 | 24.7 | 24.8 | 24.9 | 24.8 | 24.8 | 24.9 | 24.8 | 24.8 | 24.8 | 24.8 | 24.7 | 24.9 | 24.9 | 24.8 | 24.8 | 24.8 | 24.7 | 24.7 | 24.8 | 25.2 |
| Xanthobacter aminoxidans CB3 | 25.1 | 25.1 | 25.1 | 25.1 | 25 | 25 | 25 | 25 | 25 | 25.1 | 25.1 | 25 | 25.1 | 25.1 | 25.2 | 25 | 25.1 | 25.1 | 25.1 | 25.1 | 25.1 | 25.6 |
| Xanthobacter aminoxidans CB5 | 25.1 | 25.1 | 25.1 | 25.1 | 25 | 25 | 25 | 25 | 25 | 25.1 | 25.1 | 25 | 25.1 | 25.1 | 25.2 | 25 | 25.1 | 25.1 | 25.1 | 25.1 | 25.1 | 25.6 |
| Xanthobacter aminoxidans 91 | 25.3 | 25.2 | 25.1 | 25.2 | 25.3 | 25.1 | 25.2 | 25.2 | 25.1 | 25.2 | 25.2 | 25 | 25.1 | 25.1 | 25.3 | 25.1 | 25.2 | 25.2 | 25.2 | 25.2 | 25.2 | 25.6 |
| Xanthobacter aminoxidans 14a | 25.3 | 25.4 | 26 | 25.2 | 25.2 | 25.4 | 25.2 | 25.3 | 25.2 | 25.2 | 25.2 | 24.9 | 25 | 25.3 | 25.2 | 25.2 | 25.3 | 25.4 | 25.3 | 25.3 | 25.3 | 25.7 |
| Xanthobacter aminoxidans V7C-9 | 26.5 | 25.9 | 25.1 | 26.3 | 26 | 25.8 | 26.2 | 26.2 | 25.1 | 26.1 | 26.1 | 25.3 | 25.1 | 25.6 | 25.5 | 25.6 | 25.2 | 25.3 | 25.5 | 25.5 | 25.5 | 25.7 |
| Xanthobacter aminoxidans V8C-8 | 25.5 | 25.2 | 25 | 25.2 | 25.2 | 25.2 | 25.2 | 25.2 | 25.1 | 25.2 | 25.2 | 24.8 | 24.9 | 25.1 | 25 | 24.9 | 24.8 | 25 | 25 | 25 | 25 | 25.4 |
| Xanthobacter aminoxidans 126 | 25.3 | 25.2 | 25.1 | 25.2 | 25.2 | 25.2 | 25.2 | 25.2 | 25.1 | 25.1 | 25.1 | 24.9 | 24.8 | 25 | 25.1 | 25 | 25 | 25 | 25 | 25 | 25 | 25.6 |
| Xanthobacter flavus SS-20 | 25.3 | 25.3 | 25.2 | 25.2 | 25.3 | 25.1 | 25.2 | 25.3 | 25.2 | 25.2 | 25.2 | 25.1 | 25.1 | 25 | 25.2 | 25.1 | 25.1 | 25.1 | 25 | 25 | 25 | 25.5 |
| Xanthobacter flavus SS-22 | 25.3 | 25.2 | 25.2 | 25.2 | 25.3 | 25.1 | 25.2 | 25.2 | 25.2 | 25.2 | 25.2 | 25.1 | 25.1 | 25 | 25.2 | 25.2 | 25.1 | 25.1 | 25 | 25 | 25 | 25.5 |
| Xanthobacter flavus GJ10 | 26.2 | 26.1 | 25.7 | 26.1 | 26.1 | 26.7 | 26 | 26.1 | 25.6 | 25.9 | 25.9 | 25.8 | 25.5 | 26.1 | 26.4 | 26.8 | 25.8 | 25.8 | 26 | 26 | 26 | 26.1 |
| Xanthobacter flavus R-10 | 25.2 | 25.1 | 25.1 | 25.2 | 25.8 | 25.1 | 25.1 | 25.1 | 25.1 | 25.1 | 25.1 | 25 | 25 | 25.2 | 25.4 | 25.1 | 25.3 | 25.2 | 25.1 | 25.1 | 25.1 | 25.8 |
| Xanthobacter flavus V4C-10 | 26 | 25.7 | 25.1 | 26 | 25.9 | 25.6 | 26 | 25.9 | 25.1 | 26 | 26 | 25.1 | 25 | 25.2 | 25.4 | 25.3 | 25.1 | 25.2 | 25.4 | 25.4 | 25.4 | 25.6 |
| Xanthobacter flavus V4C-7 | 25.1 | 25 | 24.9 | 25 | 25.2 | 24.9 | 25 | 24.9 | 25 | 25.1 | 25.2 | 25 | 25.1 | 24.9 | 25.1 | 24.9 | 25 | 25.1 | 25 | 25 | 25 | 25.6 |
| Xanthobacter flavus 301 | 26 | 25.8 | 25.1 | 25.9 | 26.1 | 25.8 | 26.1 | 26 | 25 | 26.3 | 26.3 | 25.8 | 25.1 | 25.5 | 25.7 | 25.5 | 25 | 25.1 | 25.3 | 25.3 | 25.3 | 25.7 |
| Xanthobacter flavus YC-JY1 | 25.5 | 25.2 | 25.1 | 25.4 | 25.3 | 25.2 | 25.3 | 25.4 | 25 | 25.3 | 25.3 | 25.1 | 25 | 25.3 | 25.1 | 25.1 | 24.9 | 25 | 25 | 25.1 | 25.1 | 25.5 |
| Xanthobacter flavus V4C-9 | 25.1 | 25.1 | 25 | 25.1 | 25.2 | 25 | 25.1 | 25 | 25.1 | 25.2 | 25.2 | 25 | 25 | 25 | 25.1 | 25.1 | 25 | 25.2 | 25.1 | 25.1 | 25.1 | 26.8 |
| Xanthobacter flavus SG618 | 25.1 | 25 | 25 | 25 | 25 | 24.9 | 25 | 24.9 | 25 | 25 | 25 | 25 | 25 | 24.9 | 25.1 | 24.9 | 24.9 | 25.1 | 25.1 | 25.1 | 25.1 | 25.5 |
| Xanthobacter flavus V2C-3 | 26.2 | 25.8 | 24.9 | 26 | 25.8 | 25.9 | 26.2 | 26.1 | 25.1 | 26 | 26 | 25.5 | 25.3 | 25.6 | 25.8 | 25.5 | 25.1 | 25.2 | 25.5 | 25.5 | 25.5 | 25.5 |
| Xanthobacter gandavensis NFM-26 | 23.4 | 23.4 | 22.9 | 23.2 | 23.7 | 23.6 | 24.1 | 23.4 | 22.9 | 23.2 | 23.2 | 23.3 | 22.9 | 24.3 | 23.2 | 23 | 22.7 | 22.8 | 22.9 | 22.9 | 22.9 | 23.3 |
| Xanthobacter gandavensis NM-25 | 23.4 | 23.4 | 23 | 23.2 | 23.7 | 23.6 | 24.2 | 23.4 | 23 | 23.2 | 23.2 | 23.3 | 22.9 | 24.3 | 23.2 | 23 | 22.7 | 22.8 | 22.9 | 22.9 | 22.9 | 23.3 |
| Xanthobacter cornucopiae V4C-4 | 24 | 23.7 | 23.6 | 23.7 | 23.9 | 23.7 | 23.6 | 23.7 | 23.6 | 23.9 | 23.9 | 24.1 | 24 | 24.1 | 23.9 | 23.9 | 23.7 | 23.8 | 23.8 | 23.7 | 23.7 | 24.5 |
| Xanthobacter agilis MA37 | 23.9 | 23.7 | 23.5 | 23.7 | 23.7 | 23.6 | 23.7 | 23.9 | 23.6 | 23.7 | 23.7 | 23.4 | 23.3 | 23.4 | 23.3 | 23.4 | 23.3 | 23.4 | 23.4 | 23.4 | 23.4 | 23.9 |
| Xanthobacter agilis MA40 | 23.9 | 23.7 | 23.5 | 23.7 | 23.7 | 23.6 | 23.7 | 23.9 | 23.6 | 23.7 | 23.7 | 23.4 | 23.3 | 23.3 | 23.3 | 23.4 | 23.3 | 23.4 | 23.4 | 23.4 | 23.4 | 23.9 |
| Xanthobacter agilis SA35 | 23.8 | 23.6 | 23.3 | 23.6 | 23.5 | 23.7 | 24.3 | 23.7 | 23.4 | 23.5 | 23.5 | 23.8 | 23.3 | 24.4 | 23.8 | 23.5 | 23.4 | 23.3 | 23.5 | 23.5 | 23.5 | 23.8 |
| Xanthobacter albus V0C-6 | 24.1 | 24.1 | 23.9 | 24 | 24.2 | 23.9 | 24.1 | 24.1 | 23.9 | 24 | 24 | 23.8 | 23.8 | 23.9 | 23.8 | 23.9 | 23.7 | 23.8 | 23.9 | 23.9 | 23.9 | 24.4 |
| Xanthobacter albus V13C-5 | 24.1 | 24 | 23.8 | 24 | 24.1 | 23.9 | 24 | 24.1 | 23.9 | 23.9 | 23.9 | 23.7 | 23.7 | 23.8 | 23.7 | 23.8 | 23.7 | 23.7 | 23.8 | 23.8 | 23.8 | 24.3 |
| Xanthobacter albus V0B-10 | 24.2 | 24.1 | 23.9 | 24 | 24.2 | 23.9 | 24.1 | 24.1 | 23.9 | 24 | 24 | 23.8 | 23.8 | 23.9 | 23.8 | 23.9 | 23.7 | 23.8 | 23.9 | 23.9 | 23.9 | 24.4 |
| Xanthobacter albus V2C-4 | 24.1 | 23.9 | 23.8 | 23.9 | 23.8 | 23.8 | 24.1 | 24.7 | 23.9 | 24.3 | 24.2 | 24 | 23.9 | 23.7 | 23.6 | 23.7 | 23.7 | 23.9 | 23.9 | 23.9 | 23.9 | 24.6 |

**Supplementary Figure 6 – A comparison of average nucleotide identity values (ANI) calculated with FastANI between *X. autotrophicus* 7C$^T$ and *X. viscosus* 7d$^T$ genomes highlights extreme similarities between their genomes.** Included is the *X. autotrophicus* 7C$^T$ that we sequenced in this study, *X. autotrophicus* 7C$^T$ that was previously sequenced and uploaded to NCBI, *X. viscosus* 7d$^T$ that we sequenced in this study (sourced from DSMZ – DSM 21355$^T$), and *X. viscosus* 7d$^T$ that was previously sequenced and uploaded to JGI. All ANI values here are approximately 99.99% and above, highly indicating that *X. viscosus* 7d$^T$ should be reassigned to *X. autotrophicus*.

| | ***X. autotrophicus* 7C$^T$** (=DSM 432$^T$) (This Study - JBAFVK000000000) | ***X. autotrophicus* 7C$^T$** (=DSM 432$^T$) (NCBI – GCF_005871-85.1) | ***X. viscosus* 7d$^T$** (=DSM 21355$^T$) (This Study - JBAFVI000000000 ) | ***X. viscosus* 7d$^T$** (=DSM 21355$^T$) (JGI - Gp0538788 ) |
|---|---|---|---|---|
| ***X. autotrophicus* 7C$^T$** (=DSM 432$^T$) (This Study - JBAFVK000000000) | 100% | 99.9995% | 99.9977% | 99.9986% |
| ***X. autotrophicus* 7C$^T$** (=DSM 432$^T$) (NCBI – GCF_005871-85.1) | 99.9877% | 100% | 99.9893% | 99.9937% |
| ***X. viscosus* 7d$^T$** (=DSM 21355$^T$) (This Study - JBAFVI000000000 ) | 99.9955% | 99.9963% | 100% | 99.9992% |
| ***X. viscosus* 7d$^T$** (=DSM 21355$^T$) (JGI - Gp0538788 ) | 99.9905% | 99.9977% | 99.9955% | 100% |

**Supplementary Figure 7 – There are unique polymorphisms from the prior Doronina 2003 *X. viscosus* 7d$^T$ 16S rRNA gene sequence that are not found in any other *X. viscosus* 7d$^T$ 16S rRNA gene sequence or other Clade I Xanthobacter species—*X. autotrophicus*, *X. versatilis*, *X. oligotrophicus*, *X. toluenivorans*.** Prior studies have shown high similarity between *X. viscosus* and *X. autotrophicus* hence Clade I Xanthobacter species were used in the analysis. 16S rRNA genes were aligned using MAFFT v7.511 for the visualization. Positions that are not identical across the sequences have been highlighted: light blue for variants the same as *X. autotrophicus*, pink for variants completely unique to the *X. viscosus* 7d$^T$ sequence originally published in Doronina *et al.,* 2003 used to classify the strain as a new Xanthobacter species, and light green for all other variants. There are 10 different positions in the alignment where the Doronina *et al.,* 2003 16S rRNA gene sequence for *X. viscosus* 7d$^T$ differs across all other sequences (including 16S rRNA gene sequences from *X. viscosus* 7d$^T$ sequenced in this study and by another research group), raising concern.

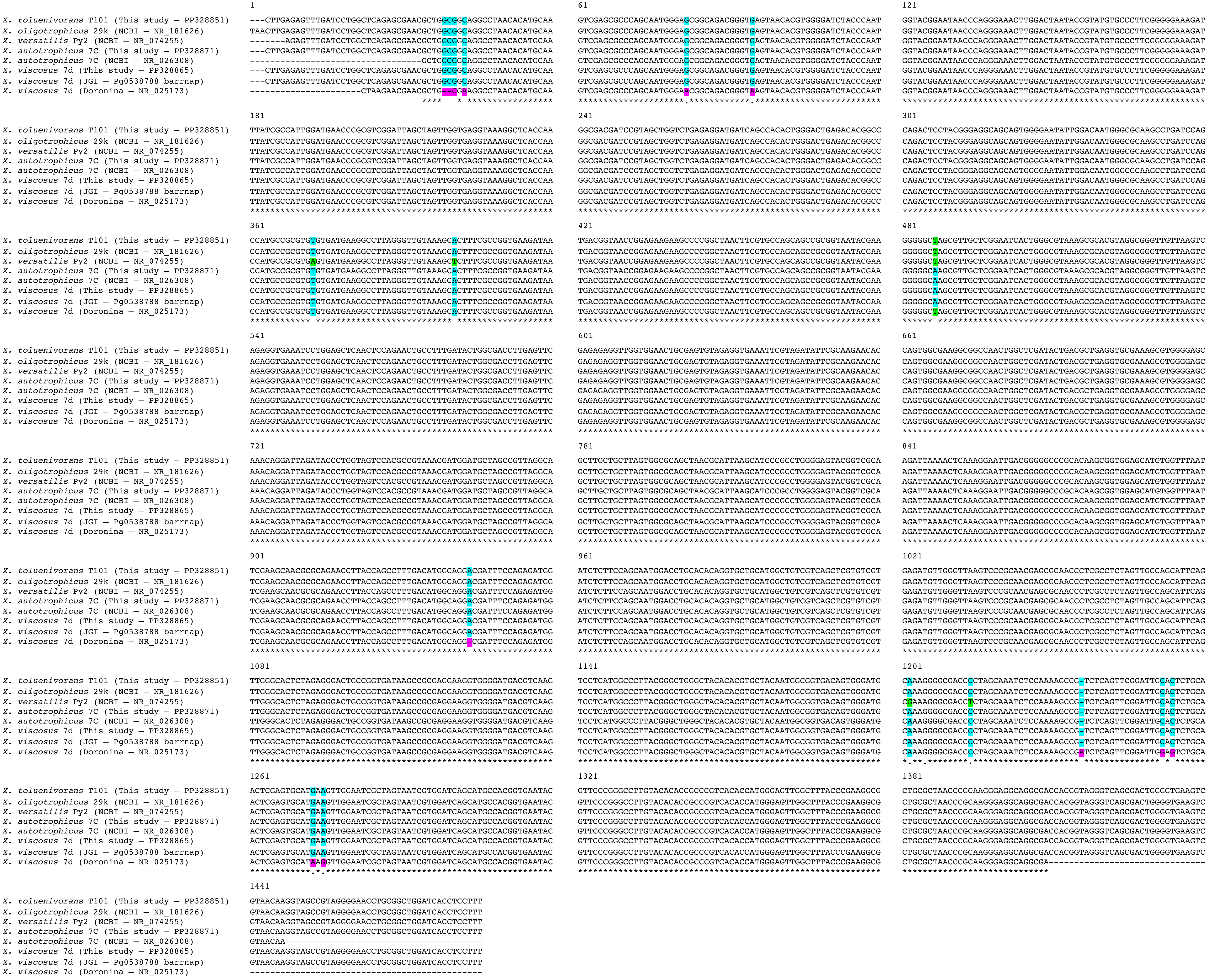

X. toluenivorans T101 (This study – PP328851)
X. oligotrophicus 29k (NCBI – NR_181626)
X. versatilis Py2 (NCBI – NR_074255)
X. autotrophicus 7C (This study – PP328871)
X. autotrophicus 7C (NCBI – NR_026308)
X. viscosus 7d (This study – PP328865)
X. viscosus 7d (JGI – Pg0538788 barrnap)
X. viscosus 7d (Doronina – NR_025173)
1
61
121
181
241
301
361
421
481
541
601
661
721
781
841
901
961
1021
1081
1141
1201
1261
1321
1381
1441

**Supplementary Figure 8 – The unique polymorphisms from the prior Doronina 2003 *X. viscosus* 7d$^T$ 16S rRNA gene sequence are not found in any other *X. viscosus* 7d$^T$ 16S rRNA gene sequence or in any other Xanthobacter species.** Prior studies have shown high similarity between *X. viscosus* and *X. autotrophicus* hence *X. viscosus* was included in Clade I. 16S rRNA genes were aligned using MAFFT v7.511 for the visualization. Positions that are not identical across the sequences have been highlighted: light blue for variants the same as *X. autotrophicus*, pink for variants completely unique to the *X. viscosus* 7d$^T$ sequence originally published in Doronina and Trotsenko 2003 used to classify the strain as a new Xanthobacter species, and light green and yellow for all other variants. There are 10 different positions in the alignment where the Doronina and Trotsenko 2003 16S rRNA gene sequence for *X. viscosus* 7d$^T$ differs across all other Xanthobacter sequences (including 16S rRNA gene sequences from *X. viscosus* 7d$^T$ sequenced in this study and by another research group), raising concern.

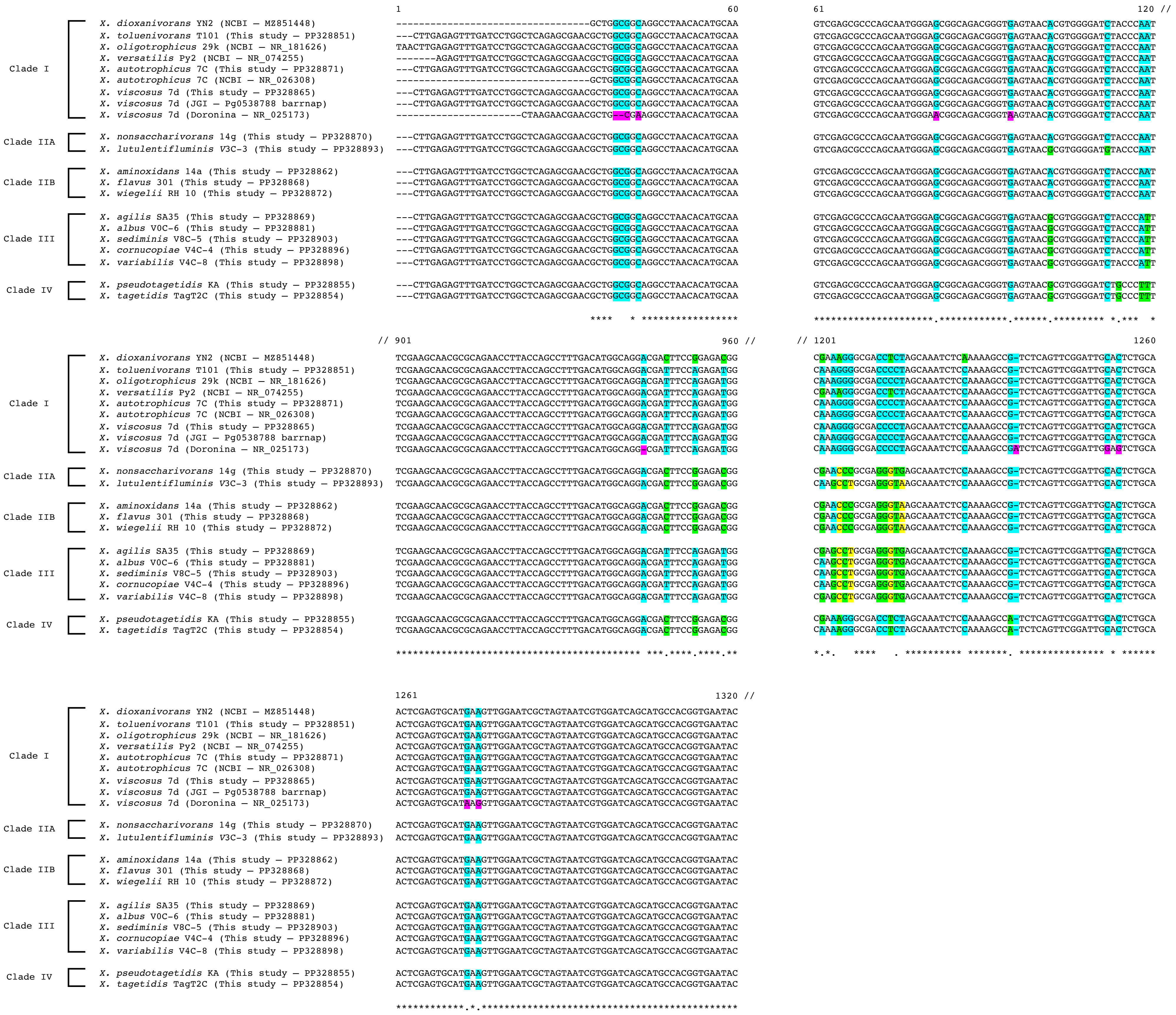
1
60
61
120 //
// 901
960 //
// 1201
1260
1261
1320 //
Clade I
X. dioxanivorans YN2 (NCBI – MZ851448)
X. toluenivorans T101 (This study – PP328851)
X. oligotrophicus 29k (NCBI – NR_181626)
X. versatilis Py2 (NCBI – NR_074255)
X. autotrophicus 7C (This study – PP328871)
X. autotrophicus 7C (NCBI – NR_026308)
X. viscosus 7d (This study – PP328865)
X. viscosus 7d (JGI – Pg0538788 barrnap)
X. viscosus 7d (Doronina – NR_025173)
Clade IIA
X. nonsaccharivorans 14g (This study – PP328870)
X. lutulentifluminis V3C-3 (This study – PP328893)
Clade IIB
X. aminoxidans 14a (This study – PP328862)
X. flavus 301 (This study – PP328868)
X. wiegelii RH 10 (This study – PP328872)
Clade III
X. agilis SA35 (This study – PP328869)
X. albus V0C-6 (This study – PP328881)
X. sediminis V8C-5 (This study – PP328903)
X. cornucopiae V4C-4 (This study – PP328896)
X. variabilis V4C-8 (This study – PP328898)
Clade IV
X. pseudotagetidis KA (This study – PP328855)
X. tagetidis TagT2C (This study – PP328854)

**Supplementary Figure 9 – Type and representative strains of Xanthobacter and Roseixanthobacter gen. nov. when grown on rich media (2xM1) or minimal media (4M) containing succinate (0.5% w/v) demonstrate slime production capabilities that largely follow along evolutionary lines.** Growth on succinate is a known slime-inducing condition for some Xanthobacter.

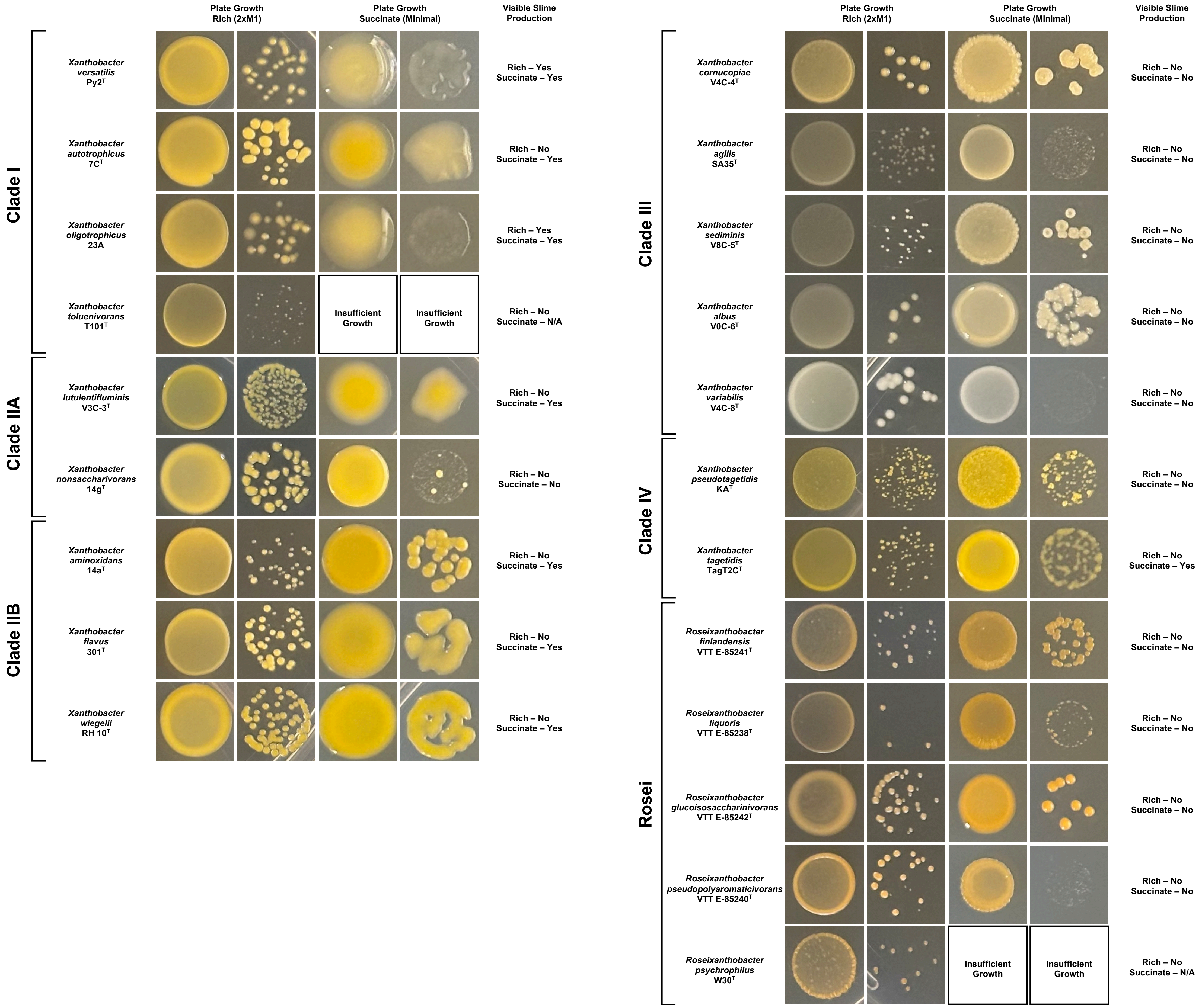

Plate Growth Rich (2xM1)
Plate Growth Succinate (Minimal)
Visible Slime Production
Clade I
Xanthobacter versatilis Py2T
Rich – Yes Succinate – Yes
Xanthobacter autotrophicus 7CT
Rich – No Succinate – Yes
Xanthobacter oligotrophicus 23A
Rich – Yes Succinate – Yes
Xanthobacter toluenivorans T101T
Insufficient Growth
Insufficient Growth
Rich – No Succinate – N/A
Clade IIA
Xanthobacter lutulentifluminis V3C-3T
Rich – No Succinate – Yes
Xanthobacter nonsaccharivorans 14gT
Rich – No Succinate – No
Clade IIB
Xanthobacter aminoxidans 14aT
Rich – No Succinate – Yes
Xanthobacter flavus 301T
Rich – No Succinate – Yes
Xanthobacter wiegelii RH 10T
Rich – No Succinate – Yes
Plate Growth Rich (2xM1)
Plate Growth Succinate (Minimal)
Visible Slime Production
Clade III
Xanthobacter cornucopiae V4C-4T
Rich – No Succinate – No
Xanthobacter agilis SA35T
Rich – No Succinate – No
Xanthobacter sediminis V8C-5T
Rich – No Succinate – No
Xanthobacter albus V0C-6T
Rich – No Succinate – No
Xanthobacter variabilis V4C-8T
Rich – No Succinate – No
Clade IV
Xanthobacter pseudotagetidis KAT
Rich – No Succinate – No
Xanthobacter tagetidis TagT2CT
Rich – No Succinate – Yes
Rosei
Roseixanthobacter finlandensis VTT E-85241T
Rich – No Succinate – No
Roseixanthobacter liquoris VTT E-85238T
Rich – No Succinate – No
Roseixanthobacter glucoisosaccharinivorans VTT E-85242T
Rich – No Succinate – No
Roseixanthobacter pseudopolyaromaticivorans VTT E-85240T
Rich – No Succinate – No
Roseixanthobacter psychrophilus W30T
Insufficient Growth
Insufficient Growth
Rich – No Succinate – N/A

**Supplementary Figure 10 – Cell morphologies of Xanthobacter and Roseixanthobacter gen. nov. species may become branched when grown in in minimal media (4M) containing succinate (0.5% w/v) in comparison to rich media (2xM1) in some evolutionarily related clades.** Growth on succinate is a known morphology-changing condition for some Xanthobacter. Imaging was performed on a Nikon Eclipse TI microscope modified for QPI (with a Phasics SID4BIO) as described in Liu *et al.,* 2020. Images were acquired using the Nikon Plan Apo 100X Phase objective with the 1.5X magnifier with a halogen lamp as the light source.

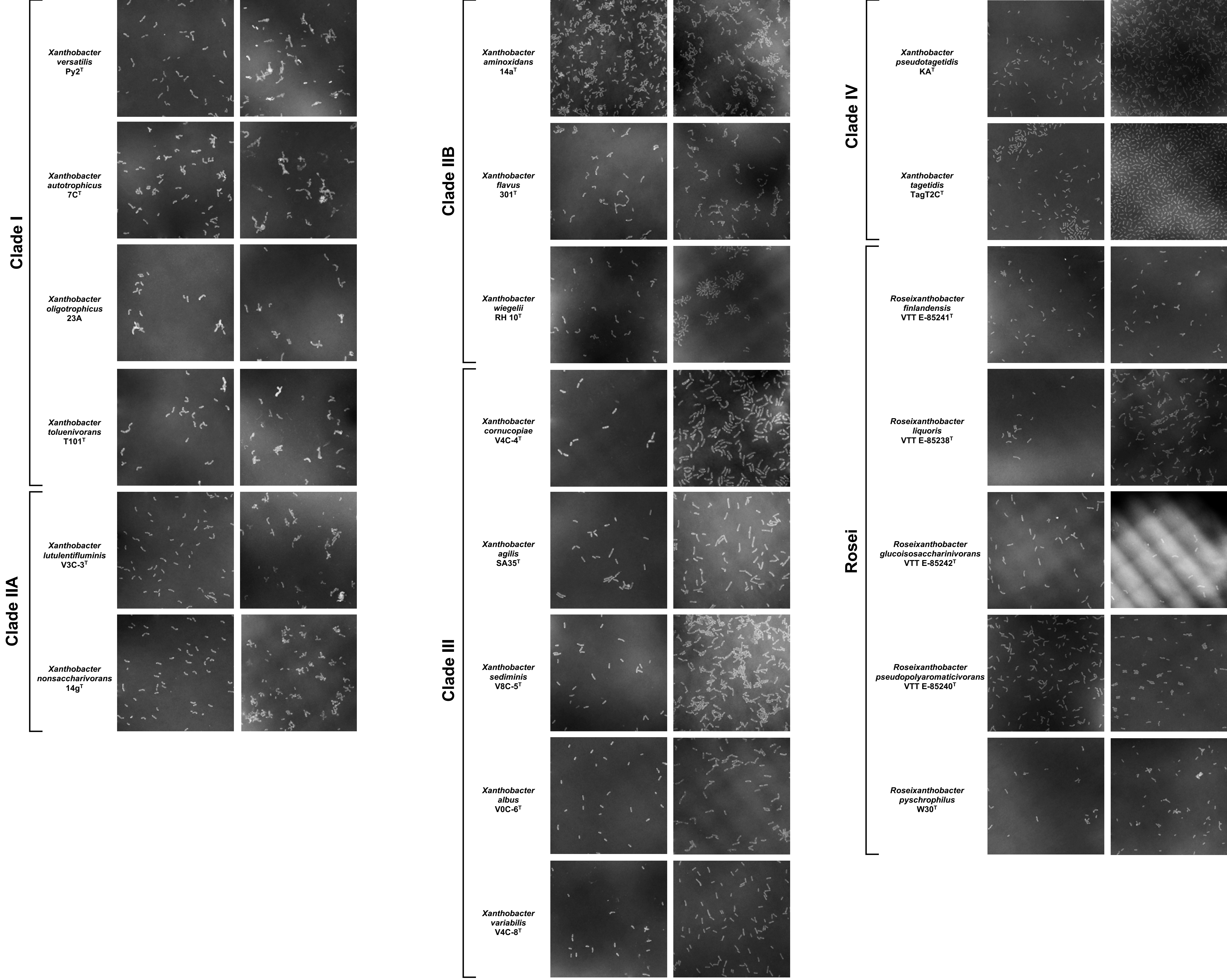
Clade I
Xanthobacter versatilis Py2$^{T}$
Xanthobacter autotrophicus 7C$^{T}$
Xanthobacter oligotrophicus 23A
Xanthobacter toluenivorans T101$^{T}$
Clade IIA
Xanthobacter lutulentifluminis V3C-3$^{T}$
Xanthobacter nonsaccharivorans 14g$^{T}$
Clade IIB
Xanthobacter aminoxidans 14a$^{T}$
Xanthobacter flavus 301$^{T}$
Xanthobacter wiegelii RH 10$^{T}$
Clade III
Xanthobacter cornucopiae V4C-4$^{T}$
Xanthobacter agilis SA35$^{T}$
Xanthobacter sediminis V8C-5$^{T}$
Xanthobacter albus V0C-6$^{T}$
Xanthobacter variabilis V4C-8$^{T}$
Clade IV
Xanthobacter pseudotagetidis KA$^{T}$
Xanthobacter tagetidis TagT2C$^{T}$
Rosei
Roseixanthobacter finlandensis VTT E-85241$^{T}$
Roseixanthobacter liquoris VTT E-85238$^{T}$
Roseixanthobacter glucoisosaccharinivorans VTT E-85242$^{T}$
Roseixanthobacter pseudopolyaromaticivorans VTT E-85240$^{T}$
Roseixanthobacter pyschrophilus W30$^{T}$

**Supplementary Table 5 – Cell size dimensions of rod-shaped cells of the type and representative strains of Xanthobacter and Roseixanthobacter gen. nov. grown in rich media (2xM1).** Imaging was performed on a Nikon Eclipse TI microscope modified for QPI (with a Phasics SID4BIO) as described in Liu *et al.,* 2020. Images were acquired using the Nikon Plan Apo 100X Phase objective with the 1.5X magnifier with a halogen lamp as the light source.

| **Species** | **Clade** | **Median Length μm (± Standard Deviation)** | **Median Width μm (± Standard Deviation)** |
|---|---|---|---|
| *X. versatilis* Py2$^T$ | I | 1.29 (± 0.38) | 0.47 (± 0.09) |
| *X. autotrophicus* 7C$^T$ | I | 1.23 (± 0.34) | 0.49 (±0.10) |
| *X. oligotrophicus* 23A | I | 1.36 (± 0.56) | 0.49 (± 0.17) |
| *X. toluenivorans* T101$^T$ | I | 1.43 (± 0.50) | 0.51 (± 0.19) |
| *X. lutulentifluminis* V3C-3$^T$ | IIA | 1.30 (± 0.36) | 0.41 (± 0.08) |
| *X. nonsaccharivorans* 14g$^T$ | IIA | 1.42 (± 0.36) | 0.48 (± 0.09) |
| *X. aminoxidans* 14a$^T$ | IIB | 1.12 (± 0.29) | 0.41 (± 0.09) |
| *X. flavus* 301$^T$ | IIB | 1.48 (± 0.42) | 0.45 (± 0.11) |
| *X. wiegelii* RH 10$^T$ | IIB | 1.38 (± 0.41) | 0.43 (± 0.08) |
| *X. cornucopiae* V4C-4$^T$ | III | 1.64 (± 0.54) | 0.48 (± 0.18) |
| *X. agilis* SA35$^T$ | III | 1.63 (± 0.32) | 0.48 (± 0.06) |
| *X. sediminis* V8C-5$^T$ | III | 1.54 (± 0.38) | 0.46 (± 0.08) |
| *X. albus* V0C-6$^T$ | III | 1.30 (± 0.34) | 0.49 (± 0.09) |
| *X. variabilis* V4C-8$^T$ | III | 1.35 (± 0.33) | 0.44 (± 0.09) |
| *X. pseudotagetidis* KA$^T$ | IV | 1.22 (± 0.32) | 0.36 (± 0.09) |
| *X. tagetidis* TagT2C$^T$ | IV | 1.12 (± 0.33) | 0.38 (0.09) |
| *R. finlandensis* VTT E-85241$^T$ | Rosei | 1.25 (± 0.48) | 0.39 (± 0.13) |
| *R. liquoris* VTT E-85238$^T$ | Rosei | 1.40 (± 0.60) | 0.41 (± 0.13) |
| *R. glucoisosaccharinivorans* VTT E-85242$^T$ | Rosei | 1.71 (± 0.58) | 0.38 (± 0.10) |
| *R. pseudopolyaromaticivorans* VTT E-85240$^T$ | Rosei | 1.27 (± 0.41) | 0.36 (± 0.07) |
| *R. psychrophilus* W30$^T$ | Rosei | 1.30 (± 0.38) | 0.42 (± 0.09) |

**Supplementary Figure 11 – Evolutionarily related Xanthobacter and Roseixanthobacter gen. nov. species have similar median rod-shaped cell size dimensions when grown in rich media (2xM1).** Imaging was performed on a Nikon Eclipse TI microscope modified for QPI (with a Phasics SID4BIO) as described in Liu *et al.,* 2020. Images were acquired using the Nikon Plan Apo 100X Phase objective with the 1.5X magnifier with a halogen lamp as the light source. Abbreviations are: XVE – *Xanthobacter versatilis* Py2$^T$; XAU – *Xanthobacter autotrophicus* 7C$^T$; XOG – *Xanthobacter oligotrophicus* 23A; XTO – *Xanthobacter toluenivorans* T101$^T$; XLU – *Xanthobacter lutulentifluminis* V3C-3$^T$; XNS – *Xanthobacter nonsaccharivorans* 14g$^T$; XAM – Xanthobacter aminoxidans 14a$^T$; XFL – Xanthobacter flavus 301$^T$; XWE – Xanthobacter wiegelii RH 10$^T$; XPS – *Xanthobacter pseudotagetidis* KA$^T$; XTA – *Xanthobacter tagetidis* TagT2C$^T$; XCO – *Xanthobacter cornucopiae* V4C-4$^T$; XAG – *Xanthobacter agilis* SA35$^T$; XSE – *Xanthobacter sediminis* V8C-5$^T$; XAL – *Xanthobacter albus* V0C-6$^T$; XVA – *Xanthobacter variabilis* V4C-8$^T$; RFI – *Roseixanthobacter finlandensis* VTT E-85241$^T$; RLI – *Roseixanthobacter liquoris* VTT E-85238$^T$; RGL – *Roseixanthobacter glucoisosaccharinivorans* VTT E-85242$^T$; RPP – *Roseixanthobacter pseudopolyaromaticivorans* VTT E-85240$^T$; RSY – *Roseixanthobacter psychrophilus* W30$^T$.

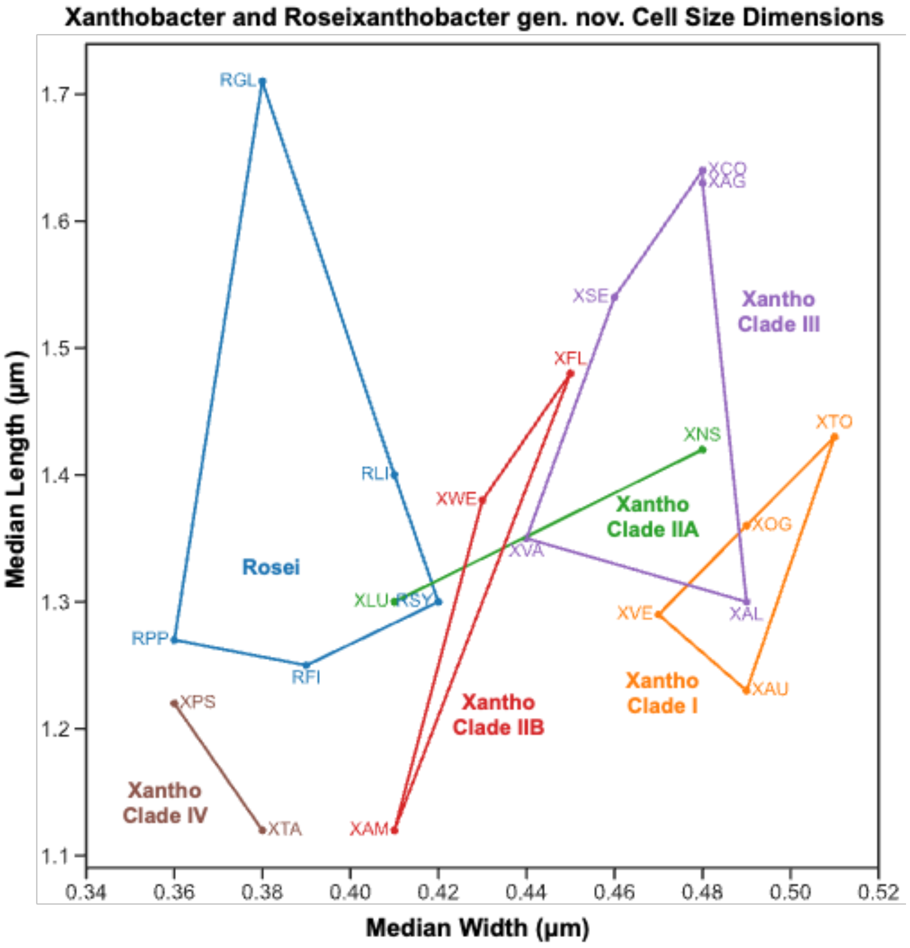

**Supplementary Table 6 – Bioinformatic analysis of flagella protein genes (*fli, flg, flh, motA/motB*) across Xanthobacter species type strains indicates all Xanthobacter have some non-operonic flagella proteins, but only Xanthobacter with an operonic flagella region are motile in semisolid agar.** Flagella proteins were identified using annotations obtained from the genomes using NCBI's PGAP software. Some Xanthobacter genomes (including *X. agilis* and *X. flavus* which have been observed to have long peritrichous flagella) contain an approximately 40 kb region in the genome with dozens of flagella protein genes arranged in operons. All Xanthobacter species (including those previously described as non-motile and *X. tagetidis* and *X. dioxanivorans* which have been observed to have short peritrichous protrusions unlike those flagella seen in *X. agilis* and *X. flavus*) have non-operonic flagella proteins scattered throughout the genome, often standalone. PBP is an abbreviation for a gene annotation described as a peptidoglycan-binding protein (similar to the function of motB). Motility as measured with the semisolid agar assay (Supplementary Table 7) in this study is indicated in comparison to motility observed and described in the literature.

| Species | Clade | Non-Operonic Flagella Proteins | | | | Xanthobacter Operonic Flagella Region (*fli, flg, flh, motA/motB*) ~40 kb | Semisolid Agar Motility (This Study) | Motility (Past Literature) | |
|---|---|---|---|---|---|---|---|---|---|
| | | *flgA* | *fliO* | *fliJ* | motA –PBP ("motB") | | | Motility | Method |
| *X. versatilis* Py2$^T$ | I | X | X | X | | | | No (van Ginkel 1986) | Microscopy (?) |
| *X. autotrophicus* 7C$^T$ | I | | X | X | X | | | No (Baumgarten 1974) | Microscopy (?) |
| *X. oligotrophicus* 29k$^T$ | I | | X | X | X | | *N/A* | No (Tikhonova 2021) | Microscopy |
| *X. toluenivorans* T101$^T$ | I | | X | X | X | | | No (Tay 1999) | Microscopy (?) |
| *X. dioxanivorans* YN2$^T$ | I | X | X | X | X | | *N/A* | Yes (Wang 2021) | Phase Contrast Microscopy |
| *X. lutulentifluminis* V3C-3$^T$ | IIA | | X | X | X | | | - | - |
| *X. nonsaccharivorans* 14g$^T$ | IIA | | X | X | X | | | No (Baumgarten 1974) | Microscopy (?) |
| *X. aminoxidans* 14a$^T$ | IIB | X | X | X | X | X | X (Inducible) | No (Doronina 1984, Doronina 2003) | Unclear (Original Publication Inaccessible) |
| *X. flavus* 301$^T$ | IIB | | X | X | X | X | X (Inducible) | Yes (Reding 1992) | Wet Mount - Light Microscopy |
| *X. wiegelii* RH 10$^T$ | IIB | | X | X | X | X | X (Inducible) | No (Baumgarten 1974) | Microscopy (?) |
| *X. cornucopiae* V4C-4$^T$ | III | | X | X | X | X | X (Constitutive) | - | - |
| *X. agilis* SA35$^T$ | III | | X | X | X | X | X (Constitutive) | Yes (Jenni 1987, Reding 1992) | Wet Mount – Light Microscopy |
| *X. sediminis* V8C-5$^T$ | III | | X | X | X | X | X (Constitutive) | - | - |
| *X. albus* V0C-6$^T$ | III | | X | X | X | X | X (Constitutive) | - | - |
| *X. variabilis* V4C-8$^T$ | III | | X | X | X | X | X (Constitutive) | - | - |
| *X. pseudotagetidis* KA$^T$ | IV | | | X | X | | | - | - |
| *X. tagetidis* TagT2C$^T$ | IV | | X | X | X | | | Yes (Padden 1997) | Light Microscopy |

**Supplementary Table 7 – Motility within semisolid agar follows along evolutionarily related clades for Xanthobacter and Roseixanthobacter gen. nov. type strains**. Strains were grown in minimal media (4M), supplemented with vitamins, triphenyltetrazolium chloride for visualization, 0.3% agar, and 0.3% w/v or v/v of the appropriate carbon source. *Xanthobacter oligotrophicus* (XOG) is represented here by strain 23A instead of type strain 29k$^T$. A positive result is indicated with a "+" where the entire assay tube was turbid. A weakly positive result is indicated with a "w", where larger regions around the inoculation line was turbid. If no growth of the strain was observed on the substrate it is indicated with "ng" on the table. Fully positive results were apparent by 5 days, with all observations below taken after 2 weeks of incubation at 30°C. Abbreviations are: XVE – *Xanthobacter versatilis* Py2$^T$; XAU – *Xanthobacter autotrophicus* 7C$^T$; XOG – *Xanthobacter oligotrophicus* 23A; XTO – *Xanthobacter toluenivorans* T101$^T$; XLU – *Xanthobacter lutulentifluminis* V3C-3$^T$; XNS – *Xanthobacter nonsaccharivorans* 14g$^T$; XAM – Xanthobacter aminoxidans 14a$^T$; XFL – Xanthobacter flavus 301$^T$; XWE – Xanthobacter wiegelii RH 10$^T$; XPS – *Xanthobacter pseudotagetidis* KA$^T$; XTA – *Xanthobacter tagetidis* TagT2C$^T$; XCO – *Xanthobacter cornucopiae* V4C-4$^T$; XAG – *Xanthobacter agilis* SA35$^T$; XSE – *Xanthobacter sediminis* V8C-5$^T$; XAL – *Xanthobacter albus* V0C-6$^T$; XVA – *Xanthobacter variabilis* V4C-8$^T$; RFI – *Roseixanthobacter finlandensis* VTT E-85241$^T$; RLI – *Roseixanthobacter liquoris* VTT E-85238$^T$; RGL – *Roseixanthobacter glucoisosaccharinivorans* VTT E-85242$^T$; RPP – *Roseixanthobacter pseudopolyaromaticivorans* VTT E-85240$^T$; RSY – *Roseixanthobacter psychrophilus* W30$^T$. XCO, XAG, XSE, XAL, XVA exhibited extremely poor growth on gluconate but nonetheless still were positive for motility with the assay.

| Condition | Species | | | | | | | | | | | | | | | | | | | | |
|---|---|---|---|---|---|---|---|---|---|---|---|---|---|---|---|---|---|---|---|---|---|
| | Clade I | | | | Clade IIA | | Clade IIB | | | Clade III | | | | | Clade IV | | Roseixanthobacter | | | | |
| | XVE | XAU | XOG* | XTO | XLU | XNS | XAM | XFL | XWE | XCO | XAG | XSE | XAL | XVA | XPS | XTA | RFI | RLI | RGL | RPP | RSY |
| Flagella motility pathway | | | | | | | + | + | + | + | + | + | + | + | | | + | + | + | + | |
| Succinate motility | | | | | | | | | | + | + | + | + | + | | | + | | | | |
| Fumarate motility | | | | ng | | | w | w | w | + | + | + | + | + | | | + | | w | w | |
| Gluconate motility | | | | | | | w | w | w | + | + | + | + | + | | | + | w | + | w | ng |
| Methanol motility | | | | | | | w | | w | + | + | + | + | + | ng | ng | w | ng | ng | ng | ng |
| Ethanol motility | | | | | | | w | w | w | + | + | + | + | + | ng | ng | + | ng | ng | ng | ng |
| Propanol motility | | | | | | | w | w | w | + | + | + | + | + | ng | ng | w | ng | ng | ng | ng |
| Isopropanol motility | | | | | | | | | w | + | + | + | + | + | | ng | w | ng | ng | ng | |

**Supplementary Table 8 – Xanthobacter and Roseixanthobacter gen. nov. type strains are mesophilic as observed during temperature testing.** Strains were grown for 7 days on rich media (2xM1) plates at the given temperature. Positive growth observed is indicated with a "+" while maximum growth conditions are indicated with "Max". *Xanthobacter oligotrophicus* (XOG) is represented here by strain 23A instead of type strain 29k$^T$. Abbreviations are: XVE – *Xanthobacter versatilis* Py2$^T$; XAU – *Xanthobacter autotrophicus* 7C$^T$; XOG – *Xanthobacter oligotrophicus* 23A; XTO – *Xanthobacter toluenivorans* T101$^T$; XLU – *Xanthobacter lutulentifluminis* V3C-3$^T$; XNS – *Xanthobacter nonsaccharivorans* 14g$^T$; XAM – Xanthobacter aminoxidans 14a$^T$; XFL – Xanthobacter flavus 301$^T$; XWE – Xanthobacter wiegelii RH 10$^T$; XPS – *Xanthobacter pseudotagetidis* KA$^T$; XTA – *Xanthobacter tagetidis* TagT2C$^T$; XCO – *Xanthobacter cornucopiae* V4C-4$^T$; XAG – *Xanthobacter agilis* SA35$^T$; XSE – *Xanthobacter sediminis* V8C-5$^T$; XAL – *Xanthobacter albus* V0C-6$^T$; XVA – *Xanthobacter variabilis* V4C-8$^T$; RFI – *Roseixanthobacter finlandensis* VTT E-85241$^T$; RLI – *Roseixanthobacter liquoris* VTT E-85238$^T$; RGL – *Roseixanthobacter glucoisosaccharinivorans* VTT E-85242$^T$; RPP – *Roseixanthobacter pseudopolyaromaticivorans* VTT E-85240$^T$; RSY – *Roseixanthobacter psychrophilus* W30$^T$.

| Temperature (°C) | Species | | | | | | | | | | | | | | | | | | | | |
|---|---|---|---|---|---|---|---|---|---|---|---|---|---|---|---|---|---|---|---|---|---|
| | Clade I | | | | Clade IIA | | Clade IIB | | | Clade III | | | | | Clade IV | | Roseixanthobacter | | | | |
| | XVE | XAU | XOG* | XTO | XLU | XNS | XAM | XFL | XWE | XCO | XAG | XSE | XAL | XVA | XPS | XTA | RFI | RLI | RGL | RPP | RSY |
| 4 | | | | | | | | | | | | | | | | | | | | | |
| 10 | + | + | + | + | | | + | + | + | + | + | | | | | + | + | + | + | + | + |
| 15 | + | + | + | + | + | + | + | + | + | + | + | + | + | + | + | + | + | + | + | + | + |
| 20 | + | + | + | + | + | + | + | + | + | + | + | + | + | + | + | + | + | + | + | + | + |
| 25 | + | + | + | + | + | + | + | + | + | + | + | + | + | + | + | + | Max | Max | Max | Max | Max |
| 30 | Max | Max | Max | Max | Max | Max | Max | Max | Max | Max | Max | Max | Max | Max | Max | Max | Max | Max | Max | Max | Max |
| 37 | Max | Max | Max | | Max | Max | Max | Max | Max | Max | Max | Max | Max | Max | Max | Max | + | + | + | + | |
| 42 | | | | | | | + | + | | | | | | + | + | | | | | | |

**Supplementary Table 9 – Antibiotic susceptibility is species specific for Xanthobacter and Roseixanthobacter gen. nov. type strains.** Antibiotic susceptibility was tested on rich media (2xM1) plates at the given concentrations. A 10-fold dilution series starting from a culture of $OD_{600}$ 0.1 was spot plated onto the antibiotic plates. Gen – gentamicin; Kan – kanamycin; Str – streptomycin; Tet – tetracycline. Resistance is indicated with an "Res" where all dilution spots for the species grew on the antibiotic at that concentration. Breakthrough (one or two colonies or very light film growth in some or one dilution spot) is indicated with a "Brk". *Xanthobacter oligotrophicus* (XOG) is represented here by strain 23A instead of type strain 29k$^T$. Abbreviations are: XVE – *Xanthobacter versatilis* Py2$^T$; XAU – *Xanthobacter autotrophicus* 7C$^T$; XOG – *Xanthobacter oligotrophicus* 23A; XTO – *Xanthobacter toluenivorans* T101$^T$; XLU – *Xanthobacter lutulentifluminis* V3C-3$^T$; XNS – *Xanthobacter nonsaccharivorans* 14g$^T$; XAM – Xanthobacter aminoxidans 14a$^T$; XFL – Xanthobacter flavus 301$^T$; XWE – Xanthobacter wiegelii RH 10$^T$; XPS – *Xanthobacter pseudotagetidis* KA$^T$; XTA – *Xanthobacter tagetidis* TagT2C$^T$; XCO – *Xanthobacter cornucopiae* V4C-4$^T$; XAG – *Xanthobacter agilis* SA35$^T$; XSE – *Xanthobacter sediminis* V8C-5$^T$; XAL – *Xanthobacter albus* V0C-6$^T$; XVA – *Xanthobacter variabilis* V4C-8$^T$; RFI – *Roseixanthobacter finlandensis* VTT E-85241$^T$; RLI – *Roseixanthobacter liquoris* VTT E-85238$^T$; RGL – *Roseixanthobacter glucoisosaccharinivorans* VTT E-85242$^T$; RPP – *Roseixanthobacter pseudopolyaromaticivorans* VTT E-85240$^T$; RSY – *Roseixanthobacter psychrophilus* W30$^T$.

| Antibiotic (ug/mL) | | Species | | | | | | | | | | | | | | | | | | | | |
|---|---|---|---|---|---|---|---|---|---|---|---|---|---|---|---|---|---|---|---|---|---|---|
| | | Clade I | | | | Clade IIA | | Clade IIB | | | Clade III | | | | | Clade IV | | Roseixanthobacter | | | | |
| | | XVE | XAU | XOG* | XTO | XLU | XNS | XAM | XFL | XWE | XCO | XAG | XSE | XAL | XVA | XPS | XTA | RFI | RLI | RGL | RPP | RSY |
| Gen | 20 | | | | | Brk | | | | | | | | | | | Res | | | | | |
| | 40 | | | | | | | | | | | | | | | | Res | | | | | |
| | 60 | | | | | | | | | | | | | | | | | | | | | |
| | 80 | | | | | | | | | | | | | | | | | | | | | |
| | 100 | | | | | | | | | | | | | | | | | | | | | |
| Kan | 12.5 | | | | | | | | | | | | | | | | Res | | | | | |
| | 25 | | | | | | | | | | | | | | | | Res | | | | | |
| | 50 | | | | | | | | | | | | | | | | Brk | | | | | |
| | 75 | | | | | | | | | | | | | | | | | | | | | |
| | 100 | | | | | | | | | | | | | | | | | | | | | |
| Str | 25 | | | | | | Brk | Brk | | Brk | | | | | | | | | | Brk | | |
| | 50 | | | | | | Brk | | | Brk | | | | | Brk | | | | | Brk | | |
| | 75 | | | | | | Brk | | | Brk | | | | | Brk | | | | | Brk | | |
| | 100 | | | | | | | | | | | | | | | | | | | | | |
| | 150 | | | | | | | | | | | | | | | | | | | | | |
| Tet | 3 | | Brk | | | Brk | Brk | | | | | | | | Res | | | | | | | |
| | 6 | | | | | Brk | | | | | | | | | | | | | | | | |
| | 9 | | | | | Brk | | | | | | | | | | | | | | | | |
| | 12 | | | | | | | | | | | | | | | | | | | | | |
| | 15 | | | | | | | | | | | | | | | | | | | | | |

**Supplementary Table 10 – Xanthobacter and Roseixanthobacter gen. nov. type strains are neutrophilic to slightly acidophilic in pH testing.** Numbers indicate the number of doublings from inoculation density ($OD_{600}$ 0.01) that have occurred based on optical density measurements. Results above 1 doubling are considered a positive in Table 3 of the main text and are underlined in the table. Results below 1 doubling are considered a negative result and are not reflected in the ranges in Table 3 in the main text. pH testing was run twice and the doubling values in the table are averages of the two repeats for pH values 3 to 7. pH values for 8 to 10 were only done once as the initial buffer used in this range for the first experiment was incompatible with Xanthobacter growth and were changed after the first experiment. *Xanthobacter oligotrophicus* (XOG) is represented here by strain 23A instead of type strain 29k$^T$. Abbreviations are: XVE – *Xanthobacter versatilis* Py2$^T$; XAU – *Xanthobacter autotrophicus* 7C$^T$; XOG – *Xanthobacter oligotrophicus* 23A; XTO – *Xanthobacter toluenivorans* T101$^T$; XLU – *Xanthobacter lutulentifluminis* V3C-3$^T$; XNS – *Xanthobacter nonsaccharivorans* 14g$^T$; XAM – Xanthobacter aminoxidans 14a$^T$; XFL – Xanthobacter flavus 301$^T$; XWE – Xanthobacter wiegelii RH 10$^T$; XPS – *Xanthobacter pseudotagetidis* KA$^T$; XTA – *Xanthobacter tagetidis* TagT2C$^T$; XCO – *Xanthobacter cornucopiae* V4C-4$^T$; XAG – *Xanthobacter agilis* SA35$^T$; XSE – *Xanthobacter sediminis* V8C-5$^T$; XAL – *Xanthobacter albus* V0C-6$^T$; XVA – *Xanthobacter variabilis* V4C-8$^T$; RFI – *Roseixanthobacter finlandensis* VTT E-85241$^T$; RLI – *Roseixanthobacter liquoris* VTT E-85238$^T$; RGL – *Roseixanthobacter glucoisosaccharinivorans* VTT E-85242$^T$; RPP – *Roseixanthobacter pseudopolyaromaticivorans* VTT E-85240$^T$; RSY – *Roseixanthobacter psychrophilus* W30$^T$.

| Condition | | Clade I | | | | Clade IIA | | Clade IIB | | | Clade III | | | | | Clade IV | | Rosei | | | | |
|---|---|---|---|---|---|---|---|---|---|---|---|---|---|---|---|---|---|---|---|---|---|---|
| | | XVE | XAU | XOG | XTO | XLU | XNS | XAM | XFL | XWE | XCO | XAG | XSE | XAL | XVA | XPS | XTA | RFI | RLI | RGL | RPP | RSY |
| Citric | pH 3 | 0 | 0.8 | 0.2 | 0.6 | 0.4 | 0.8 | 0.5 | 0.4 | 0 | 0.9 | 0.7 | 1.1 | 1 | 0.4 | 0.8 | 0 | 0.7 | 1.1 | 1.3 | 0.8 | 1 |
| | pH 4 | 0.3 | 1.2 | 0.5 | 0.8 | 0.5 | 0.5 | 0.7 | 0.9 | 0.9 | 0.8 | 0.6 | 0.7 | 0.8 | 1.5 | 0.5 | 0.6 | 0.4 | 0.9 | 0.1 | 0.6 | 0.7 |
| | pH 5 | 1.4 | 2 | 2.3 | 1.1 | 1.6 | 1.5 | 7.9 | 5.1 | 1.2 | 1.9 | 1.1 | 2.2 | 2.7 | 3.9 | 1.4 | 1.2 | 6.8 | 4.2 | 6.8 | 3.3 | 6.6 |
| Phosphate | pH 6 | 7.2 | 6.5 | 1.9 | 1.5 | 6.8 | 6.7 | 7 | 6.5 | 6.7 | 4.1 | 4.6 | 4.6 | 4.8 | 5.1 | 6 | 4.8 | 5.9 | 3.3 | 6.5 | 4.4 | 6.1 |
| | pH 7 | 7 | 5.4 | 2.9 | 0.1 | 5.4 | 6.8 | 7 | 6.7 | 6.3 | 4.9 | 3.9 | 4.6 | 5.1 | 5.5 | 6.7 | 6.6 | 6.1 | 3.4 | 6.5 | 1.2 | 0 |
| | pH 8 | 6.3 | 6.3 | 5.5 | 0 | 5.9 | 5.9 | 6.4 | 5.8 | 5.9 | 5.7 | 2.5 | 4.9 | 4.7 | 5.8 | 6.8 | 6.6 | 2.2 | 0 | 1.1 | 0 | 1.1 |
| Carbonate | pH 9 | 0 | 0 | 0 | 0 | 0 | 0 | 0 | 0 | 0 | 0 | 0 | 0 | 0 | 0 | 0 | 0 | 0 | 0 | 0 | 0 | 0 |
| | pH 10 | 0 | 0 | 0 | 0.8 | 0 | 0 | 0 | 0 | 0 | 0 | 0 | 0 | 0 | 0 | 0 | 0 | 0 | 0 | 0 | 0 | 0 |

**Supplementary Table 11 – Xanthobacter and Roseixanthobacter gen. nov. type strains largely have low salinity preferences in salinity testing.** Numbers indicate the number of doublings from inoculation density ($OD_{600}$ 0.01) that have occurred based on optical density measurements. Results above 1 doubling are considered a positive in Table 3 of the main text and are underlined in the table. Results below 1 doubling are considered a negative result and are not reflected in the ranges in Table 3 in the main text. Salinity testing was run twice and the doubling values in the table are averages of the two repeats. Note that the majority of the Roseixanthobacter strains flocculate which may affect OD readout. *Xanthobacter oligotrophicus* (XOG) is represented here by strain 23A instead of type strain 29k$^T$. Abbreviations are: XVE – *Xanthobacter versatilis* Py2$^T$; XAU – *Xanthobacter autotrophicus* 7C$^T$; XOG – *Xanthobacter oligotrophicus* 23A; XTO – *Xanthobacter toluenivorans* T101$^T$; XLU – *Xanthobacter lutulentifluminis* V3C-3$^T$; XNS – *Xanthobacter nonsaccharivorans* 14g$^T$; XAM – Xanthobacter aminoxidans 14a$^T$; XFL – Xanthobacter flavus 301$^T$; XWE – Xanthobacter wiegelii RH 10$^T$; XPS – *Xanthobacter pseudotagetidis* KA$^T$; XTA – *Xanthobacter tagetidis* TagT2C$^T$; XCO – *Xanthobacter cornucopiae* V4C-4$^T$; XAG – *Xanthobacter agilis* SA35$^T$; XSE – *Xanthobacter sediminis* V8C-5$^T$; XAL – *Xanthobacter albus* V0C-6$^T$; XVA – *Xanthobacter variabilis* V4C-8$^T$; RFI – *Roseixanthobacter finlandensis* VTT E-85241$^T$; RLI – *Roseixanthobacter liquoris* VTT E-85238$^T$; RGL – *Roseixanthobacter glucoisosaccharinivorans* VTT E-85242$^T$; RPP – *Roseixanthobacter pseudopolyaromaticivorans* VTT E-85240$^T$; RSY – *Roseixanthobacter psychrophilus* W30$^T$.

| Condition | Clade I | | | | Clade IIA | | Clade IIB | | | Clade III | | | | | Clade IV | | Rosei | | | | |
|---|---|---|---|---|---|---|---|---|---|---|---|---|---|---|---|---|---|---|---|---|---|
| | XVE | XAU | XOG | XTO | XLU | XNS | XAM | XFL | XWE | XCO | XAG | XSE | XAL | XVA | XPS | XTA | RFI | RLI | RGL | RPP | RSY |
| 0% NaCl | 7.2 | 7 | 7.2 | 5.7 | 7.1 | 7.1 | 7 | 6.6 | 6.9 | 5.8 | 4.8 | 5.7 | 5.7 | 6.5 | 7 | 5.1 | 6.4 | 4.1 | 6.1 | 6.3 | 4.2 |
| 0.5% NaCl | 6.9 | 7.3 | 6.7 | 5 | 7 | 7 | 6.7 | 6.4 | 6.5 | 6.5 | 5.5 | 5.8 | 6.1 | 6.9 | 6.9 | 6.5 | 6.2 | 4.7 | 6.4 | 6.3 | 6.8 |
| 1% NaCl | 6.9 | 6 | 2.9 | 1.1 | 4.9 | 6.5 | 6.8 | 6.3 | 6.6 | 6.2 | 4.5 | 4.9 | 5.2 | 7.1 | 6.6 | 6.1 | 6.4 | 3.6 | 6.5 | 2.7 | 4.6 |
| 1.5% NaCl | 4.1 | 2.2 | 1.7 | 1 | 3.8 | 3.8 | 5.7 | 4.9 | 5.6 | 5.1 | 2.2 | 2.7 | 3.9 | 6.1 | 6.2 | 5.2 | 4.5 | 1.7 | 6.4 | 4.2 | 2.8 |
| 2% NaCl | 1.8 | 2.2 | 2.2 | 0.8 | 1.8 | 2.1 | 4.8 | 2.9 | 2 | 1.1 | 0.3 | 1.7 | 1.6 | 5.5 | 5.6 | 4.9 | 2.2 | 0.3 | 5.1 | 0.9 | 1 |
| 3% NaCl | 3.9 | 2.2 | 2.2 | 0.4 | 0.8 | 1.8 | 1.9 | 1.7 | 0.3 | 0.7 | 0 | 2.6 | 0.6 | 4.3 | 3.4 | 2.4 | 0 | 0.1 | 1 | 0.2 | 0.1 |
| 4% NaCl | 0.7 | 1.9 | 0.6 | 0.2 | 0.8 | 0.7 | 0.7 | 1.6 | 0 | 0 | 0.2 | 1.3 | 1.2 | 1.4 | 0.2 | 0.2 | 0.1 | 0.1 | 0.7 | 0 | 0 |
| 5% NaCl | 0.5 | 1.7 | 0.6 | 0.1 | 1.3 | 1.3 | 0.3 | 0.9 | 0.2 | 0 | 0 | 0.3 | 0 | 1.1 | 0.1 | 0.2 | 0.9 | 0.4 | 0.7 | 0 | 0 |
| 6% NaCl | 0.1 | 0.4 | 0.2 | 0 | 0.8 | 0 | 0 | 0.1 | 0 | 0 | 0 | 1.2 | 0 | 1.5 | 0 | 0 | 0 | 0 | 0 | 0 | 0 |

**Supplementary Table 12 –Xanthobacter and Roseixanthobacter gen. nov. can grow heterotrophically in different rich media (Media 1/M1, double concentrated Media 1/2xM1, and R2A).** Numbers indicate the number of doublings from inoculation density ($OD_{600}$ 0.01) that have occurred based on optical density measurements. *Xanthobacter oligotrophicus* (XOG) is represented here by strain 23A instead of type strain 29k$^T$. Abbreviations are: XVE – *Xanthobacter versatilis* Py2$^T$; XAU – *Xanthobacter autotrophicus* 7C$^T$; XOG – *Xanthobacter oligotrophicus* 23A; XTO – *Xanthobacter toluenivorans* T101$^T$; XLU – *Xanthobacter lutulentifluminis* V3C-3$^T$; XNS – *Xanthobacter nonsaccharivorans* 14g$^T$; XAM – Xanthobacter aminoxidans 14a$^T$; XFL – Xanthobacter flavus 301$^T$; XWE – Xanthobacter wiegelii RH 10$^T$; XPS – *Xanthobacter pseudotagetidis* KA$^T$; XTA – *Xanthobacter tagetidis* TagT2C$^T$; XCO – *Xanthobacter cornucopiae* V4C-4$^T$; XAG – *Xanthobacter agilis* SA35$^T$; XSE – *Xanthobacter sediminis* V8C-5$^T$; XAL – *Xanthobacter albus* V0C-6$^T$; XVA – *Xanthobacter variabilis* V4C-8$^T$; RFI – *Roseixanthobacter finlandensis* VTT E-85241$^T$; RLI – *Roseixanthobacter liquoris* VTT E-85238$^T$; RGL – *Roseixanthobacter glucoisosaccharinivorans* VTT E-85242$^T$; RPP – *Roseixanthobacter pseudopolyaromaticivorans* VTT E-85240$^T$; RSY – *Roseixanthobacter psychrophilus* W30$^T$.

| Condition | Clade I | | | | Clade IIA | | Clade IIB | | | Clade III | | | | | Clade IV | | Rosei | | | | |
|---|---|---|---|---|---|---|---|---|---|---|---|---|---|---|---|---|---|---|---|---|---|
| | XVE | XAU | XOG | XTO | XLU | XNS | XAM | XFL | XWE | XCO | XAG | XSE | XAL | XVA | XPS | XTA | RFI | RLI | RGL | RPP | RSY |
| 2xM1 | 7.2 | 7 | 7.2 | 5.7 | 7.1 | 7.1 | 7 | 6.6 | 6.9 | 5.8 | 4.8 | 5.7 | 5.7 | 6.5 | 7 | 5.1 | 6.4 | 4.1 | 6.1 | 6.3 | 4.2 |
| M1 | 6.4 | 6.9 | 6.2 | 5 | 6.4 | 6.3 | 6.2 | 5.9 | 6 | 5.2 | 4 | 5 | 5.4 | 5.5 | 6.3 | 4.4 | 5.7 | 5 | 5.3 | 5.5 | 4.1 |
| R2A | 6.4 | 6.3 | 6.3 | 5.9 | 5.8 | 5.6 | 5.7 | 5.6 | 6.1 | 5 | 5.1 | 5.4 | 5.3 | 5.6 | 5.5 | 5.6 | 5.2 | 5.9 | 5 | 5.3 | 5.6 |

**Supplementary Table 13 – Carbon substrate utilization testing for Xanthobacter and Roseixanthobacter gen. nov. type strains.** Numbers indicate the number of doublings from inoculation density ($OD_{600}$ 0.01) that have occurred based on optical density measurements. Substrates were tested at a concentration of 0.5% w/v or v/v in minimal media supplemented with ammonium, vitamins, and a trace metal mixture (media 4M) and are recorded in the first column under each species. For mixed carbon testing, the base minimal media also contained 0.01% w/v succinate (or pyruvate for *X. toluenivorans*) and 0.5% w/v or v/v substrate to see if strains could grow on a carbon source if they were provided a small amount of a preferred carbon substrate and total doublings in this condition is recorded in the second column under each species. The third column under each species is the doublings observed in the mixed carbon condition subtracted from the doublings observed in a 0.01% w/v succinate condition, giving the amount of doublings due to the presence of the 0.5% w/v or v/v other carbon source. Background (optical density of each strain in the minimal media without a carbon substrate provided) was subtracted from these doubling values. *Xanthobacter oligotrophicus* (XOG) is represented here by strain 23A instead of type strain 29k$^T$. Abbreviations are: XVE – *Xanthobacter versatilis* Py2$^T$; XAU – *Xanthobacter autotrophicus* 7C$^T$; XOG – *Xanthobacter oligotrophicus* 23A; XTO – *Xanthobacter toluenivorans* T101$^T$; XLU – *Xanthobacter lutulentifluminis* V3C-3$^T$; XNS – *Xanthobacter nonsaccharivorans* 14g$^T$; XAM – Xanthobacter aminoxidans 14a$^T$; XFL – Xanthobacter flavus 301$^T$; XWE – Xanthobacter wiegelii RH 10$^T$; XPS – *Xanthobacter pseudotagetidis* KA$^T$; XTA – *Xanthobacter tagetidis* TagT2C$^T$; XCO – *Xanthobacter cornucopiae* V4C-4$^T$; XAG – *Xanthobacter agilis* SA35$^T$; XSE – *Xanthobacter sediminis* V8C-5$^T$; XAL – *Xanthobacter albus* V0C-6$^T$; XVA – *Xanthobacter variabilis* V4C-8$^T$; RFI – *Roseixanthobacter finlandensis* VTT E-85241$^T$; RLI – *Roseixanthobacter liquoris* VTT E-85238$^T$; RGL – *Roseixanthobacter glucoisosaccharinivorans* VTT E-85242$^T$; RPP – *Roseixanthobacter pseudopolyaromaticivorans* VTT E-85240$^T$; RSY – *Roseixanthobacter psychrophilus* W30$^T$.

| Type | Substrate | Clade I | | | | | | | | | | | |
|---|---|---|---|---|---|---|---|---|---|---|---|---|---|
| | | XVE | | | XAU | | | XOG | | | XTO | | |
| | | Substrate Alone | Mixed Substrate and 0.01% Succinate | Mixed Growth Due to Substrate Alone | Substrate Alone | Mixed Substrate and 0.01% Succinate | Mixed Growth Due to Substrate Alone | Substrate Alone | Mixed Substrate and 0.01% Succinate | Mixed Growth Due to Substrate Alone | Substrate Alone | Mixed Substrate and 0.01% Pyruvate | Mixed Growth Due to Substrate Alone |
| Alcohol Testing | Ethanol | 7.6 | 7.9 | 5.3 | 7.4 | 7.8 | 4.7 | 6.8 | 7.3 | 2.7 | 1.6 | 0 | 0 |
| | Isopropanol | 5.3 | 5.5 | 2.9 | 6.7 | 5.9 | 2.8 | 6.8 | 5.7 | 1.1 | 1.5 | 0 | 0 |
| | Methanol | 3.7 | 3.9 | 1.3 | 6.8 | 6.7 | 3.6 | 6.4 | 6 | 1.4 | 0 | 0 | 0 |
| | Propanol | 7.5 | 7.6 | 5 | 6.5 | 6.8 | 3.7 | 7.1 | 7.1 | 2.5 | 2.5 | 0 | 0 |
| Amino Acid Testing | L-Alanine | 0 | 0 | 0 | 0.4 | 2.8 | 0 | 0 | 2.7 | 0 | 0 | 1.9 | 1.9 |
| | L-Arginine | 0 | 3.3 | 0.7 | 0 | 3.1 | 0 | 0 | 3.2 | 0 | 0 | 0 | 0 |
| | L-Aspartate | 0 | 0 | 0 | 0 | 0 | 0 | 0 | 0 | 0 | 0 | 0 | 0 |
| | L-Cysteine | 2.2 | 3 | 0.4 | 0 | 0 | 0 | 2.3 | 2.7 | 0 | 0.8 | 2.6 | 2.6 |
| | L-Glutamine | 5.9 | 5.9 | 3.3 | 7.2 | 7.1 | 4 | 6.1 | 6.3 | 1.7 | 0.1 | 0 | 0 |
| | L-Histidine | 3.9 | 3.4 | 0.8 | 0 | 3.2 | 0.1 | 0 | 3.4 | 0 | 0 | 0 | 0 |
| | L-Isoleucine | 2.5 | 0 | 0 | 2.9 | 0 | 0 | 2.6 | 0 | 0 | 0 | 0 | 0 |
| | L-Leucine | 0 | 0 | 0 | 0 | 0 | 0 | 0 | 0 | 0 | 0 | 0 | 0 |
| | L-Lysine | 0 | 2.7 | 0.1 | 0 | 3.3 | 0.2 | 0 | 3.1 | 0 | 0 | 0 | 0 |
| | L-Methionine | 0 | 1.4 | 0 | 0 | 2.6 | 0 | 0 | 4 | 0 | 0 | 0 | 0 |
| | L-Ornithine | 0 | 3 | 0.4 | 0 | 2.6 | 0 | 0 | 2.5 | 0 | 0 | 0 | 0 |
| | L-Phenylalanine | 1.1 | 2.4 | 0 | 2.3 | 2.8 | 0 | 0 | 1.2 | 0 | 0 | 0 | 0 |
| | L-Proline | 1.5 | 3.9 | 1.3 | 1.8 | 4.4 | 1.3 | 0 | 5.6 | 1 | 0 | 0 | 0 |
| | L-Serine | 0 | 2 | 0 | 0.5 | 3 | 0 | 0 | 2.7 | 0 | 0 | 1.3 | 1.3 |
| | L-Threonine | 2.1 | 2.3 | 0 | 0 | 0.8 | 0 | 0 | 1.3 | 0 | 0 | 0 | 0 |
| | L-Tryptophan | 0 | 3.5 | 0.9 | 0 | 0 | 0 | 0 | 0.3 | 0 | 0 | 0 | 0 |
| | L-Valine | 0 | 3.7 | 1.1 | 0 | 4.7 | 1.6 | 0 | 3.3 | 0 | 0 | 0 | 0 |
| Metabolite Testing | 2-Oxogluturate | 0 | 0 | 0 | 0 | 0 | 0 | 0 | 0 | 0 | 0 | 0 | 0 |
| | Acetate | 3.9 | 3.1 | 0.5 | 6 | 5.4 | 2.3 | 2.3 | 1.6 | 0 | 0 | 0 | 0 |
| | Alpha-Aminobutyrate | 0 | 0 | 0 | 0 | 0 | 0 | 0 | 0 | 0 | 0 | 0 | 0 |
| | Beta-Hydroxybutyrate | 7.2 | 7.1 | 4.5 | 7.4 | 7.3 | 4.2 | 7.4 | 7.1 | 2.5 | 1.4 | 0 | 0 |
| | Butyrate | 6.8 | 6.8 | 4.2 | 7.7 | 7.7 | 4.6 | 5.9 | 4.9 | 0.3 | 0 | 0 | 0 |
| | Citrate | 6.1 | 5.6 | 3 | 6.1 | 5.9 | 2.8 | 5 | 4.7 | 0.1 | 0 | 0 | 0 |
| | D-Malate | 6.4 | 6.1 | 3.5 | 6.7 | 6.7 | 3.6 | 5.5 | 4.5 | 0 | 0 | 0 | 0 |
| | D-Mandelate | 0 | 0 | 0 | 0 | 0 | 0 | 0 | 0 | 0 | 0 | 0 | 0 |
| | D-Tartrate | 0 | 0 | 0 | 0 | 0 | 0 | 0 | 0 | 0 | 0 | 0 | 0 |
| | Formate | 3.9 | 3.6 | 1 | 2.8 | 3.5 | 0.4 | 4.2 | 2.9 | 0 | 0 | 0 | 0 |
| | Fumarate | 5.4 | 4.8 | 2.2 | 6.4 | 5.7 | 2.6 | 4.9 | 4 | 0 | 0 | 0 | 0 |
| | Gluconate | 7.1 | 6.9 | 4.3 | 7.6 | 7.3 | 4.2 | 7 | 6.7 | 2.1 | 2.4 | 0.6 | 0.6 |
| | D-Glucuronic Acid | 7.4 | 5.2 | 2.6 | 0 | 3.8 | 0.7 | 6.9 | 7.5 | 2.9 | 0 | 0 | 0 |
| | Glutarate | 0 | 0 | 0 | 0 | 0 | 0 | 0 | 0 | 0 | 0 | 0 | 0 |
| | Glyoxylate | 0 | 0 | 0 | 0 | 0 | 0 | 0 | 0 | 0 | 0 | 0 | 0 |
| | L-Malate | 5.4 | 5 | 2.4 | 5.4 | 6.1 | 3 | 4.8 | 4.9 | 0.3 | 0 | 0 | 0 |
| | L-Mandelate | 0 | 0 | 0 | 0 | 0 | 0 | 0 | 0 | 0 | 0 | 0 | 0 |
| | L-Tartrate | 0 | 3.3 | 0.7 | 0 | 3.2 | 0.1 | 0 | 3.8 | 0 | 0 | 0 | 0 |
| | Malonate | 0 | 0 | 0 | 0 | 0 | 0 | 0 | 0 | 0 | 0 | 0 | 0 |
| | N-Acetyl-Glucosamine | 0 | 2.9 | 0.3 | 0.3 | 3.3 | 0.2 | 0 | 4.5 | 0 | 0 | 0 | 0 |
| | Oxalate | 0 | 0 | 0 | 0 | 0 | 0 | 0 | 0 | 0 | 0 | 0 | 0 |
| | Propionate | 0 | 1.8 | 0 | 7.1 | 7.1 | 4 | 0 | 0 | 0 | 0 | 0 | 0 |
| | Pyruvate | 6.6 | 6.3 | 3.7 | 6.2 | 6.2 | 3.1 | 5.8 | 6 | 1.4 | 4.1 | 2.3 | 2.3 |
| | Succinate | 5.3 | 5.2 | 2.6 | 6.2 | 6.6 | 3.5 | 5 | 5.1 | 0.5 | 0 | 0 | 0 |
| Other Testing | Benzoate | 0 | 0 | 0 | 0 | 0 | 0 | 0 | 0 | 0 | 0 | 0 | 0 |
| | Glycerol | 0 | 3 | 0.4 | 7.6 | 7.1 | 4 | 3.9 | 5.3 | 0.7 | 0 | 0 | 0 |
| | Salicine | 0.9 | 3.8 | 1.2 | 0 | 3.6 | 0.5 | 0 | 4.4 | 0 | 0 | 0 | 0 |
| | Urea | 0 | 2.3 | 0 | 0 | 2.6 | 0 | 0 | 2.3 | 0 | 0 | 0 | 0 |
| Sugar Testing | D-Cellobiose | 0 | 2.9 | 0.3 | 0.6 | 2.9 | 0 | 0.3 | 3.1 | 0 | 0 | 0 | 0 |
| | D-Fructose | 7.5 | 7.7 | 5.1 | 8 | 7.9 | 4.8 | 7.4 | 7.3 | 2.7 | 0 | 0 | 0 |
| | D-Glucose | 8.4 | 5.1 | 2.5 | 0 | 3.1 | 0 | 0 | 3.3 | 0 | 0 | 0 | 0 |
| | D-Mannose | 0 | 1 | 0 | 2 | 3.6 | 0.5 | 0 | 3.7 | 0 | 0 | 0 | 0 |
| | D-Raffinose | 0 | 3.3 | 0.7 | 0 | 3.7 | 0.6 | 0 | 2.9 | 0 | 0 | 0 | 0 |
| | D-Xylose | 0 | 3.4 | 0.8 | 0 | 3.5 | 0.4 | 0 | 3.3 | 0 | 0 | 0 | 0 |
| | D-Galactose | 0 | 2.8 | 0.2 | 1.3 | 2.8 | 0 | 0 | 2.7 | 0 | 0 | 0 | 0 |
| | myo-Inositol | 0 | 3 | 0.4 | 0 | 3.1 | 0 | 0 | 3.1 | 0 | 0 | 0 | 0 |
| | D-Lactose | 0 | 2.9 | 0.3 | 0 | 2.7 | 0 | 0 | 2.7 | 0 | 0 | 0 | 0 |
| | L-Arabinose | 3.7 | 2.7 | 0.1 | 0 | 3.5 | 0.4 | 0 | 3.4 | 0 | 0 | 0 | 0 |
| | L-Sorbose | 0 | 2.1 | 0 | 1.6 | 3.3 | 0.2 | 0 | 3.4 | 0 | 0 | 0 | 0 |
| | L-Xylose | 0 | 3.2 | 0.6 | 0 | 3.7 | 0.6 | 0 | 3 | 0 | 0 | 0 | 0 |
| | D-Maltose | 0 | 3.2 | 0.6 | 0 | 3.1 | 0 | 0 | 4.9 | 0.3 | 0 | 0 | 0 |
| | D-Mannitol | 0 | 3.8 | 1.2 | 0 | 2.5 | 0 | 0 | 2.8 | 0 | 0 | 0 | 0 |
| | D-Melibiose | 0.3 | 3.3 | 0.7 | 0 | 3.7 | 0.6 | 0 | 3.1 | 0 | 0 | 0 | 0 |
| | L-Rhamnose | 0 | 2.8 | 0.2 | 0 | 3.1 | 0 | 0 | 3.2 | 0 | 0 | 0 | 0 |
| | D-Ribose | 0 | 3.2 | 0.6 | 1.4 | 3.9 | 0.8 | 0.5 | 3.6 | 0 | 0.9 | 0 | 0 |
| | D-Sorbitol | 3.9 | 2.7 | 0.1 | 1.5 | 3.2 | 0.1 | 0 | 3.3 | 0 | 0 | 0 | 0 |
| | Sucrose | 0 | 3.4 | 0.8 | 0 | 2.7 | 0 | 0 | 3.2 | 0 | 0 | 0 | 0 |
| | Trehalose | 0 | 3.1 | 0.5 | 0 | 2.6 | 0 | 0 | 2.4 | 0 | 0 | 0 | 0 |

| Type | Substrate | Clade IIA | | | | | | Clade IIB | | | | | | | | |
|---|---|---|---|---|---|---|---|---|---|---|---|---|---|---|---|---|
| | | XLU | | | XNS | | | XAM | | | XFL | | | XWE | | |
| | | Substrate Alone | Mixed Substrate and 0.01% Succinate | Mixed Growth Due to Substrate Alone | Substrate Alone | Mixed Substrate and 0.01% Succinate | Mixed Growth Due to Substrate Alone | Substrate Alone | Mixed Substrate and 0.01% Succinate | Mixed Growth Due to Substrate Alone | Substrate Alone | Mixed Substrate and 0.01% Succinate | Mixed Growth Due to Substrate Alone | Substrate Alone | Mixed Substrate and 0.01% Succinate | Mixed Growth Due to Substrate Alone |
| Alcohol Testing | Ethanol | 0 | 4.1 | 0.8 | 7.5 | 7.4 | 5.5 | 6.9 | 6.7 | 4.6 | 7.6 | 7.4 | 5.3 | 7.3 | 7.6 | 4.9 |
| | Isopropanol | 1.5 | 5.7 | 2.4 | 4.6 | 4.6 | 2.7 | 4.7 | 4.7 | 2.6 | 4.6 | 4.4 | 2.3 | 5.9 | 6.7 | 4 |
| | Methanol | 0 | 2.5 | 0 | 0 | 2.4 | 0.5 | 6.3 | 3.7 | 1.6 | 5.4 | 5.5 | 3.4 | 6.5 | 6.3 | 3.6 |
| | Propanol | 0.2 | 4.1 | 0.8 | 7.2 | 7.1 | 5.2 | 7.7 | 7.5 | 5.4 | 7.7 | 7.6 | 5.5 | 7.1 | 7.7 | 5 |
| Amino Acid Testing | L-Alanine | 0 | 2.9 | 0 | 6.4 | 6.4 | 4.5 | 6.9 | 7.3 | 5.2 | 6.8 | 6.6 | 4.5 | 7.1 | 6.8 | 4.1 |
| | L-Arginine | 0 | 5.2 | 1.9 | 0 | 1.6 | 0 | 0 | 2.3 | 0.2 | 0 | 2.3 | 0.2 | 2.3 | 2.8 | 0.1 |
| | L-Aspartate | 0 | 0 | 0 | 0 | 0 | 0 | 0 | 0 | 0 | 0 | 0 | 0 | 0 | 0 | 0 |
| | L-Cysteine | 0 | 4.3 | 1 | 2.6 | 0 | 0 | 0 | 0 | 0 | 0 | 0 | 0 | 5.2 | 0 | 0 |
| | L-Glutamine | 7.1 | 6.9 | 3.6 | 6.2 | 6.3 | 4.4 | 6.3 | 5.7 | 3.6 | 6.7 | 6.2 | 4.1 | 6.8 | 6.7 | 4 |
| | L-Histidine | 0 | 4.1 | 0.8 | 1.1 | 1.8 | 0 | 2.2 | 4 | 1.9 | 0.1 | 2.7 | 0.6 | 1.3 | 4 | 1.3 |
| | L-Isoleucine | 2.9 | 0 | 0 | 1.9 | 0 | 0 | 1.9 | 0 | 0 | 0 | 1.8 | 0 | 1 | 0 | 0 |
| | L-Leucine | 0 | 0 | 0 | 0 | 0 | 0 | 0 | 0 | 0 | 0 | 0 | 0 | 0 | 0 | 0 |
| | L-Lysine | 0 | 3.9 | 0.6 | 0 | 0.6 | 0 | 1 | 2.8 | 0.7 | 0 | 1.9 | 0 | 0 | 2.7 | 0 |
| | L-Methionine | 0 | 3.2 | 0 | 0 | 0 | 0 | 0 | 1.2 | 0 | 0 | 1.2 | 0 | 0 | 1.5 | 0 |
| | L-Ornithine | 0 | 2.9 | 0 | 0 | 0.9 | 0 | 0 | 2.7 | 0.6 | 0 | 2.3 | 0.2 | 0 | 2.7 | 0 |
| | L-Phenylalanine | 0 | 0 | 0 | 0 | 1.2 | 0 | 0 | 0 | 0 | 1 | 1.4 | 0 | 0 | 0 | 0 |
| | L-Proline | 1.2 | 3.8 | 0.5 | 0 | 1.2 | 0 | 0 | 1.4 | 0 | 1 | 1.8 | 0 | 4.2 | 5.9 | 3.2 |
| | L-Serine | 0 | 0 | 0 | 0 | 1.9 | 0 | 0 | 3.3 | 1.2 | 1.2 | 3.2 | 1.1 | 0 | 3.1 | 0.4 |
| | L-Threonine | 2.5 | 0.1 | 0 | 0 | 1.3 | 0 | 0 | 1.9 | 0 | 0 | 0 | 0 | 0 | 0 | 0 |
| | L-Tryptophan | 0 | 2 | 0 | 0 | 0 | 0 | 0 | 1.3 | 0 | 0 | 0 | 0 | 0 | 0 | 0 |
| | L-Valine | 0 | 3.1 | 0 | 0 | 3.2 | 1.3 | 2.1 | 4.5 | 2.4 | 0.7 | 3.7 | 1.6 | 0 | 2.6 | 0 |
| Metabolite Testing | 2-Oxogluturate | 0 | 0 | 0 | 0 | 0 | 0 | 0 | 0 | 0 | 0 | 0 | 0 | 0 | 0 | 0 |
| | Acetate | 5.9 | 5.5 | 2.2 | 6.6 | 5.9 | 4 | 6 | 5.9 | 3.8 | 6.5 | 6.5 | 4.4 | 6.5 | 6.2 | 3.5 |
| | Alpha-Aminobutyrate | 0 | 0 | 0 | 0 | 0 | 0 | 0 | 0 | 0 | 0 | 0 | 0 | 0 | 0 | 0 |
| | Beta-Hydroxybutyrate | 7.5 | 7.4 | 4.1 | 7.7 | 7.5 | 5.6 | 8 | 7.4 | 5.3 | 7.6 | 7.6 | 5.5 | 7.8 | 7.8 | 5.1 |
| | Butyrate | 7.5 | 7.2 | 3.9 | 7.9 | 7.9 | 6 | 3.7 | 7.4 | 5.3 | 7.7 | 7.6 | 5.5 | 4.1 | 7.9 | 5.2 |
| | Citrate | 0 | 2.4 | 0 | 5.8 | 5.6 | 3.7 | 6.8 | 6 | 3.9 | 5.4 | 5.2 | 3.1 | 5.5 | 5.2 | 2.5 |
| | D-Malate | 0 | 2.9 | 0 | 7.1 | 6.9 | 5 | 7 | 6.5 | 4.4 | 7 | 6.9 | 4.8 | 6.9 | 6.7 | 4 |
| | D-Mandelate | 0 | 0 | 0 | 0 | 0 | 0 | 0 | 0 | 0 | 0 | 0 | 0 | 0 | 0 | 0 |
| | D-Tartrate | 0 | 0 | 0 | 0 | 0 | 0 | 0 | 0 | 0 | 0 | 0 | 0 | 0 | 0 | 0 |
| | Formate | 4.9 | 3.9 | 0.6 | 3.5 | 3.2 | 1.3 | 0 | 0 | 0 | 3.7 | 3.9 | 1.8 | 0 | 3.5 | 0.8 |
| | Fumarate | 6.4 | 5.7 | 2.4 | 7.1 | 5.6 | 3.7 | 7.1 | 6.6 | 4.5 | 6.9 | 6.3 | 4.2 | 7.1 | 6.6 | 3.9 |
| | Gluconate | 7.7 | 7.7 | 4.4 | 7.8 | 7.5 | 5.6 | 8 | 7.3 | 5.2 | 7.9 | 7.7 | 5.6 | 7.8 | 7.7 | 5 |
| | D-Glucuronic Acid | 7.6 | 7.4 | 4.1 | 0 | 2.1 | 0.2 | 1.3 | 2.7 | 0.6 | 0 | 3 | 0.9 | 6.2 | 6.1 | 3.4 |
| | Glutarate | 0 | 0 | 0 | 0 | 0 | 0 | 0 | 0 | 0 | 0 | 0 | 0 | 0 | 0 | 0 |
| | Glyoxylate | 0 | 0 | 0 | 0 | 0 | 0 | 0 | 0 | 0 | 0 | 0 | 0 | 0 | 0 | 0 |
| | L-Malate | 6.4 | 5.6 | 2.3 | 6.1 | 5.8 | 3.9 | 6.7 | 6.3 | 4.2 | 6.6 | 6.4 | 4.3 | 6.8 | 6.4 | 3.7 |
| | L-Mandelate | 0 | 0 | 0 | 0 | 0 | 0 | 0 | 0 | 0 | 0 | 0 | 0 | 0 | 0 | 0 |
| | L-Tartrate | 0 | 5.4 | 2.1 | 0 | 1.4 | 0 | 0 | 2 | 0 | 0 | 2 | 0 | 0 | 2.5 | 0 |
| | Malonate | 0 | 0 | 0 | 0 | 0 | 0 | 0 | 0 | 0 | 0 | 0 | 0 | 0 | 0 | 0 |
| | N-Acetyl-Glucosamine | 0 | 3.4 | 0.1 | 0 | 2.5 | 0.6 | 0 | 2.1 | 0 | 0 | 2.2 | 0.1 | 0 | 2.3 | 0 |
| | Oxalate | 0 | 0 | 0 | 0 | 0 | 0 | 0 | 0 | 0 | 0 | 0 | 0 | 0 | 0 | 0 |
| | Propionate | 7.1 | 6.8 | 3.5 | 0 | 0 | 0 | 0 | 6.3 | 4.2 | 7.4 | 7.3 | 5.2 | 0 | 7.7 | 5 |
| | Pyruvate | 6.7 | 6.4 | 3.1 | 6.6 | 6.5 | 4.6 | 7 | 6.3 | 4.2 | 6.3 | 6.4 | 4.3 | 7 | 6.6 | 3.9 |
| | Succinate | 6.2 | 5.5 | 2.2 | 6.6 | 6.6 | 4.7 | 6.9 | 6.6 | 4.5 | 6.6 | 6.7 | 4.6 | 7.3 | 6.8 | 4.1 |
| Other Testing | Benzoate | 0 | 0 | 0 | 0 | 0 | 0 | 0 | 0 | 0 | 0 | 0 | 0 | 0 | 0 | 0 |
| | Glycerol | 0 | 4.6 | 1.3 | 0 | 2.8 | 0.9 | 8.7 | 8.6 | 6.5 | 4.7 | 4.8 | 2.7 | 8.8 | 8.8 | 6.1 |
| | Salicine | 0 | 3.4 | 0.1 | 0 | 2 | 0.1 | 0 | 2 | 0 | 0 | 1.8 | 0 | 0 | 2.6 | 0 |
| | Urea | 0 | 2.3 | 0 | 0 | 0 | 0 | 0 | 1.6 | 0 | 0 | 1.9 | 0 | 0 | 1.3 | 0 |
| Sugar Testing | D-Cellobiose | 0 | 1.9 | 0 | 0 | 2.1 | 0.2 | 0 | 2.2 | 0.1 | 0 | 2.1 | 0 | 0.6 | 2.7 | 0 |
| | D-Fructose | 0 | 2.3 | 0 | 0 | 2 | 0.1 | 0 | 2 | 0 | 0 | 2.4 | 0.3 | 0 | 2.5 | 0 |
| | D-Glucose | 0 | 2.5 | 0 | 0 | 1.6 | 0 | 0.4 | 2.6 | 0.5 | 0 | 2.3 | 0.2 | 0.6 | 3 | 0.3 |
| | D-Mannose | 0 | 3.1 | 0 | 0 | 3 | 1.1 | 5.1 | 5.4 | 3.3 | 2.1 | 3.9 | 1.8 | 1.8 | 3.9 | 1.2 |
| | D-Raffinose | 0 | 2.5 | 0 | 0 | 1 | 0 | 0 | 1.8 | 0 | 0 | 1.7 | 0 | 0 | 2.5 | 0 |
| | D-Xylose | 0 | 4 | 0.7 | 0 | 1.4 | 0 | 0 | 2 | 0 | 0 | 2.1 | 0 | 0 | 2.7 | 0 |
| | D-Galactose | 0 | 2.7 | 0 | 0 | 1.1 | 0 | 0 | 1.4 | 0 | 0 | 1.7 | 0 | 0 | 2.5 | 0 |
| | myo-Inositol | 0 | 2.5 | 0 | 0 | 0.9 | 0 | 0 | 1.5 | 0 | 0 | 1.7 | 0 | 0 | 2.3 | 0 |
| | D-Lactose | 0 | 3.3 | 0 | 0 | 1.1 | 0 | 0 | 2.3 | 0.2 | 0 | 1.5 | 0 | 0 | 2.2 | 0 |
| | L-Arabinose | 0 | 2.9 | 0 | 0 | 1.4 | 0 | 0 | 1.8 | 0 | 0 | 2.1 | 0 | 0 | 2.4 | 0 |
| | L-Sorbose | 2.4 | 3.3 | 0 | 0 | 2 | 0.1 | 0.5 | 2.3 | 0.2 | 1 | 2.6 | 0.5 | 0 | 2.3 | 0 |
| | L-Xylose | 0 | 2.3 | 0 | 0 | 2 | 0.1 | 0 | 2 | 0 | 0 | 2.2 | 0.1 | 0 | 2.8 | 0.1 |
| | D-Maltose | 0 | 3.3 | 0 | 0 | 1.1 | 0 | 0 | 1.8 | 0 | 0 | 1.8 | 0 | 0 | 2.7 | 0 |
| | D-Mannitol | 0 | 3.8 | 0.5 | 0 | 1.1 | 0 | 0 | 1.7 | 0 | 0 | 1.7 | 0 | 0 | 2.3 | 0 |
| | D-Melibiose | 0 | 2.5 | 0 | 0 | 1.7 | 0 | 0 | 2.1 | 0 | 0 | 1.9 | 0 | 0 | 2.7 | 0 |
| | L-Rhamnose | 0 | 0.1 | 0 | 0 | 1.2 | 0 | 0 | 2.3 | 0.2 | 0 | 1.6 | 0 | 0 | 2.4 | 0 |
| | D-Ribose | 1.4 | 2.7 | 0 | 0 | 2.1 | 0.2 | 1.2 | 2.8 | 0.7 | 2.2 | 2.4 | 0.3 | 0.2 | 2.7 | 0 |
| | D-Sorbitol | 0 | 5.6 | 2.3 | 0 | 1.1 | 0 | 0 | 1.6 | 0 | 0 | 1.7 | 0 | 0 | 2.3 | 0 |
| | Sucrose | 0 | 1.2 | 0 | 0 | 1 | 0 | 0 | 1.7 | 0 | 0 | 1.8 | 0 | 0 | 2.5 | 0 |
| | Trehalose | 0 | 2.7 | 0 | 0 | 1.1 | 0 | 0 | 1.7 | 0 | 0 | 1.8 | 0 | 0 | 2.3 | 0 |

| Type | Substrate | Clade III | | | | | | | | | | | | | | |
|---|---|---|---|---|---|---|---|---|---|---|---|---|---|---|---|---|
| | | XCO | | | XAG | | | XSE | | | XAL | | | XVA | | |
| | | Substrate Alone | Mixed Substrate and 0.01% Succinate | Mixed Growth Due to Substrate Alone | Substrate Alone | Mixed Substrate and 0.01% Succinate | Mixed Growth Due to Substrate Alone | Substrate Alone | Mixed Substrate and 0.01% Succinate | Mixed Growth Due to Substrate Alone | Substrate Alone | Mixed Substrate and 0.01% Succinate | Mixed Growth Due to Substrate Alone | Substrate Alone | Mixed Substrate and 0.01% Succinate | Mixed Growth Due to Substrate Alone |
| Alcohol Testing | Ethanol | 7.5 | 7.7 | 5.4 | 6.4 | 7.1 | 5.1 | 7.3 | 7.2 | 5 | 7.4 | 7.5 | 5.1 | 7.8 | 7.5 | 5.7 |
| | Isopropanol | 7.1 | 6.4 | 4.1 | 4.2 | 4.9 | 2.9 | 5.3 | 6.6 | 4.4 | 6.1 | 5.5 | 3.1 | 7.4 | 5.1 | 3.3 |
| | Methanol | 4.4 | 4.6 | 2.3 | 5.6 | 4.8 | 2.8 | 6.2 | 6 | 3.8 | 4.9 | 5.6 | 3.2 | 6.9 | 5.8 | 4 |
| | Propanol | 7.4 | 7.8 | 5.5 | 6.1 | 7 | 5 | 6.7 | 7.2 | 5 | 7.2 | 7.3 | 4.9 | 7.5 | 6.7 | 4.9 |
| Amino Acid Testing | L-Alanine | 7.1 | 7.1 | 4.8 | 4.4 | 5.3 | 3.3 | 7 | 6 | 3.8 | 6.6 | 6 | 3.6 | 7.4 | 7.1 | 5.3 |
| | L-Arginine | 2 | 2.2 | 0 | 1.5 | 2.1 | 0.1 | 2.6 | 2.8 | 0.6 | 1.8 | 2.8 | 0.4 | 2 | 2.4 | 0.6 |
| | L-Aspartate | 0 | 0 | 0 | 0 | 0 | 0 | 0 | 0 | 0 | 0 | 0 | 0 | 0 | 0 | 0 |
| | L-Cysteine | 2.7 | 1.3 | 0 | 3.2 | 2.6 | 0.6 | 2.5 | 3.3 | 1.1 | 0.1 | 2.1 | 0 | 3.1 | 3.4 | 1.6 |
| | L-Glutamine | 6.1 | 6.1 | 3.8 | 4.1 | 4.6 | 2.6 | 6.9 | 6 | 3.8 | 6.8 | 6.8 | 4.4 | 6.3 | 4.8 | 3 |
| | L-Histidine | 3 | 2.3 | 0 | 2.3 | 4 | 2 | 3.7 | 3 | 0.8 | 3.8 | 3 | 0.6 | 5.2 | 3.5 | 1.7 |
| | L-Isoleucine | 0 | 0 | 0 | 0 | 0 | 0 | 0 | 0 | 0 | 0 | 0 | 0 | 0 | 0 | 0 |
| | L-Leucine | 0 | 0 | 0 | 0 | 0 | 0 | 0 | 0 | 0 | 0 | 0 | 0 | 0 | 0 | 0 |
| | L-Lysine | 2.4 | 1.6 | 0 | 2.5 | 2.4 | 0.4 | 2.2 | 2.6 | 0.4 | 2.3 | 3.7 | 1.3 | 3.3 | 1.3 | 0 |
| | L-Methionine | 0 | 1.3 | 0 | 0 | 1.5 | 0 | 0.9 | 1.6 | 0 | 0 | 1.7 | 0 | 0 | 0.7 | 0 |
| | L-Ornithine | 0 | 0 | 0 | 0 | 2.3 | 0.3 | 0.3 | 2.4 | 0.2 | 0 | 1.9 | 0 | 0 | 1.6 | 0 |
| | L-Phenylalanine | 0.5 | 0 | 0 | 0 | 0 | 0 | 0 | 0 | 0 | 0 | 1.2 | 0 | 0 | 0 | 0 |
| | L-Proline | 1.7 | 3 | 0.7 | 1.2 | 3.1 | 1.1 | 2.7 | 2.4 | 0.2 | 2.6 | 2.7 | 0.3 | 2.9 | 3 | 1.2 |
| | L-Serine | 2.9 | 3 | 0.7 | 1.1 | 3.1 | 1.1 | 2.8 | 2.4 | 0.2 | 2.3 | 2.7 | 0.3 | 3 | 2.8 | 1 |
| | L-Threonine | 0 | 0 | 0 | 0 | 0 | 0 | 0 | 0 | 0 | 0 | 0 | 0 | 1 | 0 | 0 |
| | L-Tryptophan | 0 | 2.2 | 0 | 1.3 | 2.8 | 0.8 | 2.3 | 1.2 | 0 | 0.1 | 2.5 | 0.1 | 3.2 | 3.8 | 2 |
| | L-Valine | 0 | 2.9 | 0.6 | 0 | 2.1 | 0.1 | 1.6 | 3.5 | 1.3 | 1.4 | 3.9 | 1.5 | 1.2 | 1.9 | 0.1 |
| Metabolite Testing | 2-Oxogluturate | 0 | 0 | 0 | 0 | 0 | 0 | 0 | 0 | 0 | 0 | 0 | 0 | 0 | 0 | 0 |
| | Acetate | 6.2 | 6.5 | 4.2 | 5.7 | 4.8 | 2.8 | 6.2 | 5.9 | 3.7 | 6.3 | 6.1 | 3.7 | 6.3 | 4.7 | 2.9 |
| | Alpha-Aminobutyrate | 0 | 0 | 0 | 0 | 0 | 0 | 0 | 0 | 0 | 0 | 0 | 0 | 0 | 0 | 0 |
| | Beta-Hydroxybutyrate | 7.4 | 7.5 | 5.2 | 7.4 | 6.9 | 4.9 | 7.5 | 7.5 | 5.3 | 7.6 | 7.6 | 5.2 | 7.8 | 7.5 | 5.7 |
| | Butyrate | 7.7 | 7.7 | 5.4 | 7.7 | 7.3 | 5.3 | 7.7 | 7.6 | 5.4 | 7.9 | 7.8 | 5.4 | 7.9 | 7.6 | 5.8 |
| | Citrate | 0 | 1.8 | 0 | 0 | 1.2 | 0 | 0 | 1.9 | 0 | 0 | 2.5 | 0.1 | 0 | 1.6 | 0 |
| | D-Malate | 2.9 | 4.9 | 2.6 | 0 | 1.7 | 0 | 0 | 0 | 0 | 0 | 0 | 0 | 0 | 0 | 0 |
| | D-Mandelate | 0 | 0 | 0 | 0 | 0 | 0 | 0 | 0 | 0 | 0 | 0 | 0 | 0 | 0 | 0 |
| | D-Tartrate | 0 | 0 | 0 | 0 | 0 | 0 | 0 | 0 | 0 | 0 | 0 | 0 | 0 | 0 | 0 |
| | Formate | 3.4 | 4.2 | 1.9 | 0.8 | 1.8 | 0 | 3.8 | 3.5 | 1.3 | 3.8 | 3.7 | 1.3 | 3.8 | 2.8 | 1 |
| | Fumarate | 6.9 | 7 | 4.7 | 5.8 | 5 | 3 | 6.3 | 6.1 | 3.9 | 6.9 | 6.4 | 4 | 7.2 | 5.3 | 3.5 |
| | Gluconate | 0 | 2.4 | 0.1 | 0.1 | 2.6 | 0.6 | 0.2 | 2.1 | 0 | 0 | 2.2 | 0 | 0 | 2.3 | 0.5 |
| | D-Glucuronic Acid | 0 | 2.3 | 0 | 0.7 | 2.7 | 0.7 | 1.2 | 2.4 | 0.2 | 0 | 2.3 | 0 | 0.8 | 2.4 | 0.6 |
| | Glutarate | 0 | 0 | 0 | 0 | 0 | 0 | 0 | 0 | 0 | 0 | 0 | 0 | 0 | 0 | 0 |
| | Glyoxylate | 0 | 0 | 0 | 0 | 0 | 0 | 0 | 0 | 0 | 0 | 0 | 0 | 0 | 0 | 0 |
| | L-Malate | 6.9 | 6.6 | 4.3 | 5.9 | 4.7 | 2.7 | 6.6 | 6.2 | 4 | 6.6 | 6.4 | 4 | 6.9 | 5.8 | 4 |
| | L-Mandelate | 0 | 0 | 0 | 0 | 0 | 0 | 0 | 0 | 0 | 0 | 0 | 0 | 0 | 0 | 0 |
| | L-Tartrate | 0 | 2.1 | 0 | 0.4 | 2.3 | 0.3 | 0 | 2.2 | 0 | 0 | 2.6 | 0.2 | 0 | 2.6 | 0.8 |
| | Malonate | 0 | 0 | 0 | 0 | 0 | 0 | 0 | 0 | 0 | 0 | 0 | 0 | 0 | 0 | 0 |
| | N-Acetyl-Glucosamine | 0 | 2.9 | 0.6 | 0.1 | 2.6 | 0.6 | 0.7 | 2.7 | 0.5 | 0 | 3.1 | 0.7 | 0.8 | 2.4 | 0.6 |
| | Oxalate | 0 | 0 | 0 | 0 | 0 | 0 | 0 | 0 | 0 | 0 | 0 | 0 | 0 | 0 | 0 |
| | Propionate | 7.4 | 7.8 | 5.5 | 0 | 7.1 | 5.1 | 7.1 | 7.2 | 5 | 7.8 | 7.5 | 5.1 | 7.6 | 7.2 | 5.4 |
| | Pyruvate | 7 | 6.7 | 4.4 | 6.5 | 5.9 | 3.9 | 6.6 | 6.5 | 4.3 | 6.8 | 6.6 | 4.2 | 7 | 5.6 | 3.8 |
| | Succinate | 6.9 | 6.8 | 4.5 | 5.9 | 4.9 | 2.9 | 6.9 | 6.3 | 4.1 | 6.7 | 6.4 | 4 | 7.2 | 5.7 | 3.9 |
| Other Testing | Benzoate | 0 | 0 | 0 | 0 | 0 | 0 | 0 | 0 | 0 | 0 | 0 | 0 | 0 | 0 | 0 |
| | Glycerol | 0 | 0 | 0 | 0 | 1.6 | 0 | 0 | 1.4 | 0 | 0 | 2.4 | 0 | 1.6 | 2.9 | 1.1 |
| | Salicine | 0 | 2.1 | 0 | 0.6 | 2.4 | 0.4 | 0 | 2.4 | 0.2 | 0 | 2.7 | 0.3 | 0 | 1.6 | 0 |
| | Urea | 0 | 0.3 | 0 | 0 | 1.7 | 0 | 0 | 1.4 | 0 | 0 | 0 | 0 | 0 | 0 | 0 |
| Sugar Testing | D-Cellobiose | 0 | 2.4 | 0.1 | 0 | 2.3 | 0.3 | 1.5 | 2.1 | 0 | 0 | 2.9 | 0.5 | 0.3 | 1.9 | 0.1 |
| | D-Fructose | 0 | 1.9 | 0 | 0 | 1.9 | 0 | 0 | 2 | 0 | 0 | 1.8 | 0 | 0 | 2.1 | 0.3 |
| | D-Glucose | 0 | 1.4 | 0 | 0 | 1.4 | 0 | 0 | 1.8 | 0 | 0 | 2 | 0 | 0 | 1 | 0 |
| | D-Mannose | 0 | 1.8 | 0 | 0.8 | 0.9 | 0 | 0 | 1.8 | 0 | 0 | 2.3 | 0 | 0 | 1.1 | 0 |
| | D-Raffinose | 0 | 2 | 0 | 0 | 1.9 | 0 | 0.1 | 2.2 | 0 | 0 | 2.1 | 0 | 0 | 1.8 | 0 |
| | D-Xylose | 0 | 1.9 | 0 | 0 | 1.9 | 0 | 0 | 1.9 | 0 | 0 | 2.3 | 0 | 0 | 1.9 | 0.1 |
| | D-Galactose | 0 | 2 | 0 | 0 | 2.1 | 0.1 | 0 | 1.8 | 0 | 0 | 2 | 0 | 0 | 1.2 | 0 |
| | myo-Inositol | 0 | 2.1 | 0 | 0 | 1.9 | 0 | 0.1 | 2 | 0 | 0 | 1.9 | 0 | 0 | 1.5 | 0 |
| | D-Lactose | 0 | 1 | 0 | 0 | 1.9 | 0 | 0.5 | 1.8 | 0 | 0 | 2.1 | 0 | 0 | 1.6 | 0 |
| | L-Arabinose | 0 | 1.6 | 0 | 0 | 1.6 | 0 | 0.6 | 1.9 | 0 | 0 | 1.8 | 0 | 0 | 1.5 | 0 |
| | L-Sorbose | 0 | 1.9 | 0 | 0 | 2.3 | 0.3 | 0.3 | 2.1 | 0 | 0 | 2.4 | 0 | 0 | 2.3 | 0.5 |
| | L-Xylose | 0 | 1.7 | 0 | 0 | 1.9 | 0 | 0 | 1.9 | 0 | 0 | 2.1 | 0 | 0 | 2 | 0.2 |
| | D-Maltose | 0 | 1.9 | 0 | 0 | 1.7 | 0 | 0 | 1.9 | 0 | 0 | 2.2 | 0 | 0 | 1.9 | 0.1 |
| | D-Mannitol | 0 | 1.9 | 0 | 0 | 1.9 | 0 | 0 | 2 | 0 | 0 | 2.2 | 0 | 0 | 1.7 | 0 |
| | D-Melibiose | 0 | 2.1 | 0 | 1 | 2.1 | 0.1 | 0 | 2.1 | 0 | 0 | 2 | 0 | 2 | 1.7 | 0 |
| | L-Rhamnose | 0 | 1.4 | 0 | 0 | 2 | 0 | 1 | 1.6 | 0 | 0 | 1.1 | 0 | 0 | 1.7 | 0 |
| | D-Ribose | 0 | 1.9 | 0 | 0 | 2.2 | 0.2 | 3.2 | 1.9 | 0 | 0 | 2.1 | 0 | 0 | 2.2 | 0.4 |
| | D-Sorbitol | 0 | 2.2 | 0 | 0 | 1.8 | 0 | 0 | 2.1 | 0 | 0 | 2.1 | 0 | 0 | 1.6 | 0 |
| | Sucrose | 0 | 1.8 | 0 | 0 | 2 | 0 | 0 | 1.9 | 0 | 0 | 2.3 | 0 | 0 | 1.7 | 0 |
| | Trehalose | 0 | 1.9 | 0 | 0 | 1.6 | 0 | 0 | 1.8 | 0 | 0 | 2 | 0 | 0 | 1.4 | 0 |

| Type | Substrate | Clade IV | | | | | |
|---|---|---|---|---|---|---|---|
| | | XPS | | | XTA | | |
| | | Substrate Alone | Mixed Substrate and 0.01% Succinate | Mixed Growth Due to Substrate Alone | Substrate Alone | Mixed Substrate and 0.01% Succinate | Mixed Growth Due to Substrate Alone |
| Alcohol Testing | Ethanol | 0 | 1.3 | 0.4 | 0 | 6.2 | 5 |
| | Isopropanol | 0 | 1.1 | 0.2 | 0 | 1.1 | 0 |
| | Methanol | 0 | 1.5 | 0.6 | 0 | 3.6 | 2.4 |
| | Propanol | 0 | 0.8 | 0 | 0 | 6.2 | 5 |
| Amino Acid Testing | L-Alanine | 0 | 2.2 | 1.3 | 0.1 | 7 | 5.8 |
| | L-Arginine | 0 | 0.8 | 0 | 0 | 0.5 | 0 |
| | L-Aspartate | 0 | 0 | 0 | 0 | 0 | 0 |
| | L-Cysteine | 0 | 1.4 | 0.5 | 0 | 3.2 | 2 |
| | L-Glutamine | 0.4 | 3.5 | 2.6 | 4.1 | 4.2 | 3 |
| | L-Histidine | 0 | 1.7 | 0.8 | 0 | 1.6 | 0.4 |
| | L-Isoleucine | 0 | 0 | 0 | 2.6 | 0 | 0 |
| | L-Leucine | 0 | 0 | 0 | 0 | 0 | 0 |
| | L-Lysine | 0 | 1.1 | 0.2 | 0 | 1.2 | 0 |
| | L-Methionine | 0 | 0.8 | 0 | 0 | 1 | 0 |
| | L-Ornithine | 0 | 1.4 | 0.5 | 0 | 1.8 | 0.6 |
| | L-Phenylalanine | 0 | 1.7 | 0.8 | 0 | 1.2 | 0 |
| | L-Proline | 0 | 3.3 | 2.4 | 2.9 | 4 | 2.8 |
| | L-Serine | 0 | 2 | 1.1 | 0 | 2.7 | 1.5 |
| | L-Threonine | 0 | 2 | 1.1 | 0 | 0 | 0 |
| | L-Tryptophan | 0 | 0.7 | 0 | 0 | 0.7 | 0 |
| | L-Valine | 0 | 2 | 1.1 | 0 | 3.1 | 1.9 |
| Metabolite Testing | 2-Oxogluturate | 0 | 0 | 0 | 0 | 0 | 0 |
| | Acetate | 6.7 | 6.2 | 5.3 | 6.6 | 6.7 | 5.5 |
| | Alpha-Aminobutyrate | 0 | 0 | 0 | 0 | 0 | 0 |
| | Beta-Hydroxybutyrate | 7.8 | 7.8 | 6.9 | 7.8 | 7.7 | 6.5 |
| | Butyrate | 8 | 7.1 | 6.2 | 4.3 | 6.6 | 5.4 |
| | Citrate | 0 | 0 | 0 | 0 | 1.5 | 0.3 |
| | D-Malate | 0 | 0.6 | 0 | 0 | 3.8 | 2.6 |
| | D-Mandelate | 0 | 0 | 0 | 0 | 0 | 0 |
| | D-Tartrate | 0 | 0 | 0 | 0 | 0 | 0 |
| | Formate | 0 | 0 | 0 | 0 | 0 | 0 |
| | Fumarate | 6.5 | 6.5 | 5.6 | 6.3 | 3.1 | 1.9 |
| | Gluconate | 3.3 | 6 | 5.1 | 7.8 | 7.7 | 6.5 |
| | D-Glucuronic Acid | 0 | 3.4 | 2.5 | 4.7 | 7.5 | 6.3 |
| | Glutarate | 0 | 0 | 0 | 0 | 0 | 0 |
| | Glyoxylate | 0 | 0 | 0 | 0 | 0 | 0 |
| | L-Malate | 6.4 | 6.2 | 5.3 | 4.2 | 7.2 | 6 |
| | L-Mandelate | 0 | 0 | 0 | 0 | 0 | 0 |
| | L-Tartrate | 0 | 0.9 | 0 | 0 | 2.1 | 0.9 |
| | Malonate | 0 | 0 | 0 | 0 | 0 | 0 |
| | N-Acetyl-Glucosamine | 0 | 1.7 | 0.8 | 0 | 1 | 0 |
| | Oxalate | 0 | 0 | 0 | 0 | 0 | 0 |
| | Propionate | 0 | 0 | 0 | 0 | 0 | 0 |
| | Pyruvate | 0 | 4.9 | 4 | 3.8 | 6.8 | 5.6 |
| | Succinate | 6.8 | 6.5 | 5.6 | 7.1 | 6.6 | 5.4 |
| Other Testing | Benzoate | 0 | 0 | 0 | 0 | 0 | 0 |
| | Glycerol | 0 | 0.8 | 0 | 0 | 0.8 | 0 |
| | Salicine | 0 | 1.1 | 0.2 | 0 | 1.9 | 0.7 |
| | Urea | 0 | 0.1 | 0 | 0 | 0.5 | 0 |
| Sugar Testing | D-Cellobiose | 0 | 1.1 | 0.2 | 0 | 0.8 | 0 |
| | D-Fructose | 0 | 1.5 | 0.6 | 0 | 1.7 | 0.5 |
| | D-Glucose | 0 | 2.4 | 1.5 | 0 | 1.9 | 0.7 |
| | D-Mannose | 0 | 1 | 0.1 | 0 | 1.5 | 0.3 |
| | D-Raffinose | 0 | 1 | 0.1 | 0 | 0.8 | 0 |
| | D-Xylose | 0 | 1.1 | 0.2 | 0 | 1.2 | 0 |
| | D-Galactose | 0 | 0.9 | 0 | 0 | 1 | 0 |
| | myo-Inositol | 0 | 1.3 | 0.4 | 0 | 0.9 | 0 |
| | D-Lactose | 0 | 0.8 | 0 | 0 | 1.6 | 0.4 |
| | L-Arabinose | 0 | 0.8 | 0 | 0 | 2.1 | 0.9 |
| | L-Sorbose | 0 | 2.7 | 1.8 | 0.6 | 2.4 | 1.2 |
| | L-Xylose | 0 | 0.9 | 0 | 0 | 2 | 0.8 |
| | D-Maltose | 0 | 1.2 | 0.3 | 0 | 0.9 | 0 |
| | D-Mannitol | 0 | 1 | 0.1 | 0 | 0.7 | 0 |
| | D-Melibiose | 0 | 1 | 0.1 | 0 | 1.4 | 0.2 |
| | L-Rhamnose | 0 | 1 | 0.1 | 0 | 1.1 | 0 |
| | D-Ribose | 0 | 1.9 | 1 | 1.4 | 2.4 | 1.2 |
| | D-Sorbitol | 0 | 1.2 | 0.3 | 0 | 1 | 0 |
| | Sucrose | 0 | 2.2 | 1.3 | 0 | 0.9 | 0 |
| | Trehalose | 0 | 0.9 | 0 | 0 | 0.6 | 0 |

| Type | Substrate | Rosei | | | | | | | | | | | | | | |
|---|---|---|---|---|---|---|---|---|---|---|---|---|---|---|---|---|
| | | RFI | | | RLI | | | RGL | | | RPP | | | RSY | | |
| | | Substrate Alone | Mixed Substrate and 0.01% Succinate | Mixed Growth Due to Substrate Alone | Substrate Alone | Mixed Substrate and 0.01% Succinate | Mixed Growth Due to Substrate Alone | Substrate Alone | Mixed Substrate and 0.01% Succinate | Mixed Growth Due to Substrate Alone | Substrate Alone | Mixed Substrate and 0.01% Succinate | Mixed Growth Due to Substrate Alone | Substrate Alone | Mixed Substrate and 0.01% Succinate | Mixed Growth Due to Substrate Alone |
| Alcohol Testing | Ethanol | 7.2 | 6.6 | 4.8 | 0.2 | 1.5 | 0 | 0 | 1.7 | 0 | 0.6 | 1.9 | 0 | 0 | 6.6 | 6.6 |
| | Isopropanol | 5.5 | 3.8 | 2 | 3.3 | 1.4 | 0 | 0 | 1.8 | 0.1 | 0 | 1.4 | 0 | 0 | 3 | 3 |
| | Methanol | 3.3 | 3 | 1.2 | 0 | 1.4 | 0 | 0 | 1.7 | 0 | 0 | 2 | 0.1 | 0 | 3.4 | 3.4 |
| | Propanol | 6.3 | 6.3 | 4.5 | 0 | 1.4 | 0 | 0 | 1.4 | 0 | 0 | 0.9 | 0 | 0 | 6 | 6 |
| Amino Acid Testing | L-Alanine | 7 | 6.5 | 4.7 | 0 | 4.5 | 3 | 0 | 3.3 | 1.6 | 0.5 | 0 | 0 | 0 | 6.3 | 6.3 |
| | L-Arginine | 0 | 2.5 | 0.7 | 0 | 1.7 | 0.2 | 0 | 1.4 | 0 | 0 | 0 | 0 | 0 | 2.2 | 2.2 |
| | L-Aspartate | 0 | 0 | 0 | 0 | 0 | 0 | 0 | 0 | 0 | 0 | 0 | 0 | 0 | 0 | 0 |
| | L-Cysteine | 0 | 3.3 | 1.5 | 4.9 | 0.6 | 0 | 4.2 | 3.8 | 2.1 | 0 | 2.4 | 0.5 | 0.8 | 1.7 | 1.7 |
| | L-Glutamine | 6.6 | 6.3 | 4.5 | 0 | 4.8 | 3.3 | 6.5 | 6.6 | 4.9 | 6.8 | 2 | 0.1 | 2.8 | 3.5 | 3.5 |
| | L-Histidine | 0 | 1.7 | 0 | 0 | 0 | 0 | 0 | 2.1 | 0.4 | 0 | 0 | 0 | 0 | 2.9 | 2.9 |
| | L-Isoleucine | 0 | 0 | 0 | 4.3 | 0 | 0 | 2 | 0 | 0 | 2.1 | 0 | 0 | 3.5 | 0 | 0 |
| | L-Leucine | 0 | 0 | 0 | 0.5 | 0 | 0 | 0 | 0 | 0 | 0 | 0 | 0 | 0 | 0 | 0 |
| | L-Lysine | 0.9 | 0 | 0 | 0 | 0 | 0 | 0 | 0 | 0 | 0 | 0 | 0 | 0 | 2.4 | 2.4 |
| | L-Methionine | 0 | 0.8 | 0 | 0 | 0 | 0 | 0 | 0.5 | 0 | 0 | 0 | 0 | 0 | 0.2 | 0.2 |
| | L-Ornithine | 0 | 0 | 0 | 0 | 0 | 0 | 0 | 0 | 0 | 0 | 0 | 0 | 0 | 2.3 | 2.3 |
| | L-Phenylalanine | 0 | 0 | 0 | 1.8 | 0 | 0 | 1.4 | 0 | 0 | 1.8 | 1 | 0 | 0.6 | 0 | 0 |
| | L-Proline | 2.8 | 3 | 1.2 | 0.6 | 3.4 | 1.9 | 0.1 | 1.8 | 0.1 | 0 | 1 | 0 | 0 | 0 | 0 |
| | L-Serine | 2.6 | 2.5 | 0.7 | 0 | 2.6 | 1.1 | 0 | 2.9 | 1.2 | 0 | 0 | 0 | 0 | 0 | 0 |
| | L-Threonine | 0 | 0 | 0 | 0 | 0 | 0 | 0 | 4.1 | 2.4 | 0 | 0 | 0 | 0 | 0 | 0 |
| | L-Tryptophan | 0 | 3.5 | 1.7 | 0 | 1.1 | 0 | 0 | 1.7 | 0 | 0 | 0.9 | 0 | 0 | 0.9 | 0.9 |
| | L-Valine | 0 | 1.6 | 0 | 0 | 0.8 | 0 | 0 | 1.4 | 0 | 0 | 0 | 0 | 0 | 0 | 0 |
| Metabolite Testing | 2-Oxogluturate | 0 | 0 | 0 | 0 | 0 | 0 | 0 | 0 | 0 | 0 | 0 | 0 | 0 | 0 | 0 |
| | Acetate | 0 | 0 | 0 | 4.8 | 4.6 | 3.1 | 0 | 6.4 | 4.7 | 6.5 | 0 | 0 | 0 | 5.7 | 5.7 |
| | Alpha-Aminobutyrate | 0 | 0 | 0 | 0 | 0 | 0 | 0 | 0 | 0 | 0 | 0 | 0 | 0 | 0 | 0 |
| | Beta-Hydroxybutyrate | 7.6 | 7.4 | 5.6 | 7 | 7.4 | 5.9 | 7.5 | 7.9 | 6.2 | 7.6 | 3.6 | 1.7 | 0 | 7.7 | 7.7 |
| | Butyrate | 0 | 0 | 0 | 0 | 5.3 | 3.8 | 0 | 0 | 0 | 0 | 0 | 0 | 0 | 7.8 | 7.8 |
| | Citrate | 0 | 0 | 0 | 0 | 0 | 0 | 2.5 | 0 | 0 | 0 | 0 | 0 | 0 | 0 | 0 |
| | D-Malate | 3.9 | 4.8 | 3 | 0 | 1.5 | 0 | 0 | 4.6 | 2.9 | 0 | 0 | 0 | 3.5 | 0.2 | 0.2 |
| | D-Mandelate | 0 | 0 | 0 | 0 | 0 | 0 | 0 | 0 | 0 | 0 | 0 | 0 | 0 | 0 | 0 |
| | D-Tartrate | 0 | 0 | 0 | 1.9 | 0 | 0 | 2.7 | 0 | 0 | 0 | 0 | 0 | 0 | 0.8 | 0.8 |
| | Formate | 0 | 0 | 0 | 0 | 0 | 0 | 0 | 0 | 0 | 0 | 0 | 0 | 0 | 2.2 | 2.2 |
| | Fumarate | 5.8 | 4.7 | 2.9 | 6.7 | 4.8 | 3.3 | 6.8 | 6.6 | 4.9 | 5.7 | 0 | 0 | 0.1 | 6.7 | 6.7 |
| | Gluconate | 7.8 | 7.6 | 5.8 | 7.8 | 7.8 | 6.3 | 7.8 | 7.7 | 6 | 7.8 | 0 | 0 | 0 | 0 | 0 |
| | D-Glucuronic Acid | 0 | 2.1 | 0.3 | 0 | 1.9 | 0.4 | 0 | 1.8 | 0.1 | 0 | 1.2 | 0 | 0 | 1.2 | 1.2 |
| | Glutarate | 0 | 0 | 0 | 0 | 0 | 0 | 0 | 0 | 0 | 0 | 0 | 0 | 0 | 0 | 0 |
| | Glyoxylate | 0 | 0 | 0 | 0 | 0 | 0 | 0 | 0 | 0 | 0 | 0 | 0 | 0 | 0 | 0 |
| | L-Malate | 5.7 | 4.7 | 2.9 | 6.8 | 4.9 | 3.4 | 6.2 | 7.8 | 6.1 | 5.8 | 0 | 0 | 3.7 | 6.3 | 6.3 |
| | L-Mandelate | 0 | 0 | 0 | 0 | 0 | 0 | 0 | 0 | 0 | 0 | 0 | 0 | 0 | 0 | 0 |
| | L-Tartrate | 2.5 | 3.5 | 1.7 | 0 | 1.6 | 0.1 | 0 | 1.9 | 0.2 | 0 | 0 | 0 | 0 | 1.1 | 1.1 |
| | Malonate | 0 | 0 | 0 | 0 | 0 | 0 | 0.8 | 0 | 0 | 0 | 0 | 0 | 0 | 0 | 0 |
| | N-Acetyl-Glucosamine | 0 | 2.3 | 0.5 | 0 | 1.7 | 0.2 | 0 | 1.7 | 0 | 0 | 0 | 0 | 0 | 0.9 | 0.9 |
| | Oxalate | 0 | 0 | 0 | 0 | 0 | 0 | 0 | 0 | 0 | 0 | 0 | 0 | 0.5 | 0 | 0 |
| | Propionate | 0 | 0 | 0 | 0 | 0 | 0 | 0 | 0 | 0 | 0 | 0 | 0 | 0 | 7.3 | 7.3 |
| | Pyruvate | 6.5 | 6.6 | 4.8 | 6.4 | 6.6 | 5.1 | 6.5 | 7.1 | 5.4 | 5.1 | 0 | 0 | 6.5 | 6 | 6 |
| | Succinate | 6 | 5.6 | 3.8 | 6.8 | 6.4 | 4.9 | 6.6 | 7.2 | 5.5 | 6 | 1.8 | 0 | 6 | 5.5 | 5.5 |
| Other Testing | Benzoate | 0 | 0 | 0 | 0 | 0 | 0 | 0 | 0 | 0 | 0 | 0 | 0 | 0 | 0 | 0 |
| | Glycerol | 0 | 1.9 | 0.1 | 0 | 1.7 | 0.2 | 0 | 2.2 | 0.5 | 0 | 0.6 | 0 | 0 | 0 | 0 |
| | Salicine | 0 | 2.1 | 0.3 | 0 | 1.9 | 0.4 | 0 | 1.9 | 0.2 | 0 | 1.6 | 0 | 0 | 0 | 0 |
| | Urea | 0 | 1.9 | 0.1 | 0 | 1.6 | 0.1 | 0 | 1.5 | 0 | 0 | 0.7 | 0 | 0 | 0 | 0 |
| Sugar Testing | D-Cellobiose | 0 | 1.9 | 0.1 | 0 | 1.5 | 0 | 0 | 1.8 | 0.1 | 0 | 1 | 0 | 0 | 0 | 0 |
| | D-Fructose | 0 | 2.1 | 0.3 | 0 | 2 | 0.5 | 0 | 2 | 0.3 | 0 | 2 | 0.1 | 0 | 0 | 0 |
| | D-Glucose | 0 | 2.2 | 0.4 | 0 | 1.7 | 0.2 | 0 | 2.4 | 0.7 | 0 | 1.3 | 0 | 0 | 0 | 0 |
| | D-Mannose | 0 | 1 | 0 | 0 | 1.2 | 0 | 0 | 0 | 0 | 0 | 0 | 0 | 0 | 1.7 | 1.7 |
| | D-Raffinose | 0 | 1.8 | 0 | 0 | 1.3 | 0 | 0 | 1.3 | 0 | 0 | 0.9 | 0 | 0 | 0 | 0 |
| | D-Xylose | 0 | 2.3 | 0.5 | 0 | 1.8 | 0.3 | 0 | 2 | 0.3 | 0 | 1.6 | 0 | 0 | 0 | 0 |
| | D-Galactose | 0 | 1.7 | 0 | 0 | 1.5 | 0 | 0 | 1.7 | 0 | 0 | 1.3 | 0 | 0 | 0 | 0 |
| | myo-Inositol | 0 | 1.8 | 0 | 0 | 1.4 | 0 | 0 | 1.7 | 0 | 0 | 1.1 | 0 | 0 | 0.5 | 0.5 |
| | D-Lactose | 0 | 1.6 | 0 | 0 | 1.3 | 0 | 0 | 1.6 | 0 | 0 | 0.9 | 0 | 0 | 0 | 0 |
| | L-Arabinose | 0 | 2 | 0.2 | 0 | 1.9 | 0.4 | 0 | 1.7 | 0 | 0 | 1.3 | 0 | 0 | 0 | 0 |
| | L-Sorbose | 0 | 2.3 | 0.5 | 0 | 2.4 | 0.9 | 0 | 2.4 | 0.7 | 1 | 2.2 | 0.3 | 0.9 | 1.7 | 1.7 |
| | L-Xylose | 0 | 2.2 | 0.4 | 0.2 | 2.6 | 1.1 | 0 | 2 | 0.3 | 0 | 1.5 | 0 | 0 | 0 | 0 |
| | D-Maltose | 0 | 1.6 | 0 | 0 | 1.5 | 0 | 0 | 2 | 0.3 | 0 | 1.4 | 0 | 0 | 0 | 0 |
| | D-Mannitol | 0 | 1.5 | 0 | 0 | 1.4 | 0 | 0 | 1.7 | 0 | 0 | 0.9 | 0 | 0 | 0.2 | 0.2 |
| | D-Melibiose | 0 | 1.5 | 0 | 0 | 1.4 | 0 | 0 | 2.1 | 0.4 | 0 | 1 | 0 | 0 | 0 | 0 |
| | L-Rhamnose | 0 | 1.5 | 0 | 0 | 1.3 | 0 | 0 | 1.8 | 0.1 | 0 | 1.6 | 0 | 0 | 0 | 0 |
| | D-Ribose | 1.3 | 3.2 | 1.4 | 0 | 3 | 1.5 | 0 | 2.5 | 0.8 | 2.3 | 2.7 | 0.8 | 0 | 2.1 | 2.1 |
| | D-Sorbitol | 0 | 1.6 | 0 | 0 | 1.5 | 0 | 0 | 1.7 | 0 | 0 | 1.1 | 0 | 0 | 0 | 0 |
| | Sucrose | 0 | 1.9 | 0.1 | 0 | 1.4 | 0 | 0 | 2.6 | 0.9 | 0 | 1.1 | 0 | 0 | 1.1 | 1.1 |
| | Trehalose | 0 | 1.5 | 0 | 0 | 1.2 | 0 | 0 | 1.7 | 0 | 0 | 1.1 | 0 | 0 | 0 | 0 |

**Supplementary Table 14 – A dilution series of a subset of carbon substrates tested indicates some concentration dependent effects on growth in Xanthobacter and Roseixanthobacter gen. nov. type strains.** Substrates were tested in minimal media (4M), at 1% w/v, 0.5% w/v, 0.1% w/v, and 0.05% w/v (and 0.01% w/v for succinate and pyruvate). Numbers indicate the number of doublings from inoculation density ($OD_{600}$ 0.01) that have occurred based on optical density measurements. Background (optical density of each strain in the minimal media without a carbon substrate provided) was subtracted from these doubling values. *Xanthobacter oligotrophicus* (XOG) is represented here by strain 23A instead of type strain 29k$^T$. Abbreviations are: XVE – *Xanthobacter versatilis* Py2$^T$; XAU – *Xanthobacter autotrophicus* 7C$^T$; XOG – *Xanthobacter oligotrophicus* 23A; XTO – *Xanthobacter toluenivorans* T101$^T$; XLU – *Xanthobacter lutulentifluminis* V3C-3$^T$; XNS – *Xanthobacter nonsaccharivorans* 14g$^T$; XAM – Xanthobacter aminoxidans 14a$^T$; XFL – Xanthobacter flavus 301$^T$; XWE – Xanthobacter wiegelii RH 10$^T$; XPS – *Xanthobacter pseudotagetidis* KA$^T$; XTA – *Xanthobacter tagetidis* TagT2C$^T$; XCO – *Xanthobacter cornucopiae* V4C-4$^T$; XAG – *Xanthobacter agilis* SA35$^T$; XSE – *Xanthobacter sediminis* V8C-5$^T$; XAL – *Xanthobacter albus* V0C-6$^T$; XVA – *Xanthobacter variabilis* V4C-8$^T$; RFI – *Roseixanthobacter finlandensis* VTT E-85241$^T$; RLI – *Roseixanthobacter liquoris* VTT E-85238$^T$; RGL – *Roseixanthobacter glucoisosaccharinivorans* VTT E-85242$^T$; RPP – *Roseixanthobacter pseudopolyaromaticivorans* VTT E-85240$^T$; RSY – *Roseixanthobacter psychrophilus* W30$^T$.

| Condition | | Clade I | | | | Clade IIA | | Clade IIB | | | Clade III | | | | | Clade IV | | Rosei | | | | |
|---|---|---|---|---|---|---|---|---|---|---|---|---|---|---|---|---|---|---|---|---|---|---|
| | | XVE | XAU | XOG | XTO | XLU | XNS | XAM | XFL | XWE | XCO | XAG | XSE | XAL | XVA | XPS | XTA | RFI | RLI | RGL | RPP | RSY |
| Succinate | 1% | 4.1 | 4.8 | 3 | 0 | 4.4 | 5.4 | 7 | 7.1 | 7.9 | 7.9 | 4.3 | 5.8 | 6.3 | 5 | 6.9 | 6.8 | 6.2 | 6.1 | 7.4 | 3.6 | 6.7 |
| | 0.50% | 5.2 | 6 | 4.5 | 0 | 5.8 | 5.9 | 7.1 | 6.9 | 7.2 | 7.2 | 4.9 | 6.6 | 6.9 | 6.1 | 7 | 7 | 5.5 | 6.4 | 7 | 5.5 | 6.5 |
| | 0.10% | 4.5 | 5.3 | 4.8 | 0 | 4.8 | 4.9 | 5.3 | 5.1 | 5.4 | 5.7 | 4.8 | 5.2 | 5.4 | 4.8 | 4.9 | 5.2 | 4.9 | 4.6 | 4.2 | 4.9 | 5.7 |
| | 0.05% | 3.9 | 4.9 | 4.4 | 0 | 4.2 | 4 | 4.7 | 4.6 | 4.7 | 4.8 | 4.2 | 4.5 | 4.5 | 3.6 | 4 | 4.2 | 3.9 | 3.4 | 3.2 | 3.5 | 3.4 |
| | 0.01% | 2.6 | 3.1 | 4.6 | 0 | 3.3 | 1.9 | 2.1 | 2.1 | 2.7 | 2.3 | 2 | 2.2 | 2.4 | 1.8 | 0.9 | 1.2 | 1.8 | 1.5 | 1.7 | 1.9 | 0 |
| DL-Lactate | 1% | 4.7 | 5.3 | 1.4 | 0 | 0 | 7.4 | 8.1 | 7.7 | 7.5 | 0 | 0 | 0 | 0 | 0 | 0 | 0 | 0.8 | 0 | 0 | 0 | 0 |
| | 0.50% | 5.8 | 6.9 | 1.8 | 0 | 0 | 7.5 | 7.4 | 7.5 | 7.4 | 0 | 0 | 0 | 0 | 0 | 0 | 0 | 0 | 0 | 0 | 0 | 0 |
| | 0.10% | 5.1 | 5.7 | 4.5 | 0 | 0 | 5.5 | 5.8 | 5.7 | 5.6 | 3.9 | 0 | 0 | 0 | 0 | 0 | 0 | 3.5 | 0 | 0 | 0 | 0 |
| | 0.05% | 4.4 | 5 | 3.7 | 0 | 0 | 4.5 | 4.8 | 4.9 | 4.7 | 3.7 | 0 | 0 | 0 | 0 | 0 | 0 | 3.7 | 0 | 0 | 0 | 0 |
| Glyoxylate | 1% | 0 | 0 | 0 | 0 | 0 | 0 | 0 | 0 | 0 | 0 | 0 | 0 | 0 | 0 | 0 | 0 | 0 | 0 | 0 | 0 | 0 |
| | 0.50% | 0 | 0 | 0 | 0 | 0 | 0 | 0 | 0 | 0 | 0 | 0 | 0 | 0 | 0 | 0 | 0 | 0 | 0 | 0 | 0 | 0 |
| | 0.10% | 4.5 | 2.2 | 4.7 | 1 | 4.5 | 4.3 | 4.3 | 4.6 | 4.4 | 3.2 | 1.8 | 1.8 | 1.3 | 0 | 3.5 | 3.7 | 4 | 4.3 | 4 | 4.2 | 2.5 |
| | 0.05% | 3.8 | 1.4 | 4 | 0.2 | 3.6 | 3.2 | 3.4 | 3.7 | 3.2 | 3.1 | 2.2 | 1.4 | 0.8 | 0 | 1.7 | 2.5 | 2.9 | 3 | 2.4 | 3 | 0 |
| Pyruvate | 1% | 4.9 | 5.4 | 0 | 0 | 6.4 | 6.5 | 7.8 | 7.6 | 5 | 7.8 | 6.7 | 5.7 | 6.9 | 3.3 | 0.6 | 3.5 | 7.3 | 6.1 | 8.4 | 0 | 5.3 |
| | 0.50% | 6.2 | 6.6 | 6.2 | 2.3 | 6.8 | 7 | 7.6 | 7 | 7.4 | 7.3 | 6.5 | 6.7 | 7.1 | 5.1 | 0.1 | 2.4 | 7 | 7 | 7.2 | 0 | 6.5 |
| | 0.10% | 5.2 | 4.7 | 5.8 | 2.2 | 5.2 | 5 | 5.1 | 5.1 | 5.2 | 5.6 | 5.4 | 5.4 | 5.6 | 4.5 | 2.9 | 1.5 | 5.2 | 5.3 | 5.1 | 2.1 | 5.3 |
| | 0.05% | 4.4 | 3.4 | 4.9 | 0 | 4.3 | 4 | 4.4 | 4.2 | 4.4 | 4.6 | 4.7 | 4.7 | 4.6 | 3.7 | 3.4 | 2.5 | 4.1 | 4.3 | 4.1 | 1.8 | 4.3 |
| | 0.01% | N/A | N/A | N/A | 0 | N/A | N/A | N/A | N/A | N/A | N/A | N/A | N/A | N/A | N/A | N/A | N/A | N/A | N/A | N/A | N/A | N/A |
| Ribose | 1% | 1.9 | 2.8 | 0.8 | 0 | 0.8 | 1.5 | 2.7 | 2 | 1.7 | 0.9 | 0 | 0 | 0 | 0 | 0 | 2.3 | 2.8 | N/A | 0 | 2.4 | 0 |
| | 0.50% | 0.3 | 1.6 | 0 | 0 | 0 | 0 | 2 | 1.4 | 0.8 | 0 | 0 | 0 | 0 | 0.6 | 0 | 0.3 | 2.4 | N/A | 0 | 1.6 | 0 |
| | 0.10% | 0 | 0 | 0 | 0 | 0 | 0 | 0.4 | 0.7 | 0.7 | 1.5 | 0 | 0 | 0 | 0 | 0 | 0 | 0 | N/A | 0 | 0 | 0 |
| | 0.05% | 0 | 0 | 0 | 0 | 0 | 0 | 0 | 0 | 0 | 0.8 | 0.6 | 0 | 0 | 0 | 0 | 0 | 0 | N/A | 0 | 0 | 0 |
| Fructose | 1% | 8.4 | 8.5 | 8.3 | 0 | 0 | 0 | 0 | 0.9 | 0 | 0 | 0 | 0 | 0 | 0 | 0 | 0.1 | 0.4 | N/A | 0 | 0.6 | 0 |
| | 0.50% | 7.9 | 7.9 | 7.7 | 0 | 0 | 0 | 0 | 0.4 | 0 | 0 | 0 | 0 | 0 | 0 | 0 | 0 | 0.3 | N/A | 0 | 0 | 0 |
| | 0.10% | 5.4 | 6.1 | 5.2 | 0 | 0 | 0 | 0 | 0.6 | 0.1 | 0 | 0 | 0 | 0 | 0 | 0 | 0 | 0 | N/A | 0 | 0 | 0 |
| | 0.05% | 4.7 | 5.2 | 3.7 | 0 | 0 | 0 | 0 | 0 | 0.3 | 0 | 0 | 0 | 0 | 0 | 0 | 0 | 0 | N/A | 0 | 0 | 0 |
| Glucose | 1% | 8.4 | 0 | 1.2 | 0 | 0 | 0.1 | 0 | 1.5 | 1.7 | 0 | 0 | 0 | 0 | 0 | 0 | 0 | 0 | N/A | 0 | 0 | 0 |
| | 0.50% | 8.4 | 0 | 0 | 0 | 0 | 0 | 0.2 | 0 | 2.8 | 0 | 0 | 0 | 0 | 0 | 0 | 0 | 0 | N/A | 0 | 0 | 0 |
| | 0.10% | 6.5 | 0 | 0 | 0 | 0 | 0.3 | 0 | 0.5 | 0 | 0 | 0 | 0 | 0 | 0 | 0 | 0 | 0 | N/A | 0 | 0 | 0 |
| | 0.05% | 5.4 | 0 | 0 | 0 | 0 | 0 | 0 | 0 | 0.6 | 0 | 0 | 0.1 | 0 | 0 | 0 | 0 | 0 | N/A | 0 | 0 | 0 |
| Sucrose | 1% | 0 | 0.9 | 0.9 | 0 | 0 | 0 | 0 | 0 | 0.4 | 0 | 0 | 0 | 1.2 | 0 | 0 | 0 | 0 | N/A | 0 | 0 | 0 |
| | 0.50% | 0 | 0 | 0 | 0 | 0 | 0 | 0 | 0 | 0.1 | 0 | 0 | 0 | 0 | 0 | 0 | 0 | 0 | N/A | 0 | 0 | 0 |
| | 0.10% | 0 | 0 | 0 | 0 | 0 | 0 | 0 | 0 | 0 | 0 | 0 | 0 | 0 | 0 | 0 | 0 | 0 | N/A | 0 | 0 | 0 |
| | 0.05% | 0 | 0 | 0 | 0 | 0.4 | 0 | 0 | 0 | 0 | 0 | 0 | 0 | 0 | 0 | 0 | 0 | 0 | N/A | 0 | 0 | 0 |

**Supplementary Table 14 – Enzymatic activities are largely shared across the Xanthobacter and Roseixanthobacter gen. nov. type strains in the API ZYM enzymatic panel.** A positive result is indicated with a “+”, a weakly positive result is indicated with a “W”. All assays were carried out according to manufacturer instructions (observed after 4.5 hours). *Xanthobacter oligotrophicus* (XOG) is represented here by strain 23A instead of type strain 29k$^T$. Abbreviations are: XVE – *Xanthobacter versatilis* Py2$^T$; XAU – *Xanthobacter autotrophicus* 7C$^T$; XOG – *Xanthobacter oligotrophicus* 23A; XTO – *Xanthobacter toluenivorans* T101$^T$; XLU – *Xanthobacter lutulentifluminis* V3C-3$^T$; XNS – *Xanthobacter nonsaccharivorans* 14g$^T$; XAM – Xanthobacter aminoxidans 14a$^T$; XFL – Xanthobacter flavus 301$^T$; XWE – Xanthobacter wiegelii RH 10$^T$; XPS – *Xanthobacter pseudotagetidis* KA$^T$; XTA – *Xanthobacter tagetidis* TagT2C$^T$; XCO – *Xanthobacter cornucopiae* V4C-4$^T$; XAG – *Xanthobacter agilis* SA35$^T$; XSE – *Xanthobacter sediminis* V8C-5$^T$; XAL – *Xanthobacter albus* V0C-6$^T$; XVA – *Xanthobacter variabilis* V4C-8$^T$; RFI – *Roseixanthobacter finlandensis* VTT E-85241$^T$; RLI – *Roseixanthobacter liquoris* VTT E-85238$^T$; RGL – *Roseixanthobacter glucoisosaccharinivorans* VTT E-85242$^T$; RPP – *Roseixanthobacter pseudopolyaromaticivorans* VTT E-85240$^T$; RSY – *Roseixanthobacter psychrophilus* W30$^T$.

| Test | Species | | | | | | | | | | | | | | | | | | | |
|---|---|---|---|---|---|---|---|---|---|---|---|---|---|---|---|---|---|---|---|---|
| | **Clade I** | | | | **Clade IIA** | | **Clade IIB** | | | **Clade III** | | | | | **Clade IV** | | **Roseixanthobacter** | | | | |
| | **XVE** | **XAU** | **XOG*** | **XTO** | **XLU** | **XNS** | **XAM** | **XFL** | **XWE** | **XCO** | **XAG** | **XSE** | **XAL** | **XVA** | **XPS** | **XTA** | **RFI** | **RLI** | **RGL** | **RPP** | **RSY** |
| Alkaline phosphatase | W | W | W | W | + | W | W | W | | W | + | W | W | + | W | W | + | + | + | + | W |
| Acid phosphatase | + | + | + | + | + | + | + | + | + | + | + | + | + | + | + | + | + | + | + | + | + |
| Esterase (C 4) | + | + | + | + | + | + | + | + | + | + | + | + | + | + | + | + | + | + | + | + | + |
| Esterase Lipase (C 8) | + | + | + | + | + | + | + | + | + | + | + | + | + | + | + | + | + | + | + | + | + |
| Lipase (C 14) | | | | | | | | | | | | | | | | | | | | | |
| Leucine arylamidase | | | | | | | | | | | | | | | | | | | | | |
| Valine arylamidase | W | W | W | W | W | W | W | W | W | W | W | W | W | W | W | W | W | W | W | W | W |
| Cysteine arylamidase | W | W | W | W | W | | | | | | | | | W | | | W | W | W | W | W |
| Trypsin | + | + | + | + | + | + | + | + | + | + | + | + | + | + | + | + | + | + | + | + | + |
| α-chymotrypsin | | | | | | | | | | | | | | | | | | | | | |
| Napthol-AS-BI-phospho-hydrolase | + | + | + | + | + | + | + | + | + | + | + | + | + | + | + | + | + | + | + | + | + |
| α-galactosidase | | | | | | | | | | | | | | | | | | | | | |
| β-galactosidase | | | | | | | | | | | | | | | | | | | | | |
| β-glucuronidase | | | | | | | | | | | | | | | | | | | | | |
| α-glucosidase | | | | | | | | | | | | | | | | | | | | | |
| β-glucosidase | | | | | | | | | | | | | | | | | | | | | |
| N-acetyl-β-gluco-saminidase | | | | | | | | | | | | | | | | | | | | | |
| α-mannosidase | | | | | | | | | | | | | | | | | | | | | |
| α-fucosidase | | | | | | | | | | | | | | | | | | | | | |

**Supplementary Table 15 – Enzymatic and metabolic activities follow loosely along evolutionarily related clades in Xanthobacter and Roseixanthobacter gen. nov. type strains in the API 20NE enzymatic and metabolic panel.** Readout occurred according to manufacturer instructions with the nitrate reduction and indole tests at 24 hours and the rest at 48-72 hours. A positive result is indicated with a "+", a weakly positive result is indicated with a "W". Due to the slow growth of Xanthobacter and Roseixanthobacter gen. nov. strains, assay strips were incubated for 7 days in total. Positive readouts after 5 days are indicated with a dagger "†" and after 7 days with a double dagger "‡". Cytochrome oxidase activity was determined using Millipore Sigma Oxidase test strips. *Xanthobacter oligotrophicus* (XOG) is represented here by strain 23A instead of type strain 29k$^{T}$. Abbreviations are: XVE – *Xanthobacter versatilis* Py2$^{T}$; XAU – *Xanthobacter autotrophicus* 7C$^{T}$; XOG – *Xanthobacter oligotrophicus* 23A; XTO – *Xanthobacter toluenivorans* T101$^{T}$; XLU – *Xanthobacter lutulentifluminis* V3C-3$^{T}$; XNS – *Xanthobacter nonsaccharivorans* 14g$^{T}$; XAM – Xanthobacter aminoxidans 14a$^{T}$; XFL – Xanthobacter flavus 301$^{T}$; XWE – Xanthobacter wiegelii RH 10$^{T}$; XPS – *Xanthobacter pseudotagetidis* KA$^{T}$; XTA – *Xanthobacter tagetidis* TagT2C$^{T}$; XCO – *Xanthobacter cornucopiae* V4C-4$^{T}$; XAG – *Xanthobacter agilis* SA35$^{T}$; XSE – *Xanthobacter sediminis* V8C-5$^{T}$; XAL – *Xanthobacter albus* V0C-6$^{T}$; XVA – *Xanthobacter variabilis* V4C-8$^{T}$; RFI – *Roseixanthobacter finlandensis* VTT E-85241$^{T}$; RLI – *Roseixanthobacter liquoris* VTT E-85238$^{T}$; RGL – *Roseixanthobacter glucoisosaccharinivorans* VTT E-85242$^{T}$; RPP – *Roseixanthobacter pseudopolyaromaticivorans* VTT E-85240$^{T}$; RSY – *Roseixanthobacter psychrophilus* W30$^{T}$.

| Test | Species | | | | | | | | | | | | | | | | | | | | |
|---|---|---|---|---|---|---|---|---|---|---|---|---|---|---|---|---|---|---|---|---|---|
| | Clade I | | | | Clade IIA | | Clade IIB | | | Clade III | | | | | Clade IV | | Roseixanthobacter | | | | |
| | XVE | XAU | XOG* | XTO | XLU | XNS | XAM | XFL | XWE | XCO | XAG | XSE | XAL | XVA | XPS | XTA | RFI | RLI | RGL | RPP | RSY |
| Reduction of nitrates to nitrites | W | + | + | | + | + | + | + | + | + | + | + | + | + | | | | | | | + |
| Reduction of nitrates to N2 | | | | | | | | | | | | | | | | | | | | | |
| Indole production | | | | | | | | | | | | | | | | | | | | | |
| Glucose fermentation | | | | | | | | | | | | | | | | | | | | | |
| Arginine dihydrolase | | | | | | | | | | | | | | | ‡ | | ‡ | | ‡ | | |
| Urease | W | W | | | w ‡ | | W | | w ‡ | W | | W | + | W | + | + | W | W | W | W | W |
| β-glucosidase hydrolysis | | | | | | | | | | | | | | | | | | | | | |
| Protease hydrolysis (gelatin) | | | | | | | | | | + | | | | + | | | | | | | |
| β-galactosidase | | | | | | | | | | | | | | | | | | | | | |
| Glucose assimilation | † | | | | | | | | | | | | | | | | | | | | |
| Arabinose assimilation | | | | | | | | | | | | | | | | | | | | | |
| Mannose assimilation | | | | | | | ‡ | | | | | | | | | | | | | | |
| Mannitol assimilation | | | | | | | | | | | | | | | | | | | | | |
| N-Acetyl-Glucosamine assimilation | | | | | | | | | | | | | | | | | | | | | |
| Gluconate assimilation | † | † | ‡ | | † | ‡ | † | † | † | | | | | | | ‡ | † | † | † | † | |
| Capric acid assimilation | | | | | | | | | | | | | | | | | | | | | |
| Adipic acid assimilation | | ‡ | | | ‡ | † | | † | † | † | ‡ | ‡ | ‡ | † | | | | | | | |
| Malate assimilation | | ‡ | | | | ‡ | † | † | † | † | † | † | † | † | † | † | † | † | † | † | ‡ |
| Citrate assimilation | | | | | | | † | | ‡ | | | | | | | | | | | | |
| Phenylacetic acid assimilation | | | | | | | | | | | | | | | | | | | | | |
| Cytochrome oxidase | + | + | + | + | + | + | + | + | + | + | + | + | + | + | + | + | + | + | + | + | + |
